# Increasing quantitation in spatial single-cell metabolomics by using fluorescence as ground truth

**DOI:** 10.1101/2022.08.24.505109

**Authors:** Martijn R. Molenaar, Mohammed Shahraz, Jeany Delafiori, Andreas Eisenbarth, Måns Ekelöf, Luca Rappez, Theodore Alexandrov

## Abstract

Imaging mass spectrometry (MS) is becoming increasingly applied for single-cell analyses. Multiple methods for imaging MS-based single-cell metabolomics were proposed, including our recent method SpaceM. An important step in imaging MS-based single-cell metabolomics is the assignment of MS intensities from individual pixels to single cells. In this process, referred to as pixel-cell deconvolution, the MS intensities of regions sampled by the imaging MS laser are assigned to the segmented single cells. The complexity of the contributions from multiple cells and the background, as well as lack of full understanding of how input from molecularly-heterogeneous areas translates into mass spectrometry intensities make the cell-pixel deconvolution a challenging problem.

Here, we propose a novel approach to evaluate pixel-cell deconvolution methods by using a molecule detectable both by mass spectrometry and fluorescent microscopy, namely fluorescein diacetate (FDA). FDA is a cell-permeable small molecule that becomes fluorescent after internalisation in the cell and subsequent cleavage of the acetate groups. Intracellular fluorescein can be easily imaged using fluorescence microscopy. Additionally, it is detectable by matrix-assisted laser desorption/ionisation (MALDI) imaging MS. The key idea of our approach is to use the fluorescent levels of fluorescein as the ground truth to evaluate the impact of using various pixel-cell deconvolution methods onto single-cell fluorescein intensities obtained by the SpaceM method.

Following this approach, we evaluated multiple pixel-cell deconvolution methods, the ‘weighted average’ method originally proposed in the SpaceM method as well as the novel ‘linear inverse modelling’ method. Despite the potential of the latter method in resolving contributions from individual cells, this method was outperformed by the weighted average approach. Using the ground truth approach, we demonstrate the extent of the ion suppression effect which considerably worsens the pixel-cell deconvolution quality. For compensating the ion suppression, we propose a novel data-driven approach. We show that compensating the ion suppression effect in a single-cell metabolomics dataset of co-cultured HeLa and NIH3T3 cells considerably improved the separation between both cell types. Finally, using the same ground truth, we evaluate the impact of drop-outs in the measurements and discuss the optimal filtering parameters of SpaceM processing steps before pixel-cell deconvolution.

## Introduction

Imaging mass spectrometry is becoming an increasingly popular technology for single-cell metabolomics (Rubakhin, Lanni and Sweedler, 2013; Zenobi, 2013; Liu and Yang, 2021; Taylor, Lukowski and Anderton, 2021). We recently developed SpaceM, an open-source method to perform spatial single-cell metabolomics that integrates microscopy and imaging mass spectrometry (imaging MS) (Figure 1A) (Rappez *et al.*, 2021).

**Figure 1.**
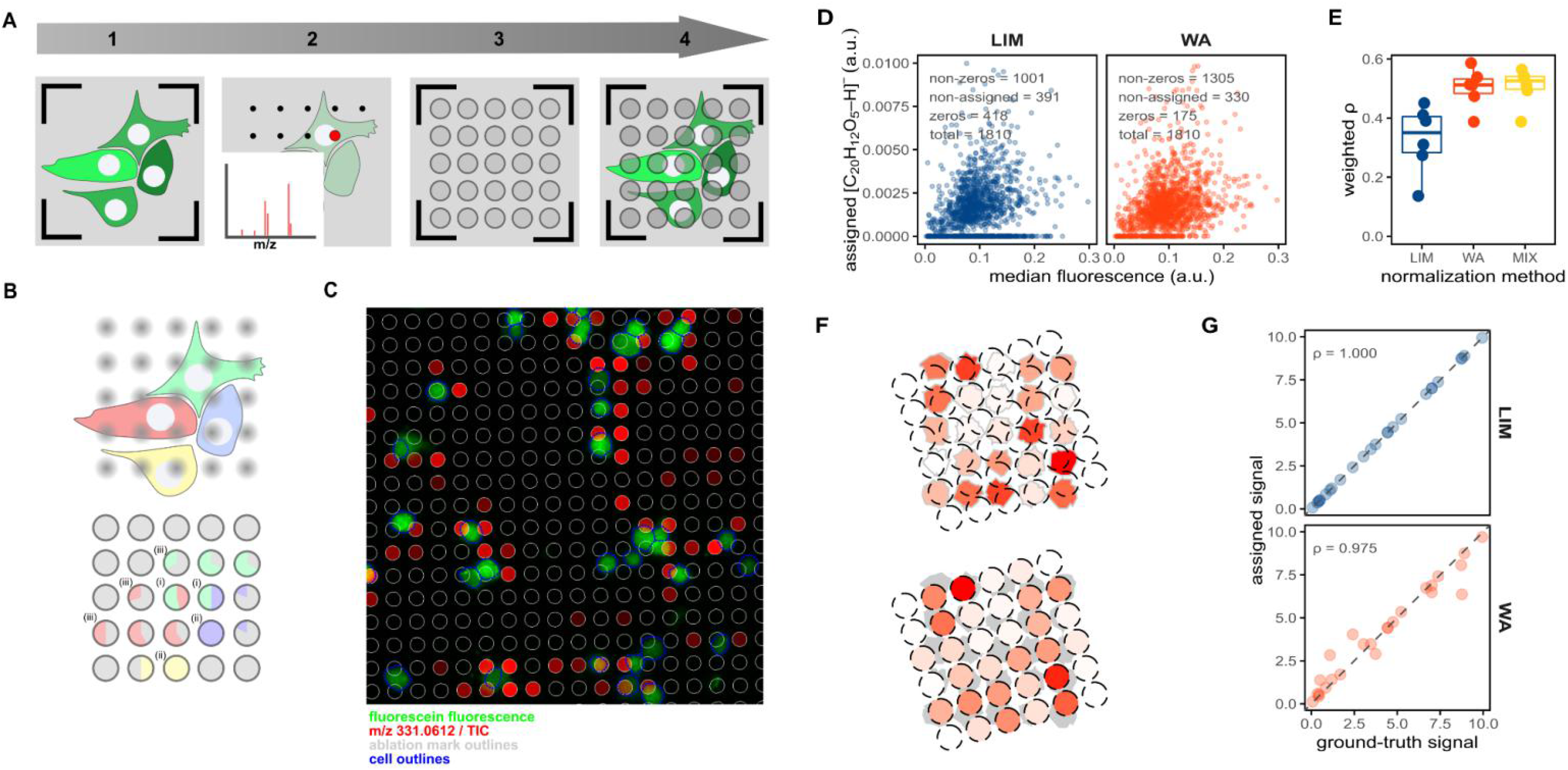
Evaluating pixel-cell deconvolution in imaging MS-based spatial single-cell metabolomics by using fluorescein. **A.** Procedure of SpaceM prior to pixel-cell deconvolution: (1) pre-MALDI microscopy, (2) MALDI-imaging MS acquisition, (3) post-MALDI microscopy, (4) image registration, ablated region selection and cell segmentation. **B.** Pixel-cell deconvolution and the confounding factors. Single-cell pie charts illustrate the contributions of different cells or the extracellular area to the ablated regions sampling (i) multiple cells, (ii) a single cell, or (iii) both intra- and extracellular areas. Pixel-cell deconvolution methods aim to estimate the metabolite levels of the single cells depicted in red, green, blue and yellow. **C.** Overlay of the fluorescein fluorescence (green), fluorescein MALDI-signals (red), laser-ablated regions (grey) and segmented cells (blue). **D.** Representative scatter plots (belonging to one replicate) of median fluorescein fluorescence versus assigned fluorescein MALDI-signal of single cells for the tested pixel-cell deconvolution methods: linear inverse modelling (LIM), weighted average (WA). The methods use sampling proportion cut-offs of 0.3, mass spectral intensities are integrated within 4 ppm m/z tolerance and TIC-normalised. **E.** Boxplot with density-weighted Spearman correlations between fluorescent and MALDI intensities of fluorescein obtained for each of the pixel-cell deconvolution methods WA, LIM and the mixed model (MIX) for 6 replicates. **F.** Illustration of the process of data simulation to evaluate the pixel-cell deconvolution methods. Cell signals are colour-scaled from white to red for cells (ground truth, top panel) and ablated regions (measured intensities, bottom panel). G. Scatter plots with Spearman correlations of the ground truth *versus*assigned signals in the simulated data when using the pixel-cell deconvolution methods LIM (blue) or WA (red).

In SpaceM, cellular metabolite intensities are calculated from imaging mass spectra of pixels overlapping with segmented cells in the process hereinafter called as pixel-cell deconvolution (previously referred to as normalisation (Rappez *et al.*, 2021)). As the regions sampled by the imaging MS laser (hereinafter, ablated regions) do not always completely overlap with the cellular regions or overlap with multiple cells, the pixel-cell deconvolution is a challenging task (Figure 1B). Moreover, there is a lack of full understanding of how co-ablated regions from multiple cells or from cells and background contribute to integrated intensities in a mass spectrum due to the ion suppression, namely when intensities of one molecular species are reduced by the presence of another abundant and easily-ionizable molecular species. Earlier, we proposed a pixel-cell deconvolution method called ‘weighted average’ that involves two steps (Rappez *et al.*, 2021). First, the MS signals of the ablated regions are normalised by dividing them by their ‘sampling proportion’, which is the fraction of their overlap with cellular regions. Second, the weighted average of all normalised ablated regions associated with a cell-of-interest is calculated. In this approach, the weights are represented by the ‘sampling specificities’: the fractions in which the cellular sampling areas are overlapping with the cell-of-interest. Finally, the resulting weighted average is assigned to the cell-of-interest as intensity, after which the procedure is repeated for all other cells.

The motivation for the weighted average method was to reduce the impact of ablated regions associated with multiple cells (hereinafter, co-ablated regions). To a certain extent, however, the co-ablated regions do still mix signals from two neighbouring cells. To this end, we additionally proposed to filter out the most extreme co-ablated regions by using a cut-off for sampling specificity (Rappez *et al.*, 2021). Furthermore, we used cut-offs for sampling proportion to reduce the effect of ablated regions with limited overlap with cells. However, stringent cut-offs may lead to a loss of data points and possibly even to a failure in assigning signals to the cell-of-interest.

Here, we propose a novel method for pixel-cell deconvolution, relying on linear inverse modelling. We propose to model the metabolite signals associated with the ablated regions (which are the knowns) as a linear combination of the cellular metabolite signals (the unknowns) multiplied by their ‘specific sampling proportion’, which is the sampling proportion multiplied by the sampling specificity of the ablated-region-of-interest (the knowns). The resulting system of linear equations is overdetermined, as the number of ablated regions is larger than the number of cells by experimental design (hence, pixel-cell deconvolution would not be possible otherwise) and can be approached and solved as a linear inverse problem.

Both methods, the weighted average (‘WA’) and linear inverse modelling (‘LIM’), assume the linearity of the signals. However, this assumption is not always satisfied in mass spectrometry. A key factor compromising the linearity is the ion suppression generally present in MS (Annesley, 2003; Furey *et al.*, 2013). In this process, the detector response for a particular molecular species in the analysed complex mixture is reduced due to the competition between various molecular species to obtain charge. As molecular species have different ionisation efficiencies, the magnitude of the response reduction for a given molecular species depends on other molecules present in the analysed sample, in particular on the abundant ones. Ion suppression is a considerable factor in imaging MS, which is performed without prior chromatography, *i.e.* all molecular species are analysed simultaneously. In imaging MS-based single-cell metabolomics, ion suppression may lead to decreased signals in cellular regions containing highly-abundant analytes as compared to regions outside of cells. Moreover, the MALDI matrix signals can have higher intensities in the extracellular regions even if their concentration is the same throughout the full analysed area. Thus, metabolite intensities from ablated regions non- or partially-overlapping with cells are expected to be overestimated, thereby introducing a potential error into the cell pixel-cell deconvolution. However, there is currently no method for quantitative estimation and compensation of the impact of ion suppression onto single-cell intensities, partially due to the lack of known metabolite concentrations in single cells.

Here, we aim to bridge this gap by proposing a novel approach to estimate the quantitation in spatial single-cell metabolomics by using a fluorescent dye, and demonstrate how it can be used to evaluate pixel-cell deconvolution methods. The key idea of the approach is to use a fluorescent dye that can be detected by both fluorescent microscopy and imaging MS. We chose fluorescein diacetate (FDA), a compound widely used in cell biology to assess cell viability (Mckinney, Spillane and Pearce, 1964; Jones and Senft, 1985), because it has high fluorescence, high mass spectrometry response, has no reported metabolic effects, and is fluorescent only when intracellular. In its esterified form, the compound is cell-permeable and not fluorescent. After internalisation in the cell, the acetate groups are cleaved by cellular esterases, and free fluorescein is formed (Rotman and Papermaster, 1966). Intracellular fluorescein can be easily imaged using widely available fluorescence filters used for green fluorescent protein. Additionally, fluorescein is detectable by matrix-assisted laser desorption/ionisation (MALDI) imaging MS (Liu *et al.*, 2013) used in SpaceM and other approaches for single-cell metabolomics. As a result, the fluorescent levels of fluorescein can serve as the ground truth and can be compared with MS-derived intensities of fluorescein obtained by a particular pixel-cell deconvolution strategy.

Following this approach, we evaluated two pixel-cell deconvolution approaches for SpaceM, as well as optimised their parameters. We observed that, although the LIM method is expected to perform superior by theoretical considerations, it is outperformed by the WA method. Furthermore, we show that non-linearity in MS measurements due to effects such as ion suppression considerably worsens the quantitation and propose a method to compensate for this effect.

## Results

### Experimental ground-truth model with multi-modal fluorescent and MALDI-MS readout

To assess the pixel-cell deconvolution quality of single-cell metabolomics obtained by MALDI-imaging MS, we sought for a fluorescent probe that is retained in cells, has little cytotoxicity or effect on metabolism, is chemically inert and can be detected by MALDI-MS. After testing several candidates, we selected FDA as the most promising dye. To evaluate pixel-cell deconvolution methods, we aimed to maximise the dynamic range of fluorescein inside the cells. Therefore, we incubated adherent HeLa cells with four different concentrations of FDA (see Methods). After incubation, we trypsinized, pooled the cells together and immobilised them on the cell culture surface of an 8-well chamber slide by centrifugation. Subsequently, we performed spatial single-cell metabolomics using the SpaceM method as described previously (Rappez *et al.*, 2021).

After registration of pre- and post-MALDI microscopy images based on the fiducial pen marks (Figure 1A), we were able to overlay the fluorescence of fluorescein with the ion image of m/z 331.0612, corresponding to the [M-H]^−^ ion of fluorescein (Figure 1C). As expected, we observed higher fluorescein signals in cellular areas and a good agreement between the two modalities.

### Impact of pixel-cell deconvolution methods on the quantitation

We then segmented the cells using the brightfield channel of the pre-MALDI microscopy image, applied a grid-fitting approach with fixed-radius circular shapes to estimate the ablated regions and calculated the sampling proportions, sampling specificities, and specific sampling proportions (Figure S2) to be used as inputs for the two pixel-cell deconvolution methods WA and LIM. After using the optimal pre-processing settings of the input measurements (as discussed in the next sections), we applied the two pixel-cell deconvolution methods. We then calculated weighted Spearman correlations between both fluorescent and MALDI modalities as a measure of pixel-cell deconvolution quality (Figure 1D). For each method, a scatter plot showed the expected relation between microscopy and MS signal. Surprisingly, the total number of cells with non-zero assignments was higher in the WA method. Furthermore, the weighted Spearman correlation coefficients of the WA method were considerably higher (median ρ of ca. 0.50) than for the LIM method (median ρ of ca. 0.35) (Figure 1E). This is surprising, as the ablated regions co-sampling multiple cells should represent a linear combination of the ions coming from the sampled cells, especially considering the frequent occurrence of co-ablated regions in our data (Figure S1). Indeed, applying both methods on simulated data lacking any measuring error (Figure 1F) results in a perfect correlation between the ground-truth and calculated values for the LIM method, whereas WA cannot fully recapitulate the ground-truth signals (Figure 1G).

We speculated that the LIM method might suffer from local underdetermined systems (networks of cells that are connected by overlapping ablated regions, with more cells than ablated regions present). Indeed, we found that about 12% of the cells were members of a local underdetermined system. We tested whether a mixed method could further improve pixel-cell deconvolution. To this end, we first applied LIM, after which we reapplied WA on the underdetermined cells. However, this approach improved the pixel-cell deconvolution only to a little extent (Figure 1E, ‘MIX’).

### Investigating ion suppression in single-cell metabolomics

A major factor affecting quantitation in MS and in particular in imaging MS is ion suppression. We have evaluated whether mass spectrometry intensities of fluorescein are reduced by the ion suppression in the cells compared to the background. First, for each ablated region we defined the ratio *η* as its fluorescein [M-H]^−^intensity versus the corresponding fluorescence intensity. Next, we assumed that for an ablated region with a high sampling proportion ("cellular" ablated region), the MS intensity (and consequently the ratio *η*) is reduced compared to an ablated region with a low sampling proportion (“background” ablated region).

To test this, we plotted the ratio *η* as a function of the ablated region sampling proportion (Figure 2A). Indeed, we observed a decrease of *η* with the increase of the sampling proportion, following a negative power-law relationship (linear after log-transformation of both axes), which is in line with our hypothesis of the non-linear signal reduction due to the ion suppression.

**Figure 2.**
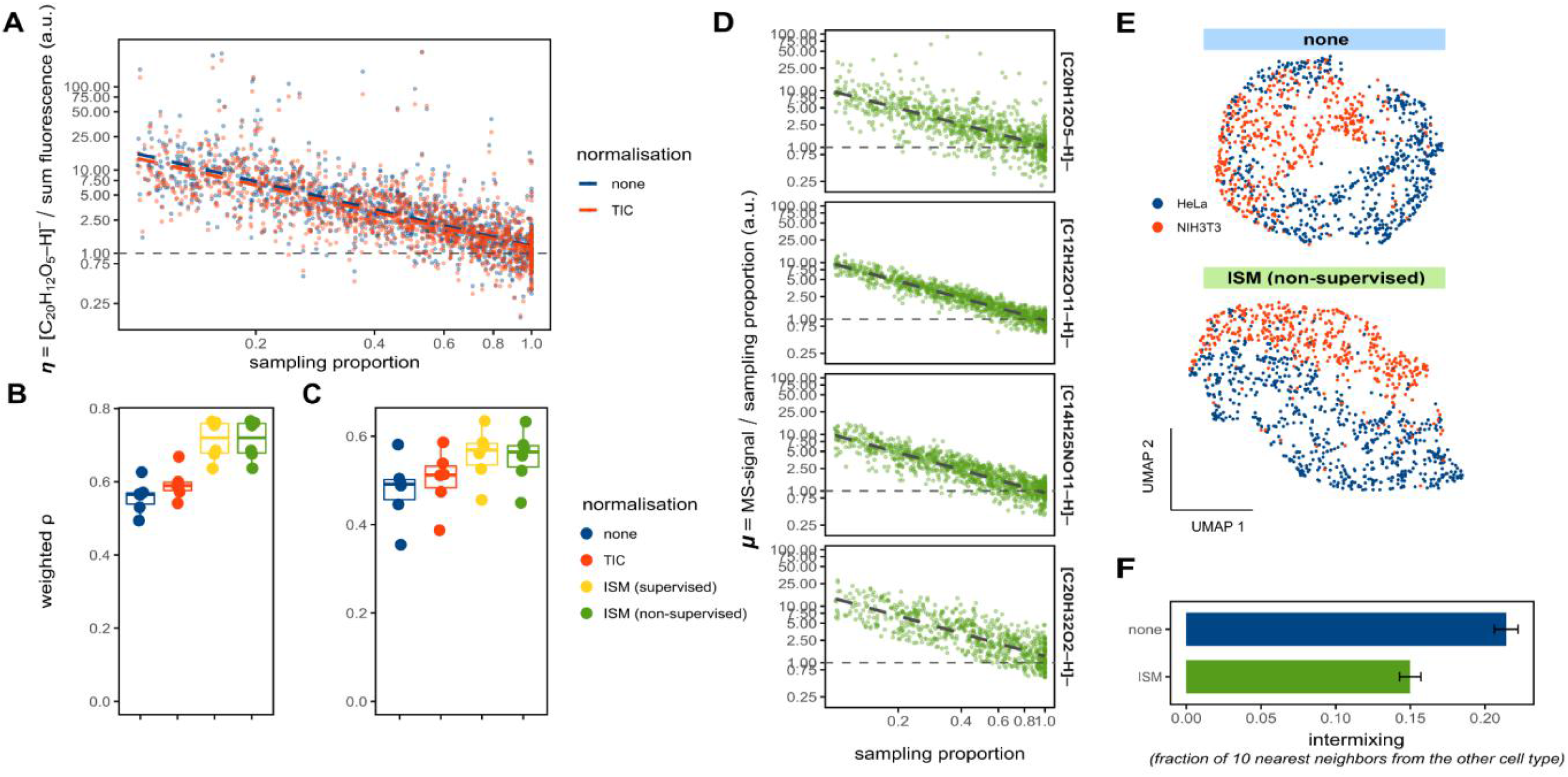
Compensating for ion suppression in spatial single-cell metabolomics. **A.** Representative scatter plot for ablated regions (belonging to one replicate), plotting the scaled ratio *η* (fluorescein ion [M-H]^−^ intensity divided by the fluorescence) against the ablated region sampling proportion; the ratios *η*are set to 1 for sampling proportions of 1; using TIC-normalised (red) and unnormalized (blue) MS intensities; both axes are log-transformed. **B-C**. Boxplots with density-weighted Spearman correlations between fluorescence and MS intensities of fluorescein for either ablated regions **(B)** or cells **(C)** using the WA method (sampling proportion of 0.3, m/z tolerance 4 ppm) for 6 replicates,with different normalisation methods including the supervised and unsupervised ion suppression method (ISM). D.Representative scatter plots of the ratio *μ*plotted against the sampling proportion for four ions ([C_20_ H_12_O_5_-H]^−^ corresponding to fluorescein, [C_12_H_22_O_11_-H]^−^ for a disaccharide, [C_14_H_25_NO_11_-H]^−^ for a polysaccharide, and [C_20_H_32_O_2_-H]^−^ for arachidonic acid); ratios are set to 1 for sampling proportions of 1; both axes are log-transformed. E. UMAP visualisation of single-cell metabolomics data obtained from a co-culture of HeLa (blue) and NIH3T3 (red) cells (Rappez *etal.*, 2021). Cells were WA-normalised with (upper panel) or without (bottom panel) applying the unsupervised method for compensating the ion suppression (ISM). F. Quantification of the intermixing between the cell types from panel E, with barplots showing the mean and standard deviation of the mean fractions of all cell’s 10 nearest-neighbours with the opposite cell type, per normalisation approach.

In untargeted metabolomics, modelling ion suppression is challenging as neither the molecular composition of the analysed material nor the relative ionisation efficiencies of the analytes are known. Nevertheless, approaches such as total-ion current (TIC)-normalisation have been suggested to decrease the impact of ion suppression in imaging MS (Taylor, Dexter and Bunch, 2018). Indeed, applying the TIC-normalisation minimised MS intensity overestimation of the ablated regions with lower sampling proportions, as revealed by a slightly lower slope (Figure 2A). Furthermore, the correlation between fluorescence and MS intensities for both ablated regions (Figure 2B) and cells (Figure 2C) improved after TIC-normalisation. To further investigate the impact of ion suppression on pixel-cell deconvolution, we estimated η of each ablated region by quantile regression of the log-transformed data points of Figure 2A. After dividing the fluorescein [M-H]- intensities by the estimated η (thus, compensating for ion suppression), the correlations between fluorescence and MS intensities for both ablated regions (Figure 2B) and cells (Figure 2C) improved considerably. This demonstrates the impact of ion suppression on single-cell intensities.

### Compensating ion suppression in spatial single-cell metabolomics

Estimating *η* to compensate for the ion suppression, however, cannot be done for endogenous molecules, since it requires corresponding fluorescent readouts. Therefore, we developed an unsupervised method for compensating for the ion suppression. First, we assumed that MS-intensities for the ablated regions are proportional to their sampling proportions in case of no ion suppression. Second, for each ablated region we considered the ratio *μ* between the MS-intensity of fluorescein [M-H]^−^and its sampling proportion. When we plotted *μ* as a function of the sampling proportion, we observed a decline of *μ* with the increase of the sampling proportion (Figure 2D, top panel), similarly to the ratio η (Figure 2A). Interestingly, other metabolites detected in this dataset showed a similar relationship (Figure 2D, bottom panels), albeit with slightly different slopes, potentially explainable by their specific susceptibility to ion suppression. Using quantile regression (Figure 2D, top panel), we estimated *μ* as a function of the sampling proportion.

Based on these findings, we propose to compensate for the ion suppression in spatial single-cell metabolomics by regressing out the impact of sampling proportion onto the MS intensities. We propose to do this by estimating the relationship between *μ* and the sampling proportion for each molecular species by using the quantile regression, followed by dividing the MS intensity by the regression value. After applying this (unsupervised) compensation to the fluorescein ion [M-H]^−^, the correlations between fluorescence and MS intensities increased substantially for both ablated regions (Figure 2B) and cells (Figure 2C) and comparable to the supervised method requiring fluorescence. Taken together, our method for compensating the ion suppression in spatial single-cell metabolomics improves quantitation, can be applied to any molecular species, and delivers the same improvement as the supervised method requiring fluorescence values as the ground truth.

Next, we applied this method for ion suppression to the single-cell metabolomics dataset of co-cultured mCherry-expressing HeLa and GFP-expressing NIH3T3 cells (Rappez *et al.*, 2021). Compensating for the ion suppression led to an overall improvement of the data quality, visibly increasing the separation between cell types in the UMAP visualisation (Figure 2E) as well as reducing the cell-type intermixing (Figure 2F).

### Effect of drop-outs on quantitation

Another potential factor affecting quantitation is the presence of zero MS intensities of fluorescein, in single-cell analyses referred to as drop-outs. Drop-outs can occur due to either biological heterogeneity or technical artefacts (*e.g.* intensities below the limit of detection, variability in mass spectrometry accuracy, errors in signal processing). Depending on their nature, multiple approaches are reported on how to deal with drop-outs in single-cell transcriptomics (Patruno *et al.*, 2021), but no recommendations are available for single-cell metabolomics.

First, we investigated the nature of drop-outs by comparing fluorescent and MS intensities for the ablated regions (Figure 3A). If drop-outs resulted from metabolite intensities being below the detection limit, one would observe low fluorescence signals for those ablated regions. This is, however, not entirely the case in our data, with mass spectrometry zeros present for ablated regions with both low and high fluorescence. Moreover, the distribution of fluorescent intensities for the MS-zeroed ablated regions (“drop-outs”) demonstrates a clear bimodality, similar to the distribution of all (both zero- and non-zero) values (Figure 3B). This finding suggests that zeros are detected not only for the signals below the limit of detection, but additionally appear to be randomly distributed in the dataset (also referred to as Missing Completely at Random, MCAR) (Sterne *et al.*, 2009). If the majority of the zeros are of MCAR-nature, filtering out zeros could improve pixel-cell deconvolution. Applying this filtering approach on our data by omitting ablated regions with zero MS values for fluorescein, however, did not further improve the correlations between fluorescence and MS intensities for neither ablated regions nor cells (Figure 3C,D).

**Figure 3.**
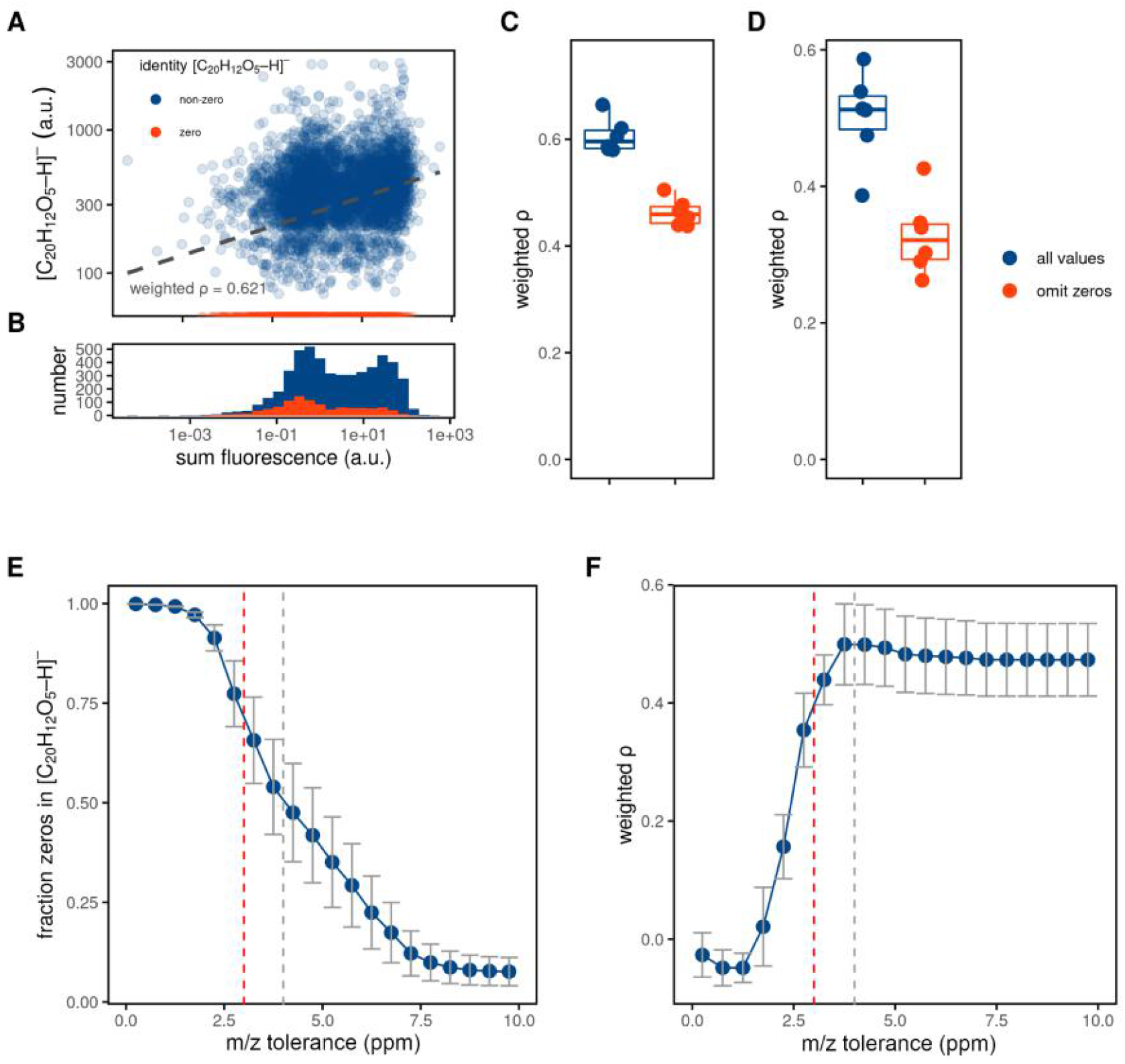
Investigating mass spectrometry drop-outs. **A.** Representative scatter plot (belonging to one replicate) showing relations between the fluorescence intensities and MS intensities for fluorescein, measured for the ablated regions. B. Histogram of the fluorescence intensities for ablated regions with non-zero (blue) and zero (red) MS intensities for the fluorescein ion. **C-D**. Boxplots with density-weighted Spearman correlations between fluorescence and MS intensities of fluorescein either for the ablated regions **(C)**or cells **(D)**using the WA method (sampling proportion of 0.3, m/z tolerance of 4 ppm) for 6 replicates, using all ablated regions (blue) or only those with non-zero MS intensities for fluorescein (red) from panels A and B. **E-F**. The effect of increasing peak integration m/z tolerance on the fraction of zeros in the input measurements **(E)**and correlation between fluorescence and MS intensity for fluorescein**(F)**. Pixel-cell deconvolution methods were performed as described for panel D. Data points indicate the mean values and standard deviation of the 6 replicates, the dashed lines show the 3 ppm (red, default in METASPACE) or 4 ppm tolerance found to be optimal for this experiment (grey).

In imaging MS, missing values with MCAR characteristics may be caused by insufficient and pixel-variable mass accuracy. To this end, we tested if the number of zeros assigned to the fluorescein ion [M-H]^−^ decreases with increasing m/z tolerance. Indeed, the fraction of zeros decreased (Figure 3E), accompanied by increasing quantitation (Figure 3F). Although the fraction of zeros further decreased when using the m/z tolerance higher than 4 ppm, this did not lead to a better quantitation, with the correlation between fluorescence and MS intensity plateauing at the 4 ppm m/z tolerance. Notably, this tolerance is higher than the 3 ppm tolerance used by default for metabolite annotation in METASPACE (Palmer *et al.*, 2017) (see Discussion).

### Optimising filtering parameters for sampling proportion and specificity

Finally, we tested whether excluding specific pixels can improve quantitation. First, we calculated the mean density-weighted Spearman correlation as a function of the sampling proportion cut-off, which excludes input measurements with lower ablated region–cell overlap proportions than the cut-off (Figure 4A). The correlation (blue line) was maximised at the cut-off of 0.3, at the expense of a decrease of the total number of cells that could be assigned (red line). Higher cut-offs resulted in both lower pixel-cell deconvolution quality as well as in much lower number of cells. Next, using a constant sampling proportion cut-off of 0.3, we calculated the mean density-weighted Spearman correlation as a function of the sampling specificity cut-off (Figure 4B) used to exclude co-ablated regions. In contrast to our expectations, increasing the cut-off, and thus excluding co-ablated regions, did not considerably improve pixel-cell deconvolutions, possibly explained by the accompanying decrease of the number of assigned cells. Furthermore, we calculated the correlation as a function of both cut-offs, shown as a heatmap in Figure 4C. We observed two local maxima: at the sampling proportion cut-offs of 0.3 and 0.75 (correlations > 0.5). However, the latter one was achieved for the sampling specificity cut-off of 1 and resulted in the loss of about 80% of all cells (Figure 4D). In summary, the sampling proportion cut-off of 0.3 and the sampling specificity cut-off of 0 were found to be optimal, resulting in the maximal correlation between fluorescence and MS yet retaining a majority of cells.

**Figure 4.**
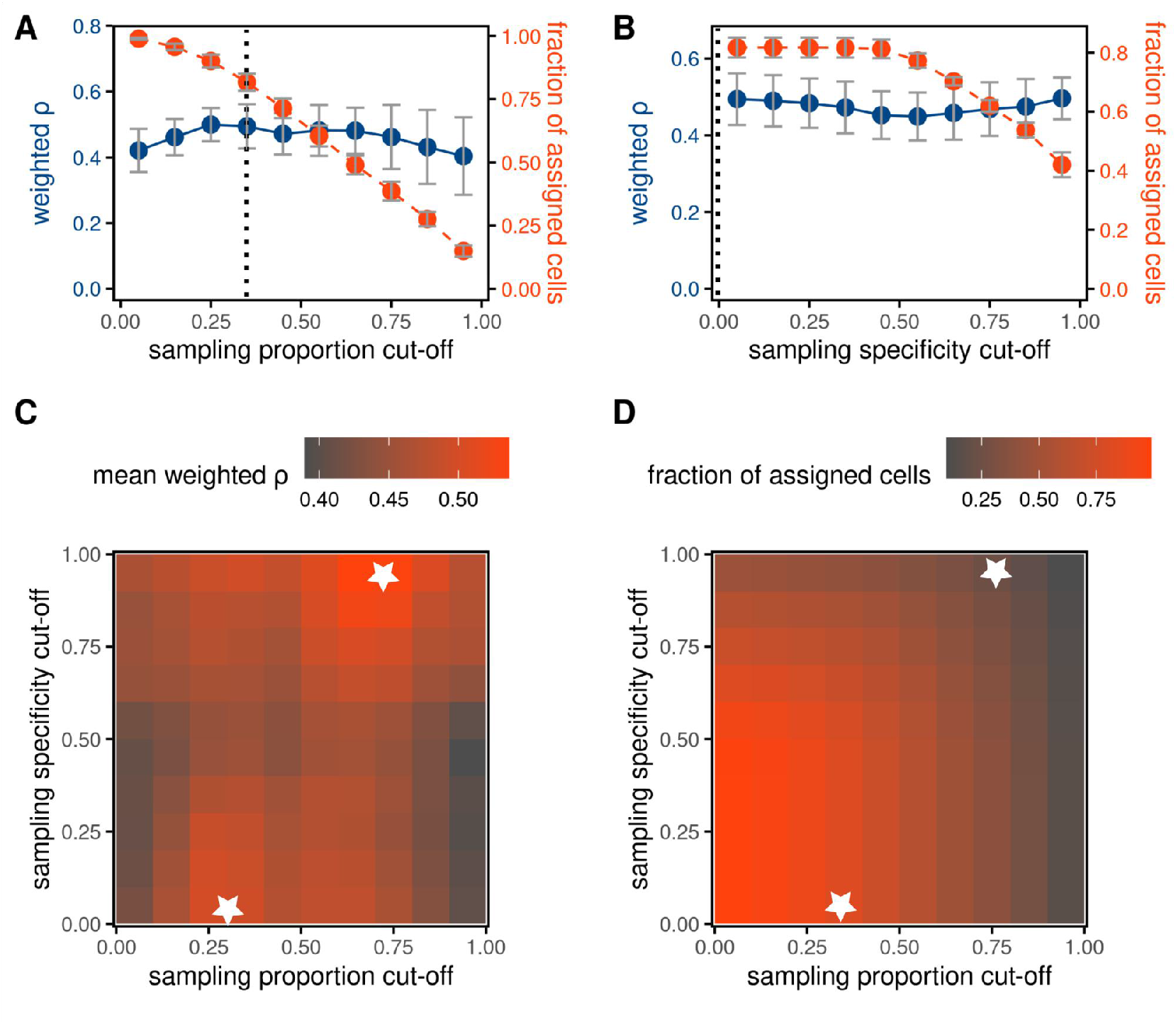
Optimising parameters of the weighted average pixel-cell deconvolution. **A-B.** The effect of increasing the sampling proportion cut-off **(A)** or sampling specificity cut-off **(B)** on the mean density-weighted Spearman correlation *ρ* (blue) between fluorescence and MS intensities of fluorescein (deconvolution method WA, m/z tolerance of 4 ppm) as well as the fraction of cells with assigned MS-signals (red). Data points indicate the mean values and standard deviation of the 6 replicates, dashed lines the optimal values. **C-D**. Heatmaps of mean weighted Spearman correlation **(C)** or fraction of assigned cells **(D)** as function of sampling (x-axis) or specificity (y-axis) proportion cut-offs. White stars indicate the values maximising the mean weighted correlation *ρ*.

## Discussion

We proposed a novel approach to evaluate quantitation in spatial single-cell metabolomics by using a fluorescent dye detectable by mass spectrometry. This helped us assess the method for pixel-cell deconvolution, in particular the earlier proposed weighted average method (WA) (Rappez *et al.*, 2021). Additionally, we evaluated whether an alternative approach using linear inverse modelling (LIM) can improve the quantitation. We used fluorescein, the fluorescent product of fluorescein diacetate that can be measured by both fluorescence microscopy and MALDI-imaging mass spectrometry.

Applying this approach to evaluate pixel-cell deconvolution methods led us to surprising findings. First, we expected superior performance by our newly proposed LIM method as this method is designed to better handle ablated regions overlapping with multiple cells. However, the earlier proposed WA method (Rappez *et al.*, 2021) outperformed the LIM method. As required for solving linear inverse problems, the overall data sets contained more known measurements (ablated regions) than unknown variables (single-cell intensities). After closer inspection, however, we found that the data harboured some local undetermined networks (groups of neighbouring co-ablated cells, with more cells than ablated regions).Underdetermined systems have a solution space rather than one solution and might therefore result in unpredictable outcomes when using the LIM approach. To deal with this, we proposed a mixed model where LIM would be initialised with WA values but this did not lead to improvements. We hypothesise that the WA method demonstrates superior results as it may be more robust in handling noise and non-linear effects present in MALDI imaging MS data.

A more detailed examination of factors influencing the MS-intensities identified a striking intensity overestimation for fluorescein in areas only partly overlapping with cells, suggesting the effect ion suppression. We found that TIC-normalisation of the MS-intensities partly counteracted the effect of ion suppression and improved the quantitation. Possibly, using TIC-normalisation could be more beneficial if a larger m/z range was used (we used 300-400 m/z to increase the sensitivity for fluorescein). Furthermore, we developed a fluorescence-relying supervised method to compensate for the effects of ion suppression, which greatly improved quantitation. Unfortunately, applying the same method for molecules that do not have a fluorescent readout is not feasible, as it relies on measurements of a complementary modality such as fluorescence. Applying the compensation method trained on fluorescein to other analytes is not necessarily justified as ion suppression can be molecule- and context-dependent. However, we observed that the ion suppression can be compensated using an unsupervised method that improved quantitation equally well. Using the unsupervised method does not require any additional measurements and can be applied to the available data and compensates for the ion suppression effect in a metabolite-specific manner. This provides an easy-to-adopt and data-driven approach to compensate for the effects of ion suppression in imaging MS-based spatial single-cell metabolomics.

We also found that increasing the m/z tolerance for the fluorescein ion [M-H]^−^ up to 4 ppm slightly improved the quantitation. This tolerance is slightly higher than the tolerance of 3 ppm that we use by default when using the Orbitrap analyzer and which is recommended in METASPACE for metabolite annotation (Palmer *et al.*, 2017). Closer inspection revealed a small m/z shift of the peak of the fluorescein ion [M-H]^−^. Surprisingly, this m/z shift appeared to be specific to fluorescein, as we did not observe m/z shifts for other analytes. As a result, we do not advise increasing the m/z tolerance in general as it can lead to false positive annotations or a reduction of the number of annotations at the fixed level of the false discovery rate.

It is tempting to generalise the optimizations in this study and to apply them to other MALDI-based single-cell metabolomics experiments. On one hand, the optimal values of parameters (*e.g.* sampling proportion cutoff) identified in this study are in line with our previous study of co-cultures of HeLa and NIH3T3 cells (Rappez *et al.*, 2021). On the other hand, it should be noted that fluorescein has properties that are unique to the compound, including a uniform distribution within cells and a low delocalisation during MALDI-acquisition. For metabolites encountered in biological samples, these characteristics might be different. Furthermore, compared to the co-culture dataset of (Rappez *et al.*, 2021), the data in our study has a relatively large proportion of ablated regions that do not completely overlap with cells: about 97% of all cell-associated ablated regions have overlaps smaller than 100% (Figure S1).This large percentage is probably the result of the experimental procedure, in which we immobilised the cells by centrifugation rather than by cell culture. While cells after longer culture have a more protruded morphology, our cells had a spherical shape and were small in diameter. We suspect that different ratios between partly and fully overlapping ablated regions could shift the optimal filtering parameters prior pixel-cell deconvolution. This is illustrated by a different optimum for sampling specificity cut-off (of 0.9) in the HeLa and NIH3T3 data set (Rappez *et al.*, 2021), *versus* a cut-off of 0 in our experiment. Whenever possible, we suggest avoiding a high occurrence of partly overlapping ablated regions for future experiments, as we observed that ion suppression is most pronounced in these regions. For instance, this can be achieved by decreasing the diameters of the laser-ablated regions, although this can lead to decreased sensitivity and increased runtime, or by using larger cells.

The observed correlations between fluorescent and MS intensities were considerably lower than one. This could have been caused by limitations of MALDI as well as other factors. For example, fluorescence microscopy has a limited dynamic range and potentially did not lead to a full linear response to the fluorescein concentration. The image registration accuracy, which was evaluated visually, and object segmentation quality might have played a role. In addition, we simplified the computational analysis by assuming a uniform energy distribution of the MALDI-laser. This is, however, most likely not the case. Making reliable estimations of these distributions is challenging and was not considered in this study.

In summary, we evaluated pixel-cell deconvolution performance by making use of the fluorescent dye fluorescein. We found that the pixel-cell deconvolution method WA outperformed the LIM method and that quantitation suffers from dependence on the ablated area sampling proportion likely caused by ion suppression. We proposed an unsupervised method to compensate for the effect of the ion suppression. Finally, we identified optimal parameters of SpaceM processing steps before pixel-cell deconvolution.

## Materials & Methods

### Cell culture

HeLa cells (ATCC) were cultured at 37°C and 5% CO_2_in regular Dulbecco’s Modified Eagle Medium (Gibco) containing 10% fetal bovine serum. For the experiments, cells were plated on 8-well dishes (Nunc) and incubated with 25, 50, 75 or 100 μM FDA (Sigma-Aldrich) for 30 minutes at 37°C. Subsequently, cells from all dishes were trypsinized with 0.25% trypsin-EDTA (Gibco), resuspended in medium and pooled into one tube. Then, cell suspensions were deposited at approximately 15,000 cells per well at 1,000 rpm for 10 minutes. After aspiration of the medium, slides were wrapped in aluminium foil to protect them from light, vacuum-desiccated for 1 hour, and marked with fiducials as described earlier (Rappez *et al.*, 2021).

### Experimental steps SpaceM

The experimental protocols of the SpaceM procedure (MALDI, pre- and post-MALDI microscopy) were performed as described previously (Rappez *et al.*, 2021), with the following modifications. Before MALDI acquisition, the slide was spayed with 1,5-diaminonapthalene (DAN, obtained by Sigma-Aldrich) by a TM sprayer (HTX) with these parameters: temperature, 80°C; number of passes, 8; flow rate, 0.07 mL min^−1^; velocity,1,350 mm min^−^1; track spacing, 3 mm min^−1^; pattern, CC; pressure, 10 psi; gas flow rate, 5 L min^−1^; drying time, 15 s and nozzle height, 41 mm. The estimated matrix density was 0.00311 mg mm^−2^. MALDI acquisition was performed using a Q-Exactive Plus mass spectrometer (ThermoFisher Scientific), equipped with an AP-SMALDI5 source (Transmit), in the negative mode using a mass range between m/z 300 - 400 (resolving power R = 140,000 at m/z 200). The pixel step size of the MALDI stage was 25 μm and the acquisition areas were 100 by 100 pixels. The MALDI laser attenuator was set to 33°. Raw acquisition files were converted to centroided imzML files using Alan Race’s converter (Race, Styles and Bunch, 2012). We used the METASPACE cloud software (https://metaspace2020.eu) to perform metabolite annotation using false discovery rate-controlled annotation as published earlier (Palmer *et al.*, 2017) against the CoreMetabolome v3 database. METASPACE was not able to annotate fluorescein at FDR 50% in the two of the initial eight data sets: these data sets were excluded.

### Computational pre-processing

The computational steps of the SpaceM procedure was performed as described previously (Rappez *et al.*, 2021), by performing (i) the image registration between pre- and post-MALDI microscopy images, (ii) the registration between the METASPACE dataset and post-MALDI microscopy, (iii) the segmentation of single cells (iv) and ablated regions, and (v) the calculation of the overlaps between cell and ablated region shapes, required to determine the sampling proportions, sampling specificities, and specific sampling proportions (Figure S2). This pre-processing was done using in-house software which mostly follows the open-source implementation from (Rappez *et al.*, 2021). Compared to the previously described version of SpaceM, there were some modifications. Briefly, the image registration between pre- and post-MALDI microscopy images was performed using the iterative closest point method (Besl and McKay, 1992). For all datasets, the registration accuracy was visually inspected by overlaying the fluorescein channels of both images. For single-cell segmentation, we made use of CellPose (Stringer *et al.*, 2021) with the following parameters: cell diameter, 30.0; flow threshold, 0.4; cell probability threshold, 0.0. The ablated regions were segmented by first visually guiding a 100 by 100 regular grid on the ablated regions of the post-MALDI microscopy brightfield channel. Then, we performed segmentation by defining circular shapes with a diameter of 30 pixels (equals about 19 μm) on the grid.

### Pixel-cell deconvolution methods

We made use of two distinct pixel-cell deconvolution methods. The weighted average (WA) method was described previously (Rappez *et al.*, 2021). Briefly, MS-signals from ablated regions (thus, MALDI-pixels) associated with the cell of interest are first divided by their sampling proportion (the fraction in which the ablated region overlaps with any cell, Figure S2). Then, the assigned MS-signal S*_cell_* is calculated by the weighted average with the sampling specificity (fraction of the sampling proportion that overlaps with the cell-of-interest, Figure S2) as weights:

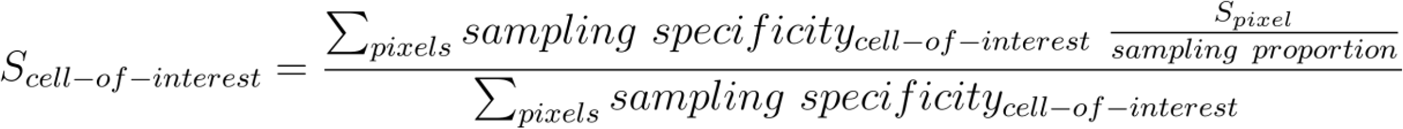

For the the linear inverse modelling (LIM) method, we store our [fluorescein-H]^−^ signals from the MALDI-pixels as vector *p* and the specific sampling proportions (the Hadamard product of sampling proportions and sampling specificities, Figure S2) in an *n* x *m* matrix *S,* where *n* is the number of pixels and *m* the number of cells. The cellular [fluorescein-H]^−^signals *c* can be determined by solving the following equation:

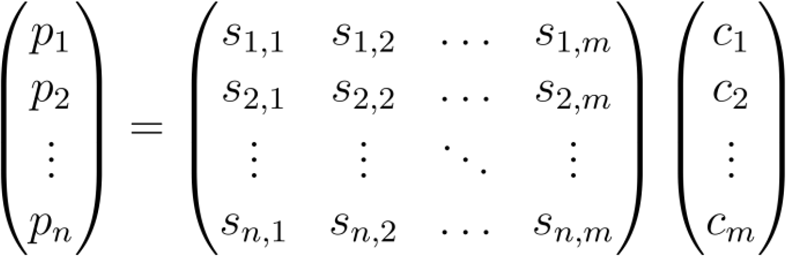

After we constrain *c* to be non-negative, *c* is estimated using linear inverse modelling (LIM) as implemented in the R-packages *nnls* v.1.4 and *LimSolve* v.1.5.6 (Soetaert, Meersche and Oevelen, 2009).

We used a combined approach (MIX) as follows. First, both the LIM and WA methods are used in parallel. Then, of each cell, it is determined whether this cell is part of either an under- or over-determined subnetwork. To this end, all cells and ablated regions are added to a network as nodes, connected by edges between cells and ablated regions in case they are overlapping (with sampling proportions of > 0.3). Using R-package *igraph* (v.1.3.1), distinct subnetworks with no links to other subnetworks are determined. Cells are defined as underdetermined when they are members of a subnetwork with more cells than ablated regions. Finally, all underdetermined cells are assigned with the values obtained from the WA method, while all others are obtained from the LIM method.

In all pixel-cell deconvolution methods, N/A (not a number) values are returned when cells are not associated to any included ablated regions. Zeros are returned when the result of the pixel-cell deconvolution assignments are zero.

### Evaluation of the pixel-cell deconvolution methods

Initial measurements and pixel-cell deconvolution methods were evaluated by calculating the weighted Spearman correlation coefficients (R-package *wCorr* v.1.9.5) between the fluorescein fluorescence and MS intensities of ablated regions (MALDI-pixels) or cells. The weights were used to adjust for the different distributions of fluorescein in the ablated regions *versus* the cells (for the ablated regions, the vast majority is extracellular and zero, whereas this is not the case for cells). Therefore, we defined weights *ω* as negatively proportional to the normalised local density *ρ* of the fluorescence vector:

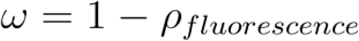

The evaluation and UMAP projection of the co-culture dataset was done with R-package *Seurat* (v.4.1.1).

## Code and Data availability

All MALDI-imaging MS data including the metabolite annotations are freely available through METASPACE under the following project link:https://metaspace2020.eu/project/FDA_evaluation.

The co-culture dataset was previously published (Rappez *et al.*, 2021) and is available in the MetaboLights repository under the accession number MTBLS78 (https://www.ebi.ac.uk/metabolights/MTBLS78). All analyses were run in RStudio v.2022.02.2 (R v.4.2.0) and are available as an RMarkdown document (Supplemental Information 1). All figures were built in RStudio and/or Inkscape v0.92.2.

