## Supplemental Figures for "Increasing quantitation in spatial single-cell metabolomics by using fluorescence as ground truth"

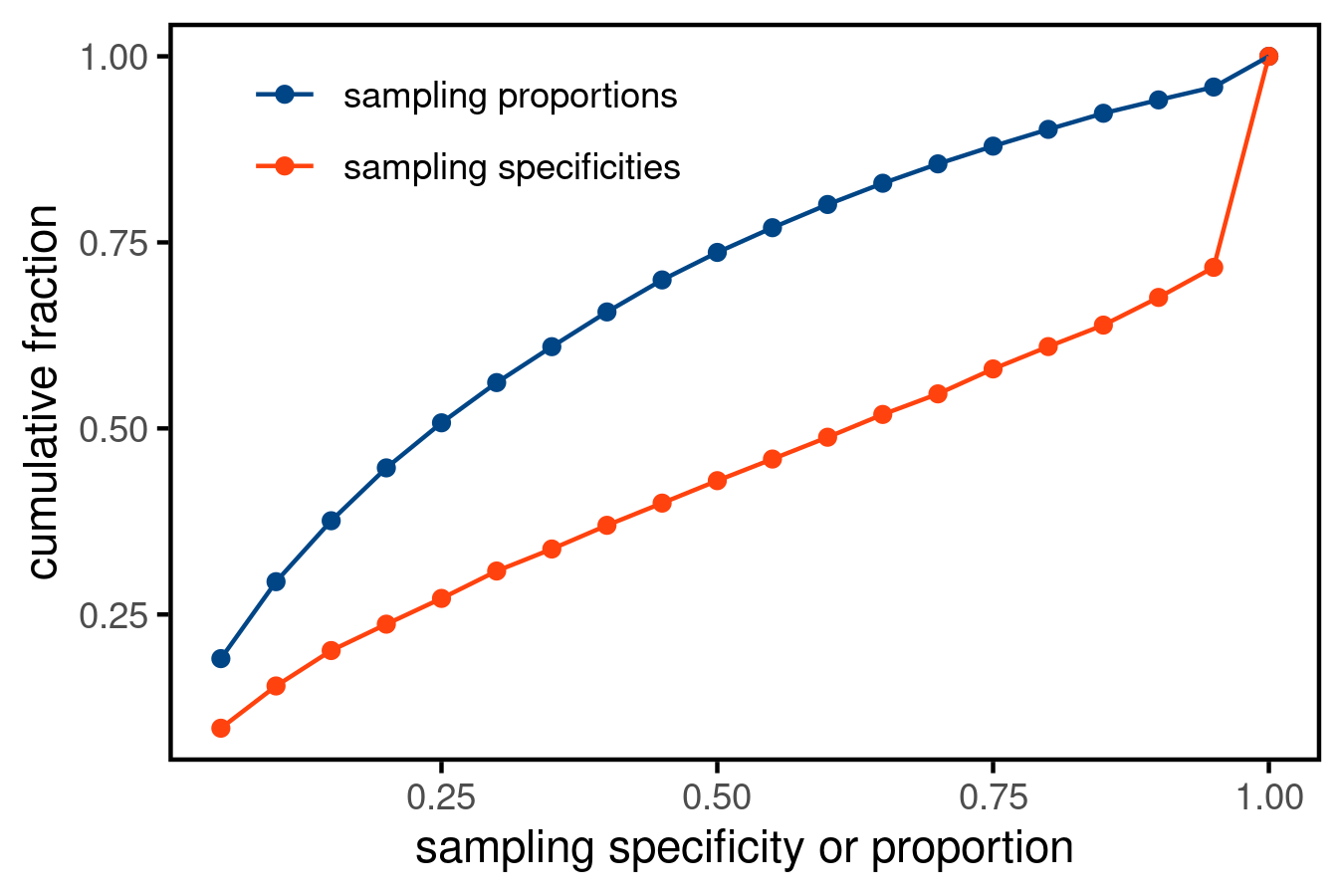

**Figure S1.** Cumulative fraction of sampling proportion (blue) or sampling specificity (red) of all ablated regions.

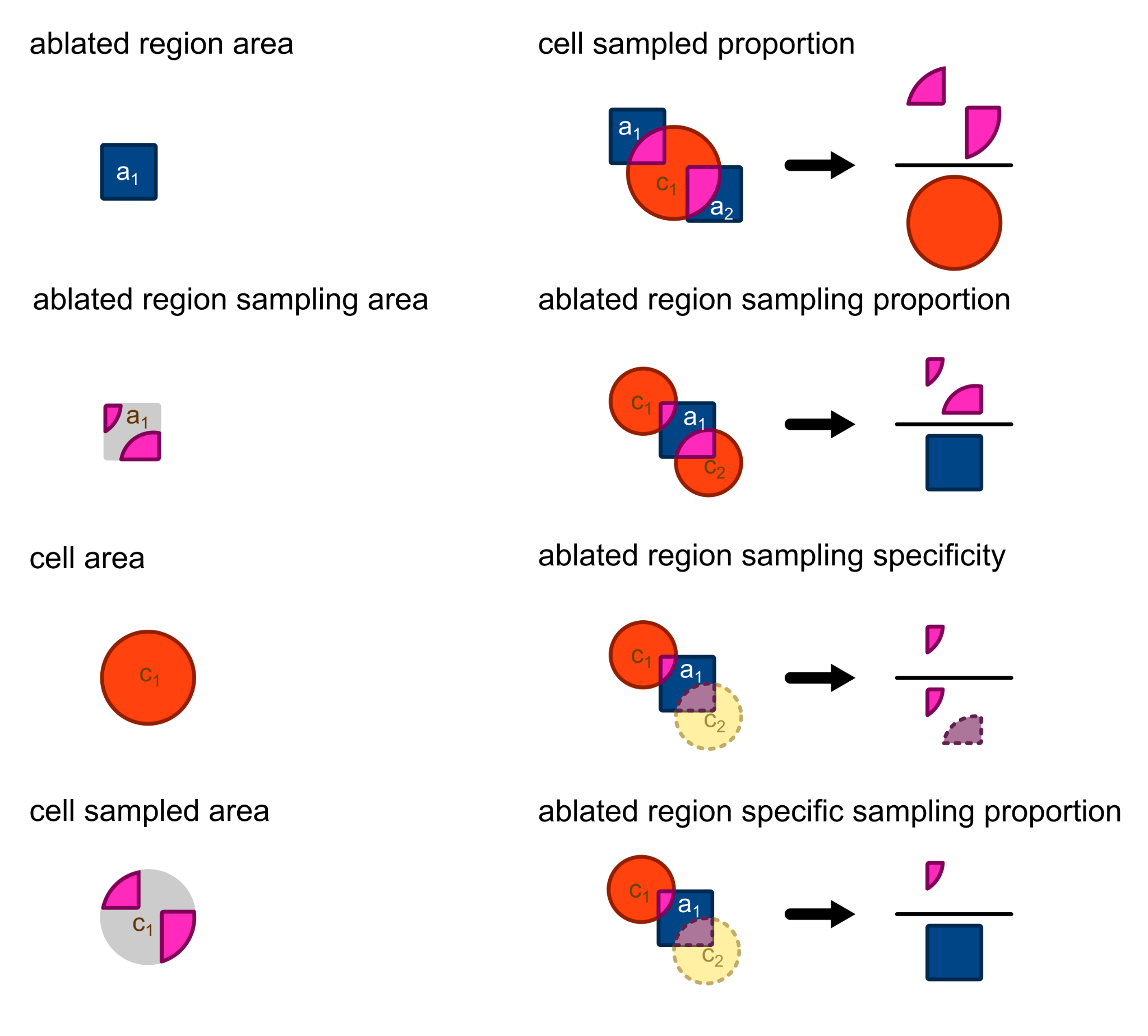

**Figure S2.** Definitions of morphological shapes and measures used in SpaceM.
