## Supplemental Information 1 for "Increasing quantitation in spatial single-cell metabolomics by using fluorescence as ground truth"

### Supplemental Information 1, R-code used for analysis

2022-06-08

```
library(dplyr)
library(ggplot2)
library(ggrepel)
library(tidyr)
library(paletteer)
library(ggthemes)
library(rawrr)          ## https://bioconductor.org/packages/release/bioc/html/rawrr.html
library(pbapply)
library(igraph)
```

```
### list all csv AM-matrice
files <-
  paste0("",
         list.files(
           path = ".",
           pattern = 'spatiomolecular_matrix_w_API_tic',
           recursive = T
         ))
files <- files[grepl("ablation_mark_analysis", files)]
files
```

```
## [1] "2022-02-18_FDA_SpaceM/w2/analysis/ablation_mark_analysis/spatiomolecular_matrix_w_API_tic.csv"
## [2] "2022-02-18_FDA_SpaceM/w3/analysis/ablation_mark_analysis/spatiomolecular_matrix_w_API_tic.csv"
## [3] "2022-02-18_FDA_SpaceM/w4/analysis/ablation_mark_analysis/spatiomolecular_matrix_w_API_tic.csv"
## [4] "2022-02-18_FDA_SpaceM/w5/analysis/ablation_mark_analysis/spatiomolecular_matrix_w_API_tic.csv"
## [5] "2022-02-18_FDA_SpaceM/w6/analysis/ablation_mark_analysis/spatiomolecular_matrix_w_API_tic.csv"
## [6] "2022-02-18_FDA_SpaceM/w7/analysis/ablation_mark_analysis/spatiomolecular_matrix_w_API_tic.csv"
## [7] "2022-02-18_FDA_SpaceM/w8/analysis/ablation_mark_analysis/spatiomolecular_matrix_w_API_tic.csv"
```

```
### load all AM data into one data.frame
data <-
  sapply(files, function(file_i) {
    data_i <- read.csv(file = file_i)
    data_i$file <-
      gsub(
        "\\Q../\\E|/analysis/ablation_mark_analysis/spatiomolecular_matrix_w_API_tic.csv",
        "",
        file_i
      )
    return(data_i)
  }) %>% bind_rows
```

```
data <-
data %>% group_by(file) %>%
  mutate(C20H12O5.H_present = !all(is.na(C20H12O5.H))) %>% ungroup() %>%
  filter(C20H12O5.H_present) %>% select(-`C20H12O5.H_present`)
```

```r
## RAW file of interest
rawfile1 <- 'FDA_raw_files/2021-11-25_FDA_W2_100x100_a25ss30rf100_DANneg.RAW'
rawfile2 <- 'FDA_raw_files/2021-11-25_FDA_W345678_100x100_a25ss30rf100_DANneg.RAW'

## m/z of target ions
mass_target <- c(331.0612)
tolerance_i <- 8

data_w_ppm <-
    list({
      RawData <- readFileHeader(rawfile = rawfile2)
      FLU_signal <-
        readChromatogram(
          rawfile = rawfile2,
          mass = mass_target,
          tol = tolerance_i,
          type = "xic"
        )

      data %>% filter(file %in% paste0("2022-02-18_FDA_SpaceM/w", 3:7)) %>%
        mutate(C20H12O5.H_4ppm = FLU_signal[[1]]$intensities[1:50000],
               ppm_tolerance = tolerance_i / 2)
    }, {
      RawData <- readFileHeader(rawfile = rawfile1)
      FLU_signal <-
        readChromatogram(
          rawfile = rawfile1,
          mass = mass_target,
          tol = tolerance_i,
          type = "xic"
        )

      data %>% filter(file %in% "2022-02-18_FDA_SpaceM/w2") %>%
        mutate(C20H12O5.H_4ppm = FLU_signal[[1]]$intensities[1:10000],
               ppm_tolerance = tolerance_i / 2)
    }) %>% bind_rows()
```

```r
plot(data_w_ppm$C20H12O5.H,data_w_ppm$C20H12O5.H_4ppm)
```

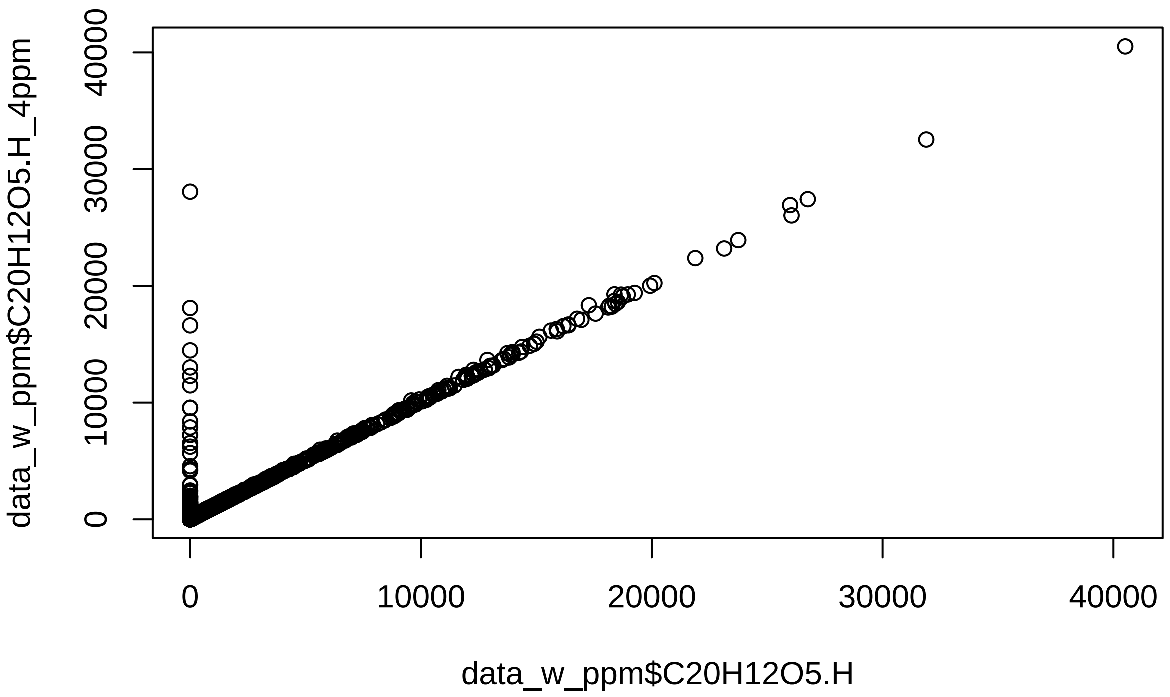


```r
library(pbmcapply)
library(nnls)
library(tidyr)
library(wCorr)

## loading SpaceM normalization functions
source('normalization_functions.r')
```

```r
## define locations of datasets for proportion matrix retreival
data_folders <- paste0("", unique(data$file), "/")
data_folders <-
  unique(gsub(
    "analysis/overlap_analysis./ablation_mark.regions.csv",
    "",
    data_folders
  ))


## obtain and calculate_proportion_matrixs in a list over de datasets

prop_spec_matrix <-
sapply(data_folders, function(sample_location){
    prop_spec_matrix <- calculate_proportion_matrix(sample_location = sample_location)
    return(prop_spec_matrix)
}, simplify = FALSE)
```

```r
## just to double-check that the order of files in both sets is the same

data.frame(AMdataset = data_w_ppm %>% group_by(file) %>% summarize(file = file[1]) %>% pull(file),
           prop_spec_matrix = names(prop_spec_matrix))
```

```
##                   AMdataset          prop_spec_matrix
## 1 2022-02-18_FDA_SpaceM/w2 2022-02-18_FDA_SpaceM/w2/
```

```
## 2 2022-02-18_FDA_SpaceM/w3 2022-02-18_FDA_SpaceM/w3/
## 3 2022-02-18_FDA_SpaceM/w4 2022-02-18_FDA_SpaceM/w4/
## 4 2022-02-18_FDA_SpaceM/w5 2022-02-18_FDA_SpaceM/w5/
## 5 2022-02-18_FDA_SpaceM/w6 2022-02-18_FDA_SpaceM/w6/
## 6 2022-02-18_FDA_SpaceM/w7 2022-02-18_FDA_SpaceM/w7/
```

```
datasets_names <- data_w_ppm %>% group_by(file) %>% summarize(file = file[1]) %>% pull(file)

## iterate over:
## datasets_names

cells <-
  sapply(datasets_names, function(dataset_i) {
    ## in every iteration, different normalizations are assessed

    ## first, retreive prop and spec matrix for dataset_i
    prop_spec_matrix_OI <-
      prop_spec_matrix[[paste0(dataset_i, '/')]]


    ## now, the normalization parameters
    list({
      ## Rappez_et_al method; TIC-norm, zeros included

      AM_ion_intensity_OI <-
        data_w_ppm %>% filter(file ==  dataset_i) %>%                       ## filter dataset of intere
        mutate(C20H12O5.H = C20H12O5.H_4ppm / TIC) %>%                ## normalize C20H12O5.H  by TIC
        pull(C20H12O5.H)

      cell_ion_intensity <-
        cell_normalization_Rappez_et_al(
          overlap_proportion_matrix = prop_spec_matrix_OI$overlap_proportion_matrix,
          overlap_specificity_matrix = prop_spec_matrix_OI$overlap_specificity_matrix,
          AM_ion_intensity = AM_ion_intensity_OI,
          AM_proportion_treshold_global = 0.3,
          skip_AM_zeros = FALSE                                         ## keep zeros in AM vector
        )$cell_intensities

      data.frame(cell_id = rownames(cell_ion_intensity),
                 C20H12O5.H_calculated = cell_ion_intensity,
                 method = 'weighted average',
                 AM_normalization = 'TIC',
                 AM_filter = '0.3',
                 AM_omit_zeros = "keep zeros",
                 file = dataset_i)
    },{
      ## NNLS method; TIC-norm, zeros included

      AM_ion_intensity_OI <-
        data_w_ppm %>% filter(file ==  dataset_i) %>%                       ## filter dataset of intere
        mutate(C20H12O5.H = C20H12O5.H_4ppm / TIC) %>%                     ## normalize C20H12O5.H by T
        pull(C20H12O5.H)

      cell_ion_intensity <-
```

```
    cell_normalization_NNLS(
      overlap_proportion_matrix = prop_spec_matrix_OI$overlap_proportion_matrix,
      AM_ion_intensity = AM_ion_intensity_OI,
      AM_proportion_treshold_per_cell = 0.3,
      skip_AM_zeros = FALSE                                    ## keep zeros in AM vector
    )$cell_intensities


  data.frame(cell_id = names(cell_ion_intensity),                 ## be carefull, somewhat inconsis
             ## here use 'names', not 'rownames'!
             C20H12O5.H_calculated = cell_ion_intensity,
             method = 'linear inverse modeling',
             AM_normalization = 'TIC',
             AM_filter = '0.3',
             AM_omit_zeros = "keep zeros",
             file = dataset_i)
},{
  ## combined; TIC-norm, zeros included

  AM_ion_intensity_OI <-
    data_w_ppm %>% filter(file ==  dataset_i) %>%                     ## filter dataset of intere
    mutate(C20H12O5.H = C20H12O5.H_4ppm / TIC) %>%                   ## normalize C20H12O5.H by T
    pull(C20H12O5.H)

  cell_ion_intensityNNLS <-
    cell_normalization_NNLS(
      overlap_proportion_matrix = prop_spec_matrix_OI$overlap_proportion_matrix,
      AM_ion_intensity = AM_ion_intensity_OI,
      AM_proportion_treshold_per_cell = 0.3,
      skip_AM_zeros = FALSE                                    ## keep zeros in AM vector
    )$cell_intensities

  cell_ion_intensityWA <-
    cell_normalization_Rappez_et_al(
      overlap_proportion_matrix = prop_spec_matrix_OI$overlap_proportion_matrix,
      overlap_specificity_matrix = prop_spec_matrix_OI$overlap_specificity_matrix,
      AM_ion_intensity = AM_ion_intensity_OI,
      AM_proportion_treshold_global = 0.3,
      skip_AM_zeros = FALSE                                    ## keep zeros in AM vector
    )$cell_intensities


  ## which cells are underdetermined?

  slim_overlap_proportion_matrix <- prop_spec_matrix_OI$overlap_proportion_matrix[,colSums(prop_spe

  edge_list <-
    pblapply(seq(dim(slim_overlap_proportion_matrix)[1]), function(i) {
      AM_i <- colnames(slim_overlap_proportion_matrix)[slim_overlap_proportion_matrix[i, ] > 0.3]

      sapply(AM_i, function(j) {
        data.frame(from = rownames(slim_overlap_proportion_matrix)[i],
                   to = j)
```

```r
          }, simplify = F) %>% bind_rows()

        })

      edge_list <- edge_list[sapply(edge_list, function(i){dim(i)[1]}) > 0] %>% bind_rows()

      node_list <-
        tibble(id = c(
          rownames(prop_spec_matrix_OI$overlap_proportion_matrix),
          colnames(prop_spec_matrix_OI$overlap_proportion_matrix)
        ))

      cell_AM_network <- graph_from_data_frame(d=edge_list, vertices = node_list, directed=F)

      cell_determined <- data.frame(node_ID = names(V(cell_AM_network)),
                                    community = as.vector(components(cell_AM_network)$membership)) %>%
        group_by(community) %>%
        mutate(
          cell = sum(grepl("cell", node_ID)),
          AM = sum(grepl("AM", node_ID)),
          underdetermined = cell > AM
        ) %>%
        filter(grepl("cell", node_ID))



      ## replace underdetermined values with WA
      cell_ion_intensityNNLS[names(cell_ion_intensityNNLS) %in%
                               cell_determined$node_ID[cell_determined$underdetermined]] <-
        cell_ion_intensityWA[names(cell_ion_intensityNNLS) %in%
                               cell_determined$node_ID[cell_determined$underdetermined]]

      ##

      data.frame(cell_id = names(cell_ion_intensity),                  ## be carefull, somewhat inconsis
                 ## here use 'names', not 'rownames'!
                 C20H12O5.H_calculated = cell_ion_intensityNNLS,
                 method = 'MIX',
                 AM_normalization = 'TIC',
                 AM_filter = '0.3',
                 AM_omit_zeros = "keep zeros",
                 file = dataset_i)
    })  %>% bind_rows()

  }, simplify = FALSE) %>% bind_rows()
```

```r
files <- paste0(list.files(pattern = 'spatiomolecular_matrix.csv', recursive = T) )
files <- files[grepl("single_cell_analysis", files)]
files
```

```
## [1] "2022-02-18_FDA_SpaceM/w2/analysis/single_cell_analysis/spatiomolecular_matrix.csv"
## [2] "2022-02-18_FDA_SpaceM/w3/analysis/single_cell_analysis/spatiomolecular_matrix.csv"
## [3] "2022-02-18_FDA_SpaceM/w4/analysis/single_cell_analysis/spatiomolecular_matrix.csv"
## [4] "2022-02-18_FDA_SpaceM/w5/analysis/single_cell_analysis/spatiomolecular_matrix.csv"
```

```
## [5] "2022-02-18_FDA_SpaceM/w6/analysis/single_cell_analysis/spatiomolecular_matrix.csv"
## [6] "2022-02-18_FDA_SpaceM/w7/analysis/single_cell_analysis/spatiomolecular_matrix.csv"
## [7] "2022-02-18_FDA_SpaceM/w8/analysis/single_cell_analysis/spatiomolecular_matrix.csv"
```

```
### list all csv cell-matrice

cells_SpaceM <-
  sapply(files, function(file_i) {
    cells_SpaceM_i <- read.csv(file = file_i)
    cells_SpaceM_i$file <-
      gsub(
        "\\Q../\\E|/analysis/single_cell_analysis/spatiomolecular_matrix.csv",
        "",
        file_i
      )
    cells_SpaceM_i$cell_id <- paste0('cell#', cells_SpaceM_i$cell_id)
    return(cells_SpaceM_i)
  }) %>% bind_rows

colnames(cells_SpaceM)[colnames(cells_SpaceM) == "C20H12O5.H"] <- 'C20H12O5.H_SpaceM'
```

```
cells <-
  cells %>% left_join(
    cells_SpaceM %>% select(file, cell_id, median_intensity.FITC, cell_area, eccentricity),
    by = c("file", "cell_id")
  )

cells <-  cells %>% group_by(method, AM_normalization,AM_filter,AM_omit_zeros,file) %>%
  mutate(n = length(C20H12O5.H_calculated))
```

```
cell_summary <-
  left_join(
    x = cells,

    y = data_w_ppm %>% group_by(file) %>%

      summarize(
        signal_C20H12O5.H = quantile(x = C20H12O5.H, probs = 1),
        signal_TIC = quantile(x = TIC, probs = 1),
        signal_fluorescein = quantile(x = median_intensity.FITC, probs = 1),
        AM_area = mean(am_area)
      ),
    by = 'file'
  ) %>%
  group_by(method, AM_normalization, AM_filter, AM_omit_zeros, file) %>%
  filter(          ## filter out NAs for the correlations
    !is.na(median_intensity.FITC) &!is.na(C20H12O5.H_calculated)) %>%
  summarize(
    signal_C20H12O5.H = signal_C20H12O5.H[1],
    signal_TIC = signal_TIC[1],
    signal_fluorescein = signal_fluorescein[1],
    AM_area = AM_area[1],
    r = cor(
      x = median_intensity.FITC,
      y = C20H12O5.H_calculated,
```

```
      method = "spearman",
      use = "pairwise.complete.obs"
    ),
    rweight = weightedCorr(
      ## weighted median_intensity.FITC
      x = median_intensity.FITC,
      y = C20H12O5.H_calculated,
      method = "Spearman",
      weights = 1 - n_neighbours(x = median_intensity.FITC) /
        max(n_neighbours(median_intensity.FITC))
    ),
    n = n[1]
  )
```

```
Fig1D <-
  cell_summary %>%  mutate(
    method = gsub("weighted average", "WA", method),
    method = gsub("linear inverse modeling", "LIM", method),
    method = factor(method, levels = c('LIM', 'WA', 'MIX'))
  ) %>% {
    ggplot(data = .,
           aes(x = method,
               y = rweight, color = method)) +
      #geom_vline(xintercept = 3, linetype = 3, color = "gray30")+
      scale_y_continuous(limits = c(0, NA)) +
      geom_boxplot(outlier.colour = NA, show.legend = F) +
      geom_jitter(size = 3,
                  width = .08,
                  show.legend = F) +
      scale_color_paletteer_d("ggthemes::calc") +
      #geom_line(aes(group = file, color = file), linetype = 2, alpha = .7)+
      labs(y = 'weighted  ',  x = "normalization method") +
      theme_minimal() +
      theme(
        axis.text.x = element_text(size = 9),
        axis.title = element_text(size = 11),
        panel.grid = element_blank(),
        axis.ticks = element_line(),
        panel.background = element_rect(colour = "black", size = 1)
      )

  }

Fig1D
```

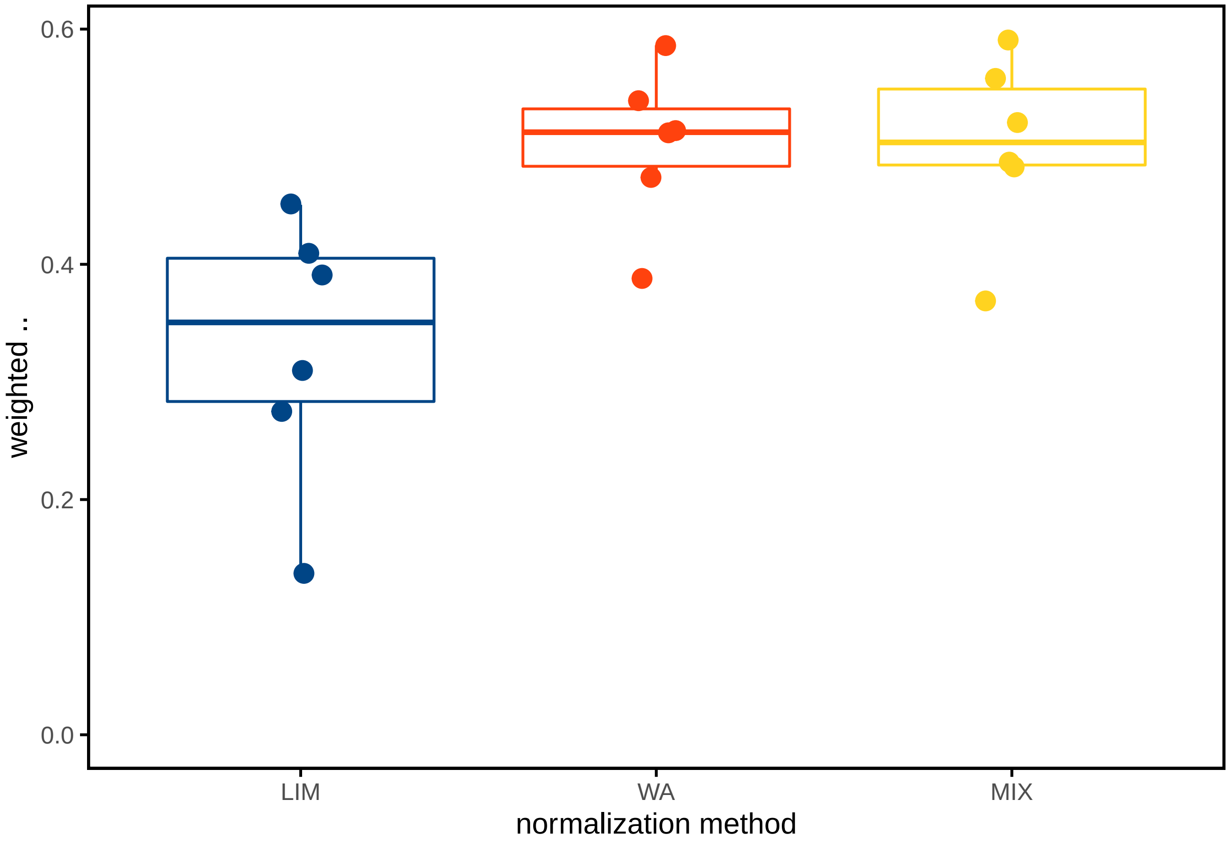


```
Fig1C <-
  cells %>% filter(file == "2022-02-18_FDA_SpaceM/w4") %>%
  filter(method != "MIX") %>%
  mutate(
    method = gsub("weighted average", "WA", method),
    method = gsub("linear inverse modeling", "LIM", method)
  ) %>%
  {
    ggplot(data = .,
           aes(x = median_intensity.FITC,
               y = C20H12O5.H_calculated, color = method)) +
      geom_point(size = 1,
                 alpha = .3,
                 show.legend = F) +
      facet_grid(. ~ method,
                 scales = "free") +
      scale_color_paletteer_d("ggthemes::calc") +
      scale_x_continuous(limits = c(0, .3)) +
      scale_y_continuous(limits = c(0, .01)) +
      geom_text(
        data = .  %>%
          group_by(method, file) %>%
          summarize(
            isNA = sum(is.na(C20H12O5.H_calculated)),
            is0 = sum(C20H12O5.H_calculated == 0, na.rm = T),
            isNum = sum(C20H12O5.H_calculated > 0, na.rm = T),
            total = length(C20H12O5.H_calculated),
```

```
          label  = paste0(
            "\n   non-zeros = ",
            format(isNum, digits = 3),
            "\n   non-assigned = ",
            format(isNA, digits = 3),
            "\n   zeros = ",
            format(is0, digits = 3),
            "\n   total = ",
            format(total, digits = 3)
          )
        ),
      size = 3,
      x = -Inf,
      y = Inf,
      hjust = 0,
      vjust = 1,
      color = "gray30",
      aes(label = label)
    ) +
    labs(x = 'median fluorescence (a.u.)',
         y = expression(assigned ~ "[C"[20] * "H"[12] * "O"[5] * "-H]" ^ "-" ~ (a.u.))) +
    #'assigned [C20H12O5-H]- (a.u.)')+
    #expression(Anthropogenic~SO[4]^{"2-"}~(ngm^-3))
    #expression(assigned~[C20H12O5-H]^- (a.u.))
    theme_minimal() +
    theme(
      axis.text.x = element_text(size = 9),
      axis.title = element_text(size = 11),
      panel.grid = element_blank(),
      axis.ticks = element_line(),
      panel.background = element_rect(colour = "black", size = 1),
      strip.text = element_text(size = 11, face = "bold")
    )
  }

Fig1C
```

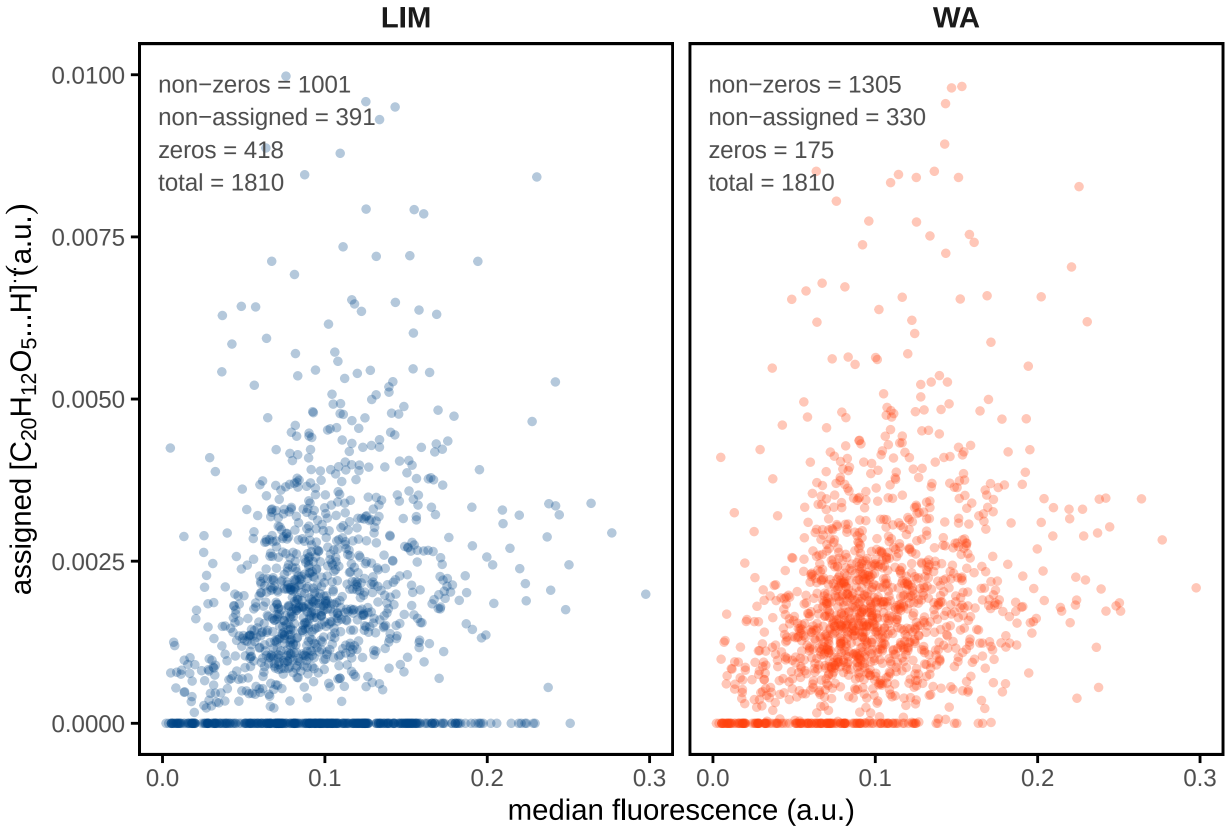


#### simulations

```
library(reshape2)
library(gpclib)
library(cowplot)

set.seed(seed = 2021)
```

```
### functions
rotate_xy <- function(xy, angle, center = c(0, 0)) {
  xRot = center[1] + cos(angle) * (xy[, 1] - center[1]) - sin(angle) * (xy[, 2] - center[2])
  yRot = center[2] + sin(angle) * (xy[, 1] - center[1]) + cos(angle) * (xy[, 2] - center[2])
  cbind(xRot, yRot)
}


convex.poly <- function(nSides, area)
{
  radius <- sqrt((2 * area) / (nSides * sin((2 * pi) / nSides)))
  angle <- (2 * pi) / nSides

  radii <- rnorm(nSides, radius, radius / 10)
  angles <- rnorm(nSides, angle, angle / 10) * 1:nSides
  angles <- sort(angles)

  points <- list(x = NULL, y = NULL)
  points$x <- cos(angles) * radii
  points$y <- sin(angles) * radii
```

```r
  m <- matrix(unlist(points), ncol = 2)
  m <- rbind(m, m[1, ])
  current.area <-
    0.5 * (sum(m[1:nSides, 1] * m[2:(nSides + 1), 2]) - sum(m[1:nSides, 2] *
                                                               m[2:(nSides + 1), 1]))

  points$x <- points$x * sqrt(area / current.area)
  points$y <- points$y * sqrt(area / current.area)

  return (points)
}

regular.poly <- function(nSides, area)
{
  radius <- sqrt((2 * area) / (nSides * sin((2 * pi) / nSides)))

  points <- list(x = NULL, y = NULL)
  angles <- (2 * pi) / nSides * 1:nSides

  points$x <- cos(angles) * radius
  points$y <- sin(angles) * radius

  return (points)

}
```

```r
n <- 5 ### generate n^2 random cells
coordinates <- rep(seq(-n, n, length.out = n), n)
x <- matrix(coordinates, ncol = n, byrow = F)
y <- matrix(coordinates, ncol = n, byrow = T)

## df with x-y coordinates of borders of cells
cells <-
  sapply(1:length(x), function(i) {
    xy <- c(x[i], y[i])
    cell <- data.frame(cell = paste("cell#", i, sep = ""),
                       convex.poly(
                         nSides = round(runif(
                           n = 1, min = 15, max = 30
                         ), 0),
                         area = rnorm(n = 1, mean = 3.5, sd = .3)
                       ))
    cell[2:3] <- t(apply(cell[2:3], 1, function(row) {
      row + xy
    }))

    return(cell)
  }, simplify = FALSE) %>% do.call("rbind", .)

cells$cell <- factor(cells$cell, levels = paste("cell#", 1:length(x), sep = ""))
```

```r
### assign random metabolite X signals

cell_signals <- data.frame(cell = levels(cells$cell),
```

```r
                              signal = runif(length(levels(cells$cell)), min = 0, max = 10))

### assign to cells data.frame

cells$signal <-
  cell_signals$signal[match(cells$cell, cell_signals$cell)]
```

```r
##### generate ablations marks
abl_n <- 6     ### generate abl_n^2 random AMs

coordinates <- rep(seq(-abl_n, abl_n, length.out = abl_n), abl_n)
x <- as.vector(matrix(coordinates, ncol = abl_n, byrow = F))
y <- as.vector(matrix(coordinates, ncol = abl_n, byrow = T))

x <- rotate_xy(cbind(x, y), angle = 0.3)[, 1]
y <- rotate_xy(cbind(x, y), angle = 0.3)[, 2]

abl_size <- 3     ## relative AM diameter

## df with x-y coordinates of borders of AMs
ablations <-
  sapply(1:length(x), function(i) {
    xy <- c(x[i], y[i])
    cell <- data.frame(ablation = paste("ablation#", i, sep = ""),
                       regular.poly(nSides = 40, area = abl_size))
    cell[2:3] <- t(apply(cell[2:3], 1, function(row) {
      row + xy
    }))

    return(cell)
  }, simplify = FALSE) %>% do.call("rbind", .)

ablations$ablation <-
  factor(ablations$ablation, levels = paste("ablation#", 1:length(x), sep = ""))
```

```r
### plot shapes

simFig1 <-
ggplot() +
    coord_fixed() +
    scale_fill_gradient(low = "white", high = "red") +
    geom_polygon(
      data = cells,
      aes(
        x = x,
        y = y,
        group = cell,
        fill = signal
      ),
      show.legend = F,
      color = "gray80",
      size = .5
    ) +
    geom_polygon(
```

```
      data = ablations,
      aes(x = x, y = y, group = ablation),
      color = "black",
      fill = NA, #"white",
      alpha = .4, linetype = 2,
      size = .5,
      show.legend = F
    ) +
    theme_void()+
    theme(panel.background = element_rect(fill = "white", color = NA))

simFig1
```

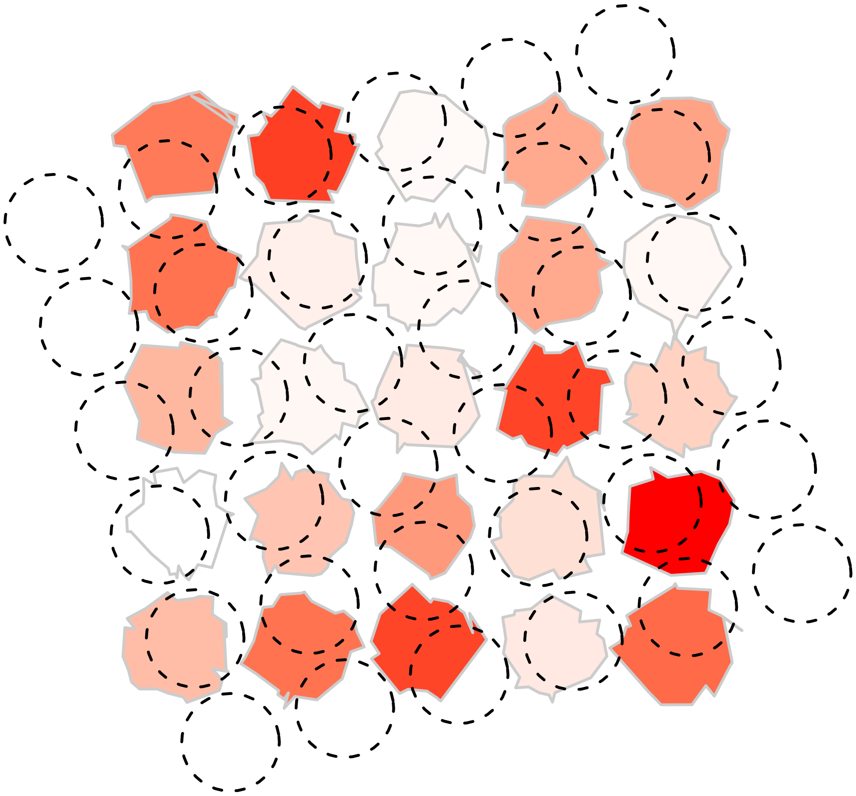

```
###### calculate expected signal in the ablations

## first, formalize polys to measure them as objects
cells_polys <-
  cells %>% group_split(cell) %>% sapply(function(cell) {
    shape <- cbind(cell$x, cell$y)
    shape <- shape[chull(shape), ]
    shape <- as(shape, "gpc.poly")
    shape
  })

ablations_polys <-
  ablations %>% group_split(ablation) %>% sapply(function(ablation) {
    shape <- cbind(ablation$x, ablation$y)
    shape <- shape[chull(shape), ]
```

```
    shape <- as(shape, "gpc.poly")
    shape
  })


## calculate area% of cell_i in abl_i (= am_proportion matrix)
overlap_matrix <-
  sapply(1:length(ablations_polys), function(abl_i) {
    pr <- round(abl_i / length(ablations_polys) * 100, digits = 5)
    # cat(pr)
    # cat(paste(rep(x = "\b", length(pr))), collapse = "")
    sapply(1:length(cells_polys), function(cell_i) {
      area.poly(intersect(ablations_polys[[abl_i]], cells_polys[[cell_i]]))
    }) / area.poly(ablations_polys[[abl_i]])

  })

overlap_matrix <- data.frame(overlap_matrix)
rownames(overlap_matrix) <- levels(cells$cell)
colnames(overlap_matrix) <- levels(ablations$ablation)

overlap_matrix[1:6,1:6]
```

```
##         ablation#1 ablation#2 ablation#3 ablation#4 ablation#5 ablation#6
## cell#1           0  0.0000000  0.0000000 0.00000000  0.0000000          0
## cell#2           0  0.1673226  0.0000000 0.00000000  0.0000000          0
## cell#3           0  0.0000000  0.4819554 0.00000000  0.0000000          0
## cell#4           0  0.0000000  0.0000000 0.73226042  0.0000000          0
## cell#5           0  0.0000000  0.0000000 0.02717766  0.6574428          0
## cell#6           0  0.0000000  0.0000000 0.00000000  0.0000000          0
```

```
## now calculate anticipated signals in ablations
ablations_signals <-
  data.frame(
    ablation = levels(ablations$ablation),
    signal = apply(overlap_matrix, 2, function(abl_i) {
      sum(abl_i * cell_signals$signal)
    })


  )
## and add them to the ablations data.frame
ablations$signal <-
  ablations_signals$signal[match(ablations$ablation, ablations_signals$ablation)]
```

```
## plot signals to check, seems to be ok
simFig2 <-

ggplot() +
  coord_fixed() +
  scale_fill_gradient(low = "white", high = "red") +
  geom_polygon(
    data = cells,
    aes(
```

```
      x = x,
      y = y,
      group = cell
    ),
    color = NA,
    fill = "gray80",
    size = .3
  ) +
   geom_polygon(
    data = ablations,
    aes(
      x = x,
      y = y,
      group = ablation,
      fill = signal
    ),
    color = "black",
      linetype = 2,
      size = .5,
      show.legend = F
  ) +
  theme_void() +
  theme(panel.background = element_rect(fill = "white", color = NA))

simFig2
```

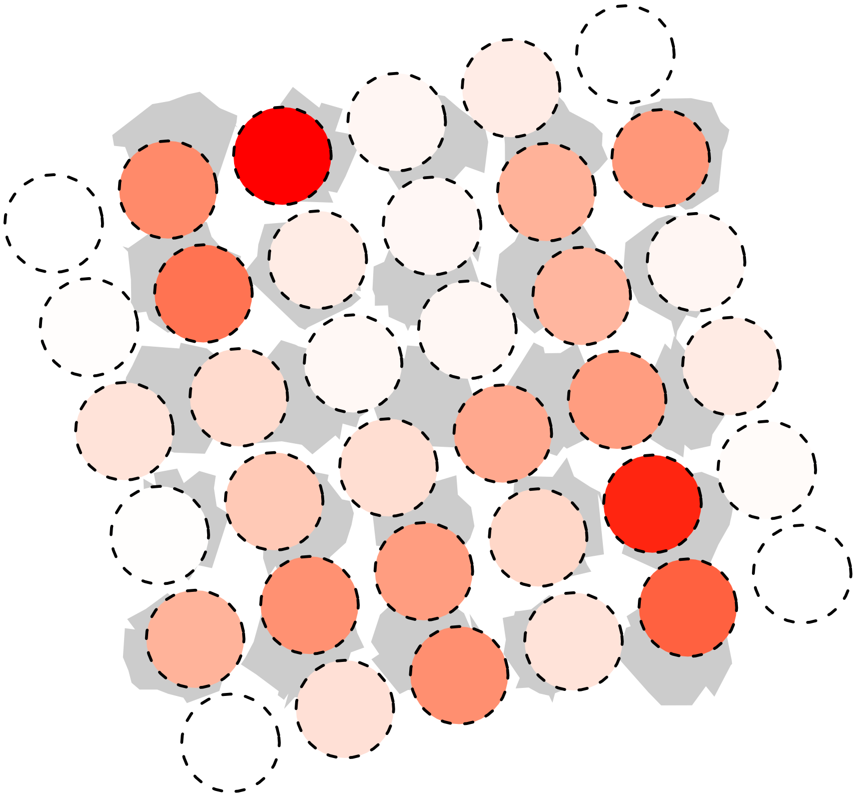

```
## for Luca's normalization method, we also need the 'specificity matrix'
```

```r
specificity_matrix <-
  as.data.frame(t(apply(overlap_matrix, 1, function(cell_i) {
    specificities <- cell_i / sum(cell_i)
    specificities[is.nan(specificities)] <- 0
    return(specificities)
  })))
```

```r
## now calculate cell_signals, based on the AM signals
## this then can be compared with in ground truth (cell_signals$signal)
## and validate the normalization

simulation_comparison <-
list(
  data.frame(
    cell_signals,
    method = "WA",
    computed_signal =
      cell_normalization_Rappez_et_al(
        overlap_proportion_matrix = overlap_matrix,
        overlap_specificity_matrix = specificity_matrix,
        AM_ion_intensity = ablations_signals$signal ,
        AM_proportion_treshold_global = 0,
        AM_specificity_treshold_cell = 0
      )$cell_intensities
  ),
  data.frame(
    cell_signals,
    method = "LIM",
    computed_signal =
      cell_normalization_NNLS(
        overlap_proportion_matrix = overlap_matrix,
        AM_ion_intensity = ablations_signals$signal ,
        AM_proportion_treshold_per_cell = 0.0
      )$cell_intensities

  )
) %>% bind_rows()
```

```r
## we can test different normalization strategies

simFig3 <-
  simulation_comparison  %>%
  {
    ggplot(data = .,
           aes(x = signal,
               y = computed_signal, color = method)) +
      geom_point(size = 3,
                 alpha = .3,
                 show.legend = F) +
      facet_grid(method ~ .) +
      scale_color_paletteer_d("ggthemes::calc") +
      coord_fixed() +
      geom_text(
        data = . %>% group_by(method) %>%
```

```
          summarise(
            r = cor(signal, computed_signal,
                    method = "spearman",
                    use = "pairwise.complete.obs")
          ),
        show.legend = F,
        aes(label = paste0("\n    = ",
                           format(
                             x = r,
                             digits = 3
                           ))),
        size = 3,
        x = -Inf,
        y = Inf,
        hjust = 0,
        vjust = 1,
        color = "gray30"
      ) +
      labs(x = 'ground-truth signal', y = "assigned signal") +
      geom_abline(slope = 1,
                  color = "gray30",
                  linetype = 2) +
      theme_minimal() +
      theme(
        axis.text.x = element_text(size = 9),
        axis.title = element_text(size = 11),
        panel.grid = element_blank(),
        axis.ticks = element_line(),
        panel.background = element_rect(colour = "black", size = 1),
        strip.text = element_text(size = 11, face = "bold")
      )
  }

simFig3
```

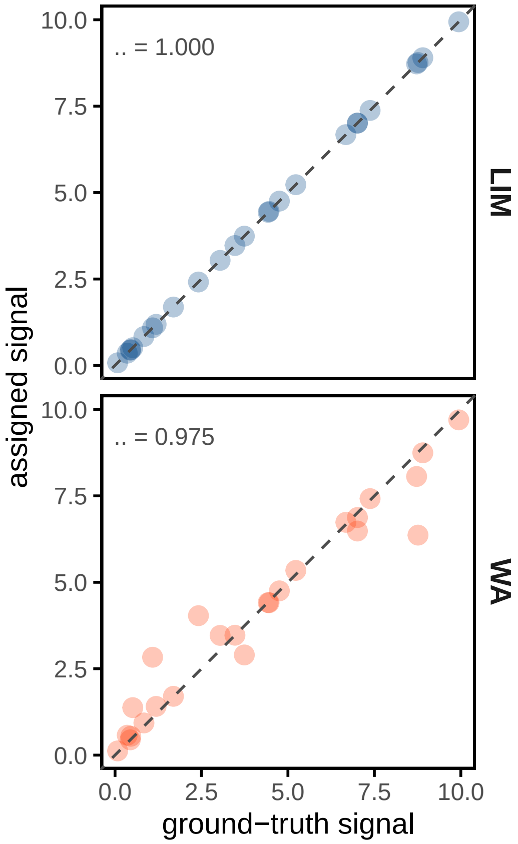


```
cells_per_AM <-
sapply(prop_spec_matrix, function(set_i){

    apply(set_i$overlap_proportion_matrix, 2, function(i){
      sum(i > 0)
    })
}, simplify = F) %>% do.call("c",.)


Fig_number_of_cells <-
ggplot(data = data.frame(table(cells_per_AM[cells_per_AM > 0])) %>%
         mutate(fraction = Freq / sum(Freq),
                Var1 = paste0(Var1, " cells")),
       aes(x = "", y = fraction, fill = Var1)) +
  geom_bar(stat = "identity",
           width = 1, alpha = .7,
           color = "white", show.legend = F) +
  geom_label_repel(data = . %>%
              arrange(fraction) %>%
              mutate(prop = fraction,
                     ypos = cumsum(prop)- 0.5*prop ),
            aes(y = ypos, x = 1.5,label = Var1 ), color = "gray70",
            size = 5, min.segment.length = 0, show.legend = F) +
  coord_polar("y", start = 0) +
  scale_color_paletteer_d("ggthemes::calc") +
  scale_fill_paletteer_d("ggthemes::calc") +
  theme_void() # remove background, grid, numeric labels
```

```r
Fig_number_of_cells
```

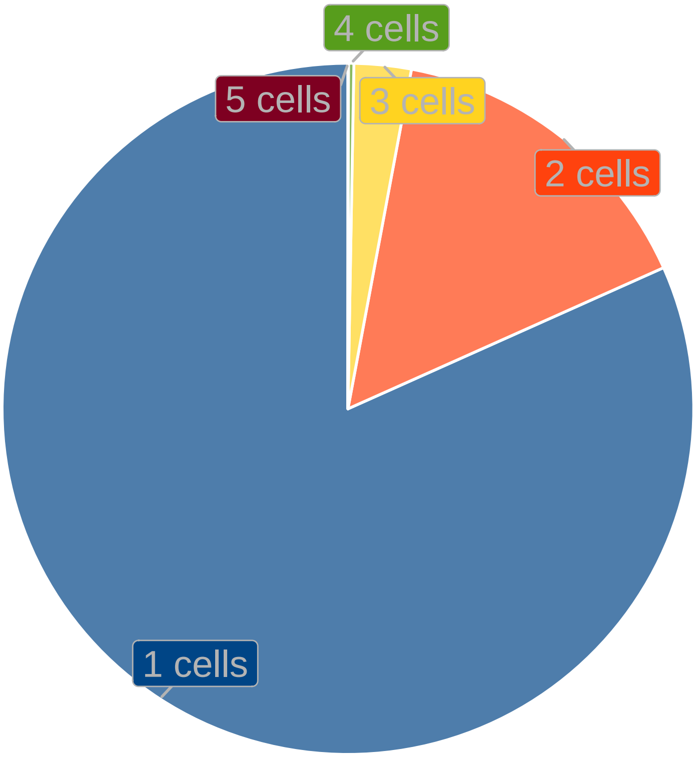


```r
Fig1 <-
plot_grid(
  ncol = 1,
  rel_heights = c(1, 1.5),
  plot_grid(
    labels = c("D", "E"),
    scale = .9,
    Fig1C,
    Fig1D,
    nrow = 1,
    rel_widths = c(2, 1),
    align = "hv",
    axis = "tb"
  ),
  plot_grid(
    scale = .95,
    labels = c("F", "G"),
    #Fig_number_of_cells,
    plot_grid(
      scale = 1,
      simFig1,
      simFig2,
      align = "v",
      ncol = 1
    ),
    simFig3,
```

```
    nrow = 1,
    align = "hv",
    rel_widths = c(1, 1, 2)
  )
)+theme(panel.background = element_rect("white", color = NA))

Fig1
```

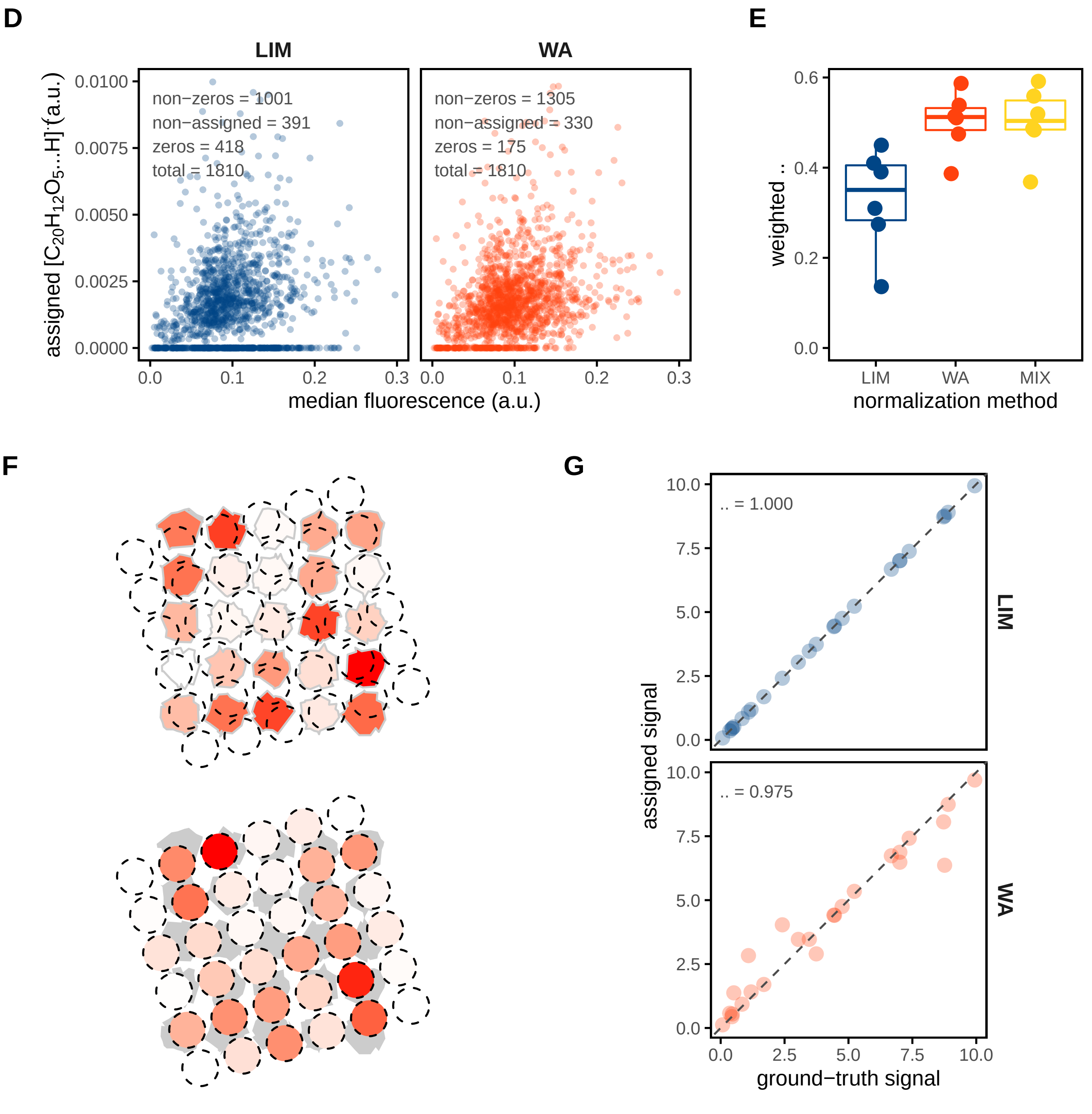


**figure 2**

```
ion_suppression <-
list(
  data_w_ppm %>% filter(
```

```
    file == '2022-02-18_FDA_SpaceM/w7',
    am_sampling_ratio > .1,
    C20H12O5.H_4ppm != 0
  ) %>%
    mutate(
      ratio = C20H12O5.H_4ppm / sum_intensity.FITC,
      ratio = ratio / median(ratio[am_sampling_ratio == 1], na.rm = T),
      normalisation = "none"
    ),
  data_w_ppm %>% filter(
    file == '2022-02-18_FDA_SpaceM/w7',
    am_sampling_ratio > .1,
    C20H12O5.H_4ppm != 0
  ) %>%
    mutate(
      ratio = (C20H12O5.H_4ppm / TIC) / sum_intensity.FITC,
      ratio = ratio / median(ratio[am_sampling_ratio == 1], na.rm = T),
      normalisation = "TIC"
    )
) %>% bind_rows()

Fig2A <-
  ggplot(data = ion_suppression,
         aes(x = am_sampling_ratio,
             y = ratio, color = normalisation)) +
  scale_y_log10(breaks = c(.25, 5, .75, 1, 2.5, 5, 7.5, 10, 25, 50, 75, 100)) +
  scale_x_log10(breaks = seq(0, 1, .2)) +
  geom_point(alpha = .35, size = .7) +
  scale_color_paletteer_d("ggthemes::calc") +
  geom_smooth(method = "lm",
              linetype = 2,
              se = F) +
  geom_hline(yintercept = 1,
             color = "gray30",
             linetype = 2) +
  labs(
    x = "sampling proportion",
    #y = expression("[C"[20]*"H"[12]*"O"[5]*"-H]"^"-"/'sum fluorescence (a.u.)'))+
    y = expression(
      bolditalic( ) ~ "= [C"[20] * "H"[12] * "O"[5] * "-H]" ^ "- " / ' sum fluorescence (a.u.)'
    )
  ) +
  theme_minimal() +
  theme(
    axis.text.x = element_text(size = 9),
    axis.title = element_text(size = 11),
    panel.grid = element_blank(),
    axis.ticks = element_line(),
    panel.background = element_rect(colour = "black", size = 1),
    strip.text = element_text(size = 15, face = "bold")
  )
```

```
Fig2A
```

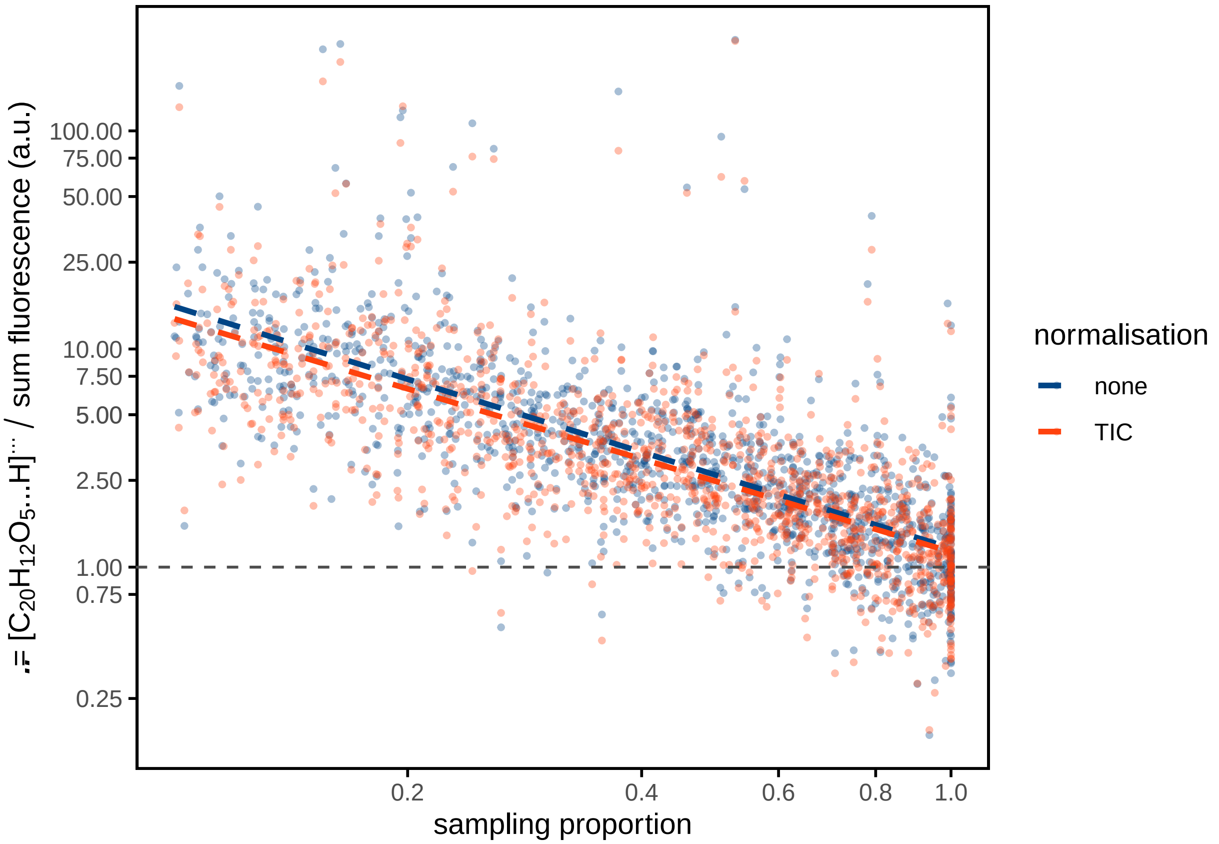


```
library(quantreg)

# model <- quantreg::rq(
#   formula = log(ratio, base = 10) ~ log(am_sampling_ratio, base = 10),
#   tau = .5,
#   data = ion_suppression %>% filter(is.finite(ratio) & ratio > 0 )
# )

model <- quantreg::rq(
  formula = ratio ~ am_sampling_ratio,
  tau = .5,
  data = ion_suppression %>% filter(is.finite(ratio) & ratio > 0 ) %>%
         mutate(ratio = log(ratio,base = 10),
                am_sampling_ratio = log(am_sampling_ratio, base = 10)))
```

```
data_w_ppm$ion_suppr_model <- 10^predict(model,
                                        newdata = data_w_ppm %>%
                                          mutate(am_sampling_ratio = log(am_sampling_ratio, base = 10)))
```

unsupervised model

```
ion_suppression_by_proportion <-
sapply(colnames(data_w_ppm)[grepl("^C",colnames(data_w_ppm))], function(analyte_i){

    analyte <- data_w_ppm %>% filter(
      file == '2022-02-18_FDA_SpaceM/w2',
```

```
    am_sampling_ratio > .1,
  ) %>% .[,analyte_i]

  data_w_ppm_set <-
  data_w_ppm %>% filter(
    file == '2022-02-18_FDA_SpaceM/w2',
    am_sampling_ratio > .1
  ) %>% data.frame

  data_w_ppm_set$ratio <- analyte[[1]] / data_w_ppm_set$am_sampling_ratio

  data_w_ppm_set$ratio <-
    data_w_ppm_set$ratio /
    median(data_w_ppm_set$ratio[data_w_ppm_set$am_sampling_ratio == 1], na.rm = T)


  data_w_ppm_set$analyte <- analyte_i
  data_w_ppm_set[data_w_ppm_set$ratio != 0,]


}, simplify = F)   %>% bind_rows()



model_2 <- quantreg::rq(
  formula = ratio ~ am_sampling_ratio,
  tau = .5,
  data = ion_suppression_by_proportion %>% filter(is.finite(ratio) & ratio > 0 ) %>%
        mutate(ratio = log(ratio,base = 10),
               am_sampling_ratio = log(am_sampling_ratio, base = 10)))


data_w_ppm$ion_suppr_model_unsupervised <-
  10 ^ predict(model_2,
               newdata = data_w_ppm %>%
                 mutate(am_sampling_ratio =
                          log(am_sampling_ratio, base = 10)))
```

```
data_summary <-
list(data_w_ppm %>% group_by(file) %>%
       mutate(C20H12O5.H_4ppm = C20H12O5.H_4ppm) %>%
       summarise(
         r = cor(
           C20H12O5.H_4ppm,
           sum_intensity.FITC,
           method = 'spearman',
           use =  "pairwise.complete.obs"
         ),
          rweight = weightedCorr(
                ## weighted median_intensity.FITC
                x = sum_intensity.FITC,
                y = C20H12O5.H_4ppm,
                method = "Spearman",
                weights = 1 - n_neighbours(x = median_intensity.FITC) /
```

```
                  max(n_neighbours(median_intensity.FITC))
      ),
      normalisation = "none"
    ),
  data_w_ppm %>% group_by(file) %>%
    mutate(C20H12O5.H_4ppm = C20H12O5.H_4ppm / TIC) %>%
    summarise(
      r = cor(
        C20H12O5.H_4ppm,
        sum_intensity.FITC,
        method = 'spearman',
        use =  "pairwise.complete.obs"
      ),

       rweight = weightedCorr(
            ## weighted median_intensity.FITC
            x = sum_intensity.FITC,
            y = C20H12O5.H_4ppm,
            method = "Spearman",
            weights = 1 - n_neighbours(x = median_intensity.FITC) /
                    max(n_neighbours(median_intensity.FITC))
      ),
      normalisation = "TIC"
    ),
  data_w_ppm %>% group_by(file) %>%
    mutate(C20H12O5.H_4ppm = C20H12O5.H_4ppm / ion_suppr_model) %>%
    summarise(
      r = cor(
        C20H12O5.H_4ppm,
        sum_intensity.FITC,
        method = 'spearman',
        use =  "pairwise.complete.obs"
      ),
       rweight = weightedCorr(
            ## weighted median_intensity.FITC
            x = sum_intensity.FITC,
            y = C20H12O5.H_4ppm,
            method = "Spearman",
            weights = 1 - n_neighbours(x = median_intensity.FITC) /
                    max(n_neighbours(median_intensity.FITC))
      ),
      normalisation = "ISM (supervised)"
    ),
   data_w_ppm %>% group_by(file) %>%
    mutate(C20H12O5.H_4ppm = C20H12O5.H_4ppm / ion_suppr_model) %>%
    summarise(
      r = cor(
        C20H12O5.H_4ppm,
        sum_intensity.FITC,
        method = 'spearman',
        use =  "pairwise.complete.obs"
      ),
       rweight = weightedCorr(
```

```
                ## weighted median_intensity.FITC
                x = sum_intensity.FITC,
                y = C20H12O5.H_4ppm,
                method = "Spearman",
                weights = 1 - n_neighbours(x = median_intensity.FITC) /
                        max(n_neighbours(median_intensity.FITC))
         ),
         normalisation = "ISM (non-supervised)"
       )) %>% bind_rows()


data_summary$normalisation <-
  factor(
    data_summary$normalisation,
    levels = c("none", "TIC", "ISM (supervised)", "ISM (non-supervised)")
  )
```

```
Fig2B <-
  data_summary %>%  {
    ggplot(data = .,
           aes(x = normalisation,
               y = rweight, color = normalisation)) +
      #geom_vline(xintercept = 3, linetype = 3, color = "gray30")+
      scale_y_continuous(limits = c(0, NA)) +
      geom_boxplot(outlier.colour = NA, show.legend = F) +
      geom_jitter(size = 3,
                  width = .08,
                  show.legend = F) +
      scale_color_paletteer_d("ggthemes::calc") +
      #geom_line(aes(group = file, color = file), linetype = 2, alpha = .7)+
      labs(y = 'weighted  ',  x = "", color = "normalisation") +
      theme_minimal() +
      theme(
        axis.text.x = element_blank(),
        axis.title = element_text(size = 11),
        panel.grid = element_blank(),
        axis.ticks = element_line(),
        panel.background = element_rect(colour = "black", size = 1)
      )


  }

Fig2B
```

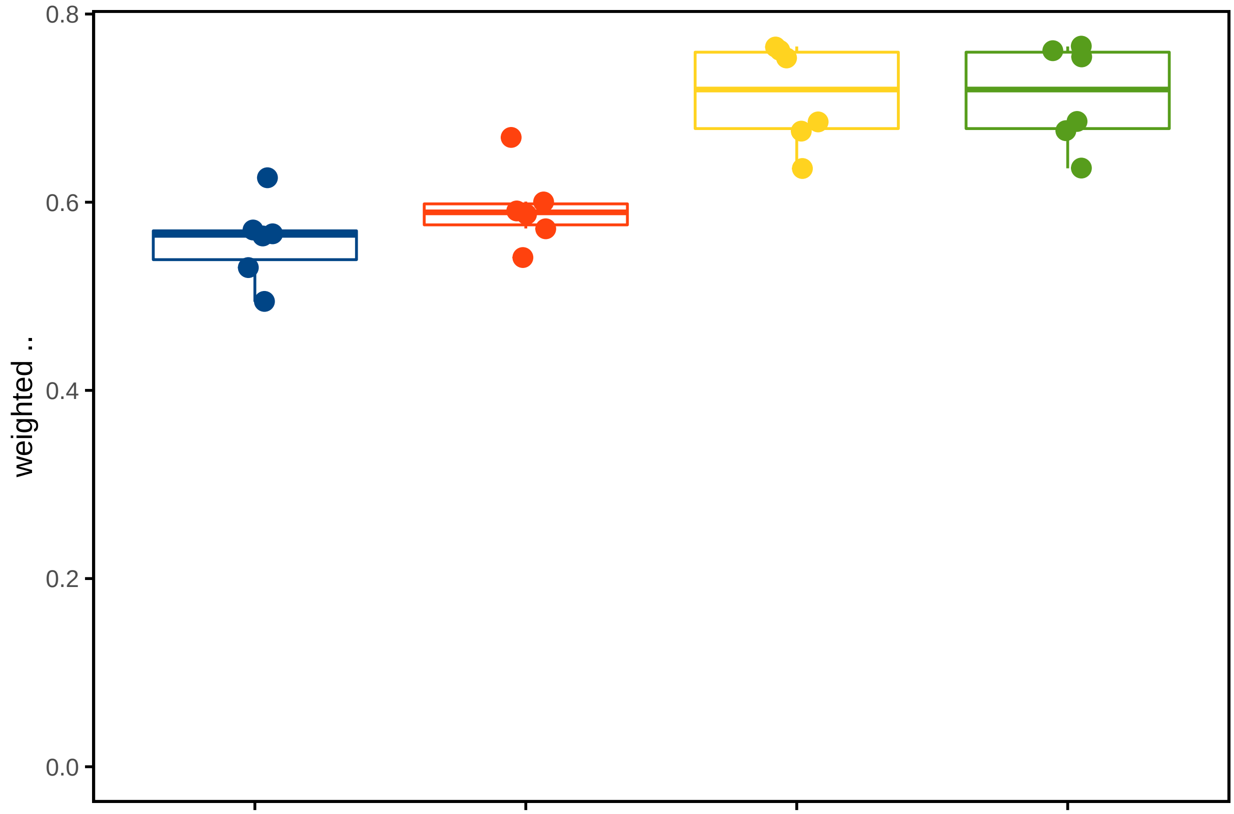


```
datasets_names <- data_w_ppm %>% group_by(file) %>% summarize(file = file[1]) %>% pull(file)

cells <-
  sapply(datasets_names, function(dataset_i) {
    ## in every iteration, different normalizations are assessed

    ## first, retreive prop and spec matrix for dataset_i
    prop_spec_matrix_OI <-
      prop_spec_matrix[[paste0(dataset_i, '/')]]


    ## now, the normalization parameters
    list({
      ## Rappez_et_al method; no TIC-norm, zeros included

      AM_ion_intensity_OI <-
        data_w_ppm %>% filter(file ==  dataset_i) %>%         ## filter dataset of interest
        mutate(C20H12O5.H = C20H12O5.H_4ppm) %>%              ## don't normalize C20H12O5.H
        pull(C20H12O5.H)

      cell_ion_intensity <-
        cell_normalization_Rappez_et_al(
          overlap_proportion_matrix = prop_spec_matrix_OI$overlap_proportion_matrix,
          overlap_specificity_matrix = prop_spec_matrix_OI$overlap_specificity_matrix,
          AM_ion_intensity = AM_ion_intensity_OI,
          AM_proportion_treshold_global = 0.3,
          skip_AM_zeros = FALSE                             ## don't omit zeros in AM vector
        )$cell_intensities
```

```
    data.frame(cell_id = rownames(cell_ion_intensity),
               C20H12O5.H_calculated = cell_ion_intensity,
               method = 'weighted average',
               AM_normalization = 'none',
               AM_filter = '0.3',
               AM_omit_zeros = "keep zeros",
               file = dataset_i)
  },
  {
    ## Rappez_et_al method;  TIC-norm, zeros included

    AM_ion_intensity_OI <-
      data_w_ppm %>% filter(file ==  dataset_i) %>%              ## filter dataset of interest
      mutate(C20H12O5.H = C20H12O5.H_4ppm / TIC) %>%             ## don't normalize C20H12O5.H
      pull(C20H12O5.H)

    cell_ion_intensity <-
      cell_normalization_Rappez_et_al(
        overlap_proportion_matrix = prop_spec_matrix_OI$overlap_proportion_matrix,
        overlap_specificity_matrix = prop_spec_matrix_OI$overlap_specificity_matrix,
        AM_ion_intensity = AM_ion_intensity_OI,
        AM_proportion_treshold_global = 0.3,
        skip_AM_zeros = FALSE                              ## don't omit zeros in AM vector
      )$cell_intensities

    data.frame(cell_id = rownames(cell_ion_intensity),
               C20H12O5.H_calculated = cell_ion_intensity,
               method = 'weighted average',
               AM_normalization = 'TIC',
               AM_filter = '0.3',
               AM_omit_zeros = "keep zeros",
               file = dataset_i)
  },
  {
    ## Rappez_et_al method; suppr model norm, zeros included

    AM_ion_intensity_OI <-
      data_w_ppm %>% filter(file ==  dataset_i) %>%               ## filter dataset of interest
      mutate(C20H12O5.H = C20H12O5.H_4ppm / ion_suppr_model) %>%  ## don't normalize C20H12O5.H
      pull(C20H12O5.H)

    cell_ion_intensity <-
      cell_normalization_Rappez_et_al(
        overlap_proportion_matrix = prop_spec_matrix_OI$overlap_proportion_matrix,
        overlap_specificity_matrix = prop_spec_matrix_OI$overlap_specificity_matrix,
        AM_ion_intensity = AM_ion_intensity_OI,
        AM_proportion_treshold_global = 0.3,
        skip_AM_zeros = FALSE                                   ## don't omit zeros in AM vector
      )$cell_intensities

    data.frame(cell_id = rownames(cell_ion_intensity),
               C20H12O5.H_calculated = cell_ion_intensity,
               method = 'weighted average',
```

```
                AM_normalization = 'ISM (supervised)',
                AM_filter = '0.3',
                AM_omit_zeros = "keep zeros",
                file = dataset_i)
    },
    {
      ## Rappez_et_al method; suppr model norm, zeros included

      AM_ion_intensity_OI <-
        data_w_ppm %>% filter(file ==  dataset_i) %>%                ## filter dataset of interest
        mutate(C20H12O5.H = C20H12O5.H_4ppm / ion_suppr_model_unsupervised) %>%
        ## don't normalize C20H12O5.H
        pull(C20H12O5.H)

      cell_ion_intensity <-
        cell_normalization_Rappez_et_al(
          overlap_proportion_matrix = prop_spec_matrix_OI$overlap_proportion_matrix,
          overlap_specificity_matrix = prop_spec_matrix_OI$overlap_specificity_matrix,
          AM_ion_intensity = AM_ion_intensity_OI,
          AM_proportion_treshold_global = 0.3,
          skip_AM_zeros = FALSE                     ## don't omit zeros in AM vector
        )$cell_intensities

      data.frame(cell_id = rownames(cell_ion_intensity),
                C20H12O5.H_calculated = cell_ion_intensity,
                method = 'weighted average',
                AM_normalization = 'ISM (non-supervised)',
                AM_filter = '0.3',
                AM_omit_zeros = "keep zeros",
                file = dataset_i)
    })  %>% bind_rows()

  }, simplify = FALSE) %>% bind_rows()
```

```
cells <-
  cells %>% left_join(
    cells_SpaceM %>% select(file, cell_id, median_intensity.FITC, cell_area, eccentricity),
    by = c("file", "cell_id")
  )

cells <-  cells %>% group_by(method, AM_normalization,AM_filter,AM_omit_zeros,file) %>%
  mutate(n = length(C20H12O5.H_calculated))



cell_summary <-
  cells %>%
  group_by(method, AM_normalization, AM_filter, AM_omit_zeros, file) %>%
  filter(          ## filter out NAs for the correlations
    !is.na(median_intensity.FITC) &!is.na(C20H12O5.H_calculated)) %>%
  summarize(
      r = cor(
      x = median_intensity.FITC,
      y = C20H12O5.H_calculated,
```

```
      method = "spearman",
      use = "pairwise.complete.obs"
    ),
    rweight = weightedCorr(
      ## weighted median_intensity.FITC
      x = median_intensity.FITC,
      y = C20H12O5.H_calculated,
      method = "Spearman",
      weights = 1 - n_neighbours(x = median_intensity.FITC) /
        max(n_neighbours(median_intensity.FITC))
    ),
    n = n[1]
  )



cell_summary$AM_normalization <-
  factor(
    cell_summary$AM_normalization,
    levels = c("none", "TIC", "ISM (supervised)", "ISM (non-supervised)")
  )
```

```
Fig2C <-
cell_summary %>%  {
  ggplot(data = .,
         aes(x = AM_normalization,
             y = rweight, color = AM_normalization))+
    #geom_vline(xintercept = 3, linetype = 3, color = "gray30")+
    scale_y_continuous(limits = c(0,NA))+
    geom_boxplot(outlier.colour = NA, show.legend = F)+
    geom_jitter(size = 3, width = .08, show.legend = T)+
    scale_color_paletteer_d("ggthemes::calc")+
    #geom_line(aes(group = file, color = file), linetype = 2, alpha = .7)+
    labs(y = 'weighted  ',  x = "", color = "normalisation")+
    theme_minimal()+
    theme(axis.text.x = element_blank(),
          axis.title = element_text(size = 11), panel.grid = element_blank(),
          axis.ticks = element_line(),
          panel.background = element_rect(colour = "black", size = 1))
}

Fig2C
```

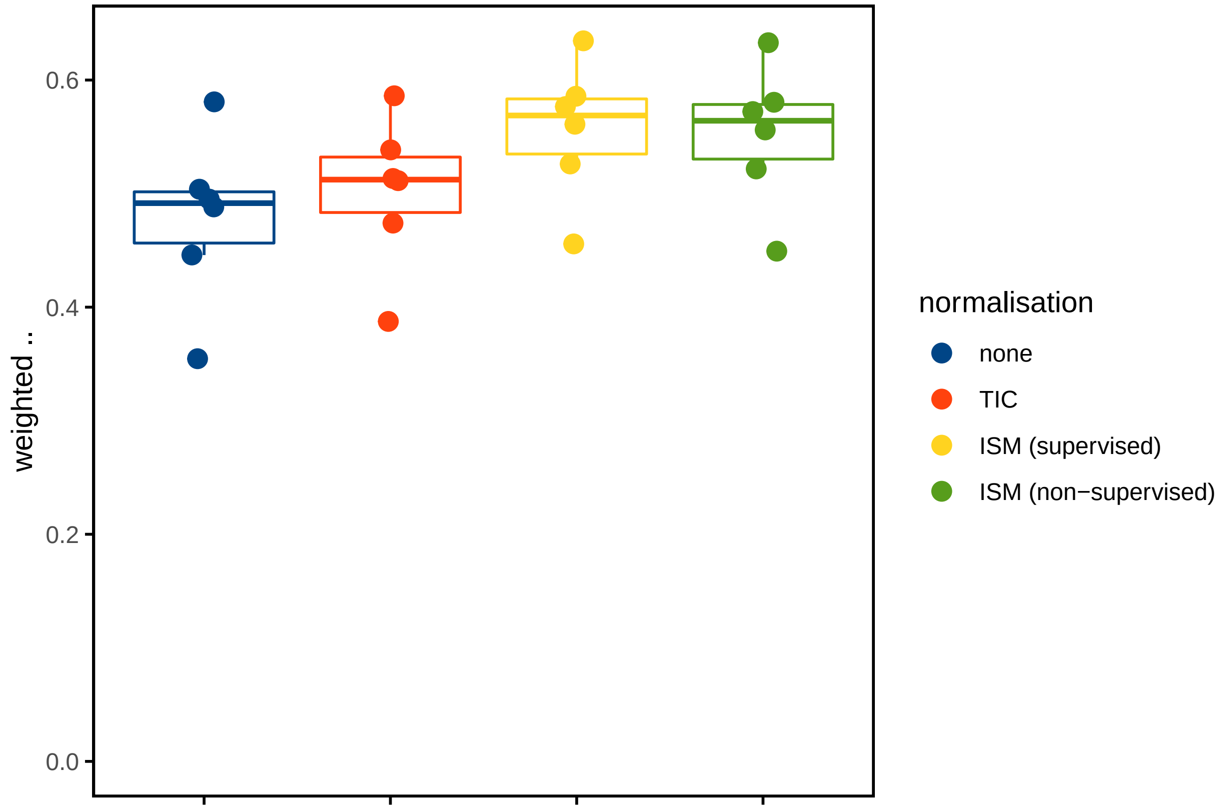


```
ion_suppression_by_proportion_viz <-
ion_suppression_by_proportion %>% group_by(analyte) %>%
  mutate(keep = mean(!(is.na(ratio) | is.infinite(ratio))) > .3) %>%
  filter(keep) %>%
  mutate(analyte = paste0("[",gsub("\\.","-",analyte),"]-"),
         analyte = gsub("_4ppm","",analyte))

ion_suppression_by_proportion_viz$analyte <-
  factor(ion_suppression_by_proportion_viz$analyte)
ion_suppression_by_proportion_viz$analyte <-
  factor(
    ion_suppression_by_proportion_viz$analyte,
    levels = levels(ion_suppression_by_proportion_viz$analyte)[c(3, 1, 2, 4)]
  )

Fig2D <-
  ion_suppression_by_proportion_viz %>%
  ggplot(data = .,
         aes(x = am_sampling_ratio,
             y = ratio)) +
  scale_y_log10(breaks = c(.25, 5, .75, 1, 2.5, 5, 7.5, 10, 25, 50, 75, 100)) +
  scale_x_log10(breaks = seq(0, 1, .2)) +
  facet_grid(analyte ~ .) +
  geom_point(alpha = .35,
             size = .7,
             color = palettes_d$ggthemes$calc[4]) +
  scale_color_paletteer_d("ggthemes::calc") +
  geom_smooth(
```

```
    method = "lm",
    linetype = 2,
    se = F,
    color = "gray30"
  ) +
  geom_hline(yintercept = 1,
             color = "gray30",
             linetype = 2) +
  labs(
    x = "sampling proportion",
    #y = paste0('MS-signal / sampling proportion (a.u.)'))+
    y = expression(bolditalic( ) ~ '=' ~ 'MS-signal ' / ' sampling proportion (a.u.)')
  ) +
  theme_minimal() +
  theme(
    axis.text.x = element_text(size = 9),
    axis.title = element_text(size = 11),
    panel.grid = element_blank(),
    axis.ticks = element_line(),
    panel.background = element_rect(colour = "black", size = 1),
    strip.text = element_text(size = 9, face = "bold")
  )

Fig2D
```

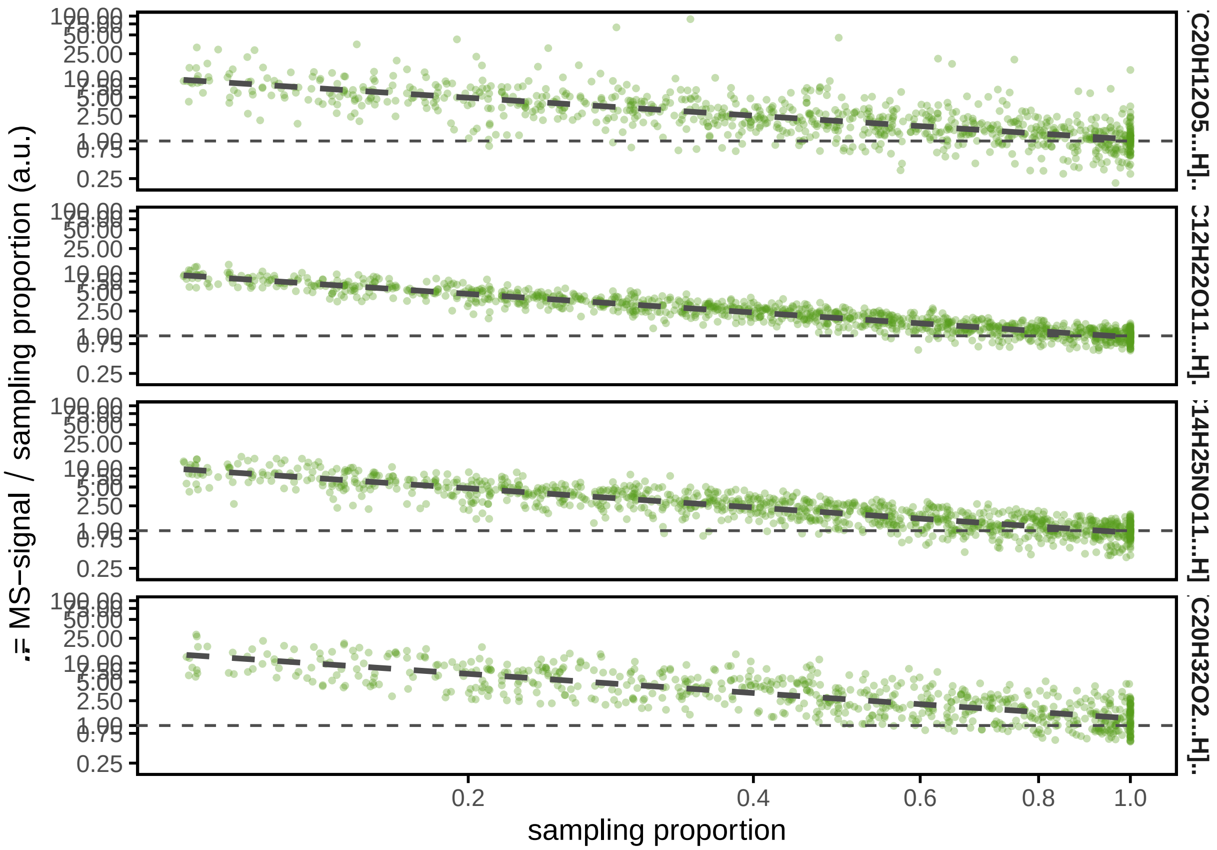


```
Fig2 <-
  cowplot::plot_grid(
```

```
  cowplot::plot_grid(
    #align = "v", axis = "l",

    Fig2A,
    ncol = 1,
    scale = .95,
    plot_grid(
      labels = c("B", "C"), label_size = 18,
      Fig2B,
      Fig2C + labs(y = ""),

      nrow = 1,
      rel_widths = c(1, 2)
    )
  ),
  labels = c("A", "D"), label_size = 18,
  Fig2D,
  scale = .95,
  nrow = 1,
  rel_widths = c(2, 1)
)+ theme(panel.background = element_rect(color = NA, fill = "white"))

Fig2
```

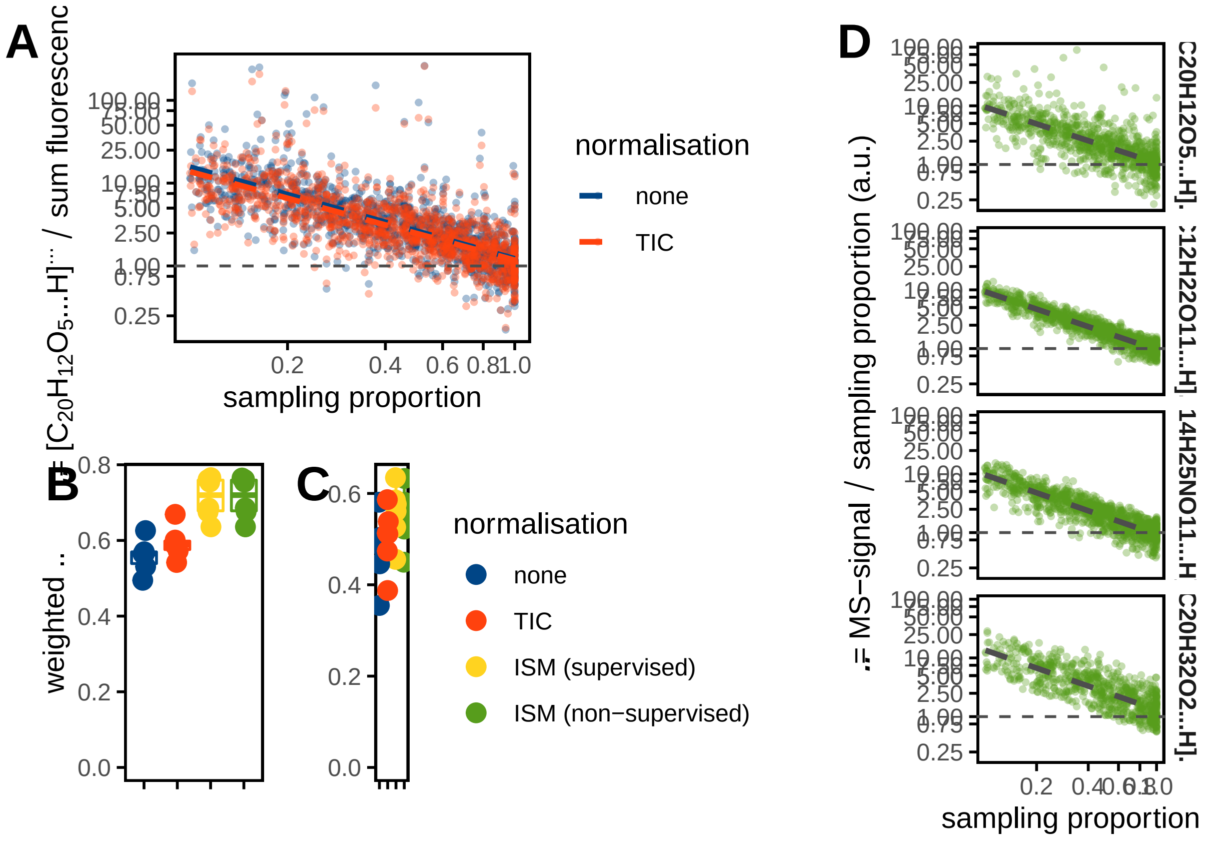

#### Fig 3

```
Fig3A <-
  data_w_ppm %>% filter(file == '2022-02-18_FDA_SpaceM/w4') %>%
  mutate(class = ifelse(C20H12O5.H_4ppm == 0, "zero", "non-zero")) %>%
  ggplot(data = .,
         aes(x = sum_intensity.FITC, color = class,
             y = C20H12O5.H_4ppm)) +
  scale_y_log10(limits = c(NA, 3000)) +
  scale_x_log10() +
  geom_point(alpha = .15,
             size = 2,
             show.legend = T) +
  scale_color_paletteer_d("ggthemes::calc", guide = guide_legend(override.aes = list(alpha = 1))) +
  labs(
    x = 'sum fluorescence (a.u.)',
    y = expression("[C"[20] * "H"[12] * "O"[5] * "-H]" ^ "-" ~ (a.u.)),
    color = expression(identity ~ "[C"[20] * "H"[12] * "O"[5] * "-H]" ^
                         "-")
  ) +
  geom_text(
    data = .  %>%
      filter(## filter out NAs for the correlations!is.na(sum_intensity.FITC) &
        !is.na(sum_intensity.FITC)) %>%
      summarize(
        rweight = weightedCorr(
          ## weighted correlation
          x = sum_intensity.FITC,
          y = C20H12O5.H_4ppm,
          method = "Spearman",
          weights = 1 - n_neighbours(x = sum_intensity.FITC) /
            max(n_neighbours(sum_intensity.FITC))
        ),
        label  = paste0("    weighted   = ",
                        format(rweight, digits = 3), '\n')
      ),
    size = 3,
    x = -Inf,
    y = -Inf,
    hjust = 0,
    vjust = 0,
    color = "gray30",
    aes(label = label)
  ) +
  geom_smooth(
    data = . %>%
      mutate(C20H12O5.H_4ppm = C20H12O5.H_4ppm + min(C20H12O5.H_4ppm[C20H12O5.H_4ppm > 0])),
    ## otherwise zeros are not included in log-scale
    se = F,
    method = "lm",
    color = "gray30",
    linetype = 2
  ) +
  theme_minimal() +
```

```
  theme(
    axis.text.x = element_blank(),
    axis.title.x = element_blank(),
    legend.position = c(0.05, 0.8),
    legend.justification = 0,
    legend.background = element_rect(color = NA, fill = NA),
    legend.title = element_text(size = 8),
    legend.text = element_text(size = 6),
    axis.title = element_text(size = 11),
    panel.grid = element_blank(),
    axis.ticks = element_line(),
    panel.background = element_rect(colour = "black", size = 1),
    strip.text = element_text(face = "bold")
  )

Fig3A
```

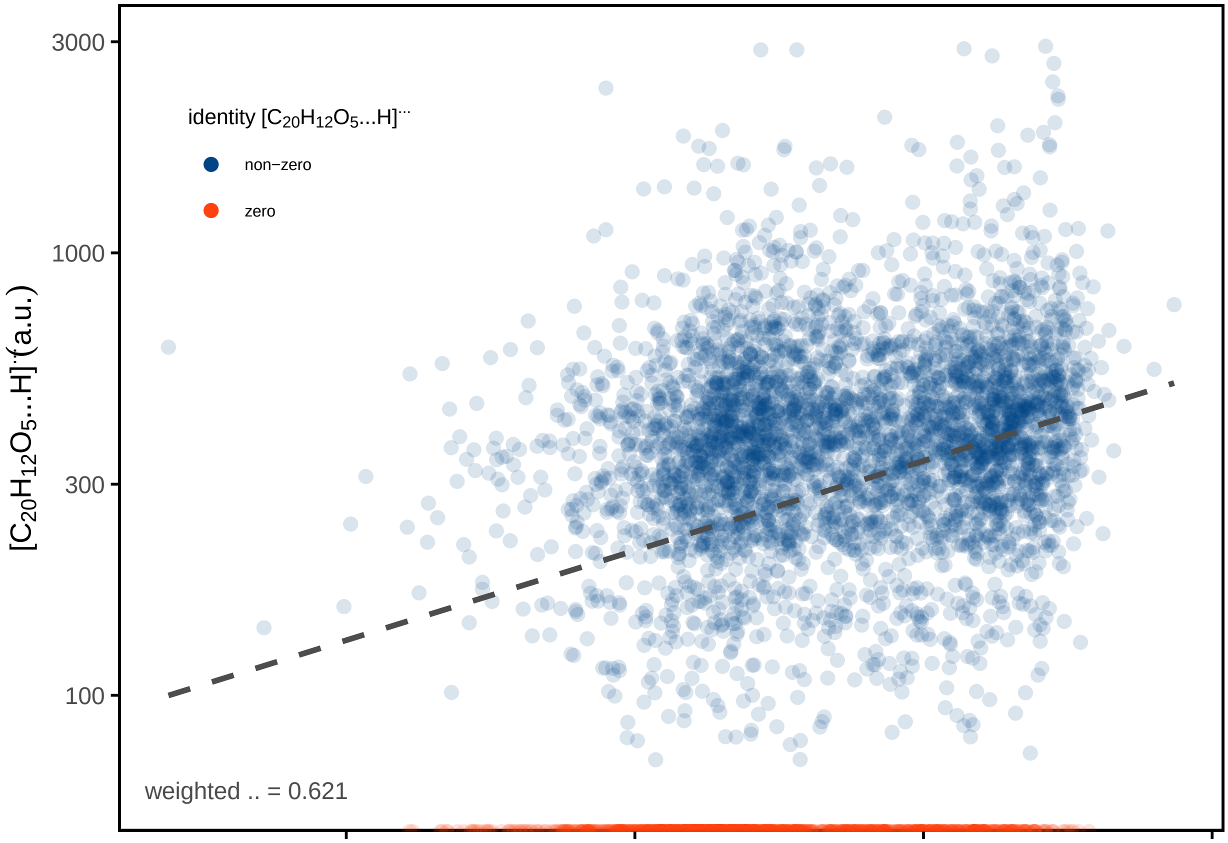


```
Fig3B <-
  data_w_ppm %>% filter(file == '2022-02-18_FDA_SpaceM/w4') %>%
  mutate(class = ifelse(C20H12O5.H_4ppm == 0, "zero", "non-zero")) %>%
  ggplot(data = ., aes(x = sum_intensity.FITC, fill = class)) +
  geom_histogram(show.legend = F) +
  labs(
    y = 'number',
    x = "sum fluorescence (a.u.)",
    fill = expression(identity ~ "[C"[20] * "H"[12] * "O"[5] * "-H]" ^
                        "-")
```

```
  ) +
  scale_x_log10() +
  scale_fill_paletteer_d("ggthemes::calc") +
  theme_minimal() +
  theme(
    axis.text.x = element_text(size = 9),
    legend.position = c(0.05, 0.8),
    legend.justification = 0,
    legend.background = element_rect(color = "black"),
    axis.title = element_text(size = 11),
    panel.grid = element_blank(),
    axis.ticks = element_line(),
    panel.background = element_rect(colour = "black", size = 1)
  )

Fig3B
```

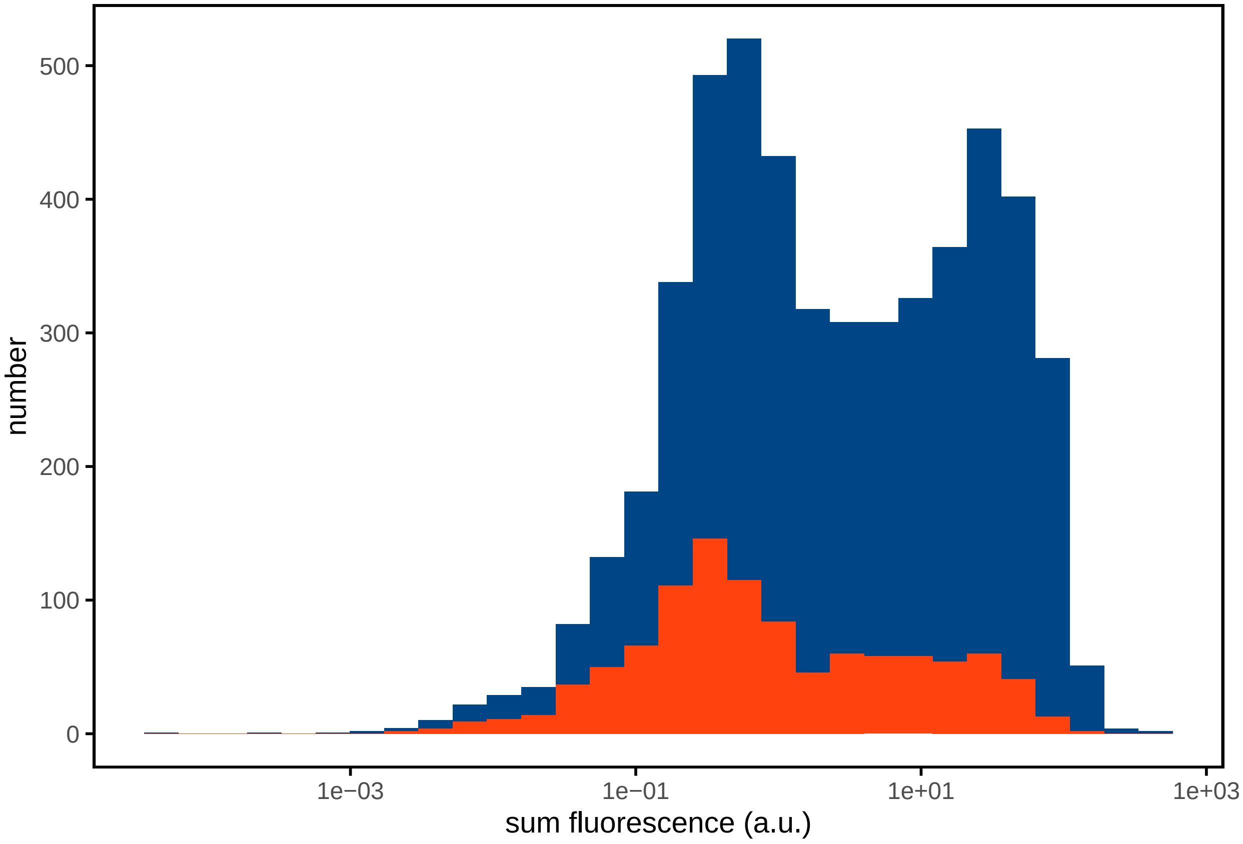


```
Fig3C <-
list(data_w_ppm %>% mutate(C20H12O5.H = C20H12O5.H_4ppm / TIC )  %>%
       group_by(file) %>%
       summarize(r = cor(sum_intensity.FITC,
                         C20H12O5.H, method = "spearman", use = "pairwise.complete.obs"),
                 rweight = weightedCorr(                         ## weighted correlation
                   x = sum_intensity.FITC,
                   y = C20H12O5.H,
                   method = "Spearman",
                   weights = 1 - n_neighbours(x = sum_intensity.FITC) /
```

```
                     max(n_neighbours(sum_intensity.FITC))
                 ),
                 n = sum(!is.na(C20H12O5.H) & !is.na(sum_intensity.FITC)),
                 label  = "all values"),
      data_w_ppm %>% filter(C20H12O5.H_4ppm != 0) %>%
       mutate(C20H12O5.H = C20H12O5.H_4ppm / TIC)  %>%
       group_by(file) %>%
       summarize(r = cor(sum_intensity.FITC,
                         C20H12O5.H, method = "spearman", use = "pairwise.complete.obs"),
                 rweight = weightedCorr(                          ## weighted correlation
                   x = sum_intensity.FITC,
                   y = C20H12O5.H,
                   method = "Spearman",
                   weights = 1 - n_neighbours(x = sum_intensity.FITC) /
                     max(n_neighbours(sum_intensity.FITC))
                 ),
                 n = sum(!is.na(C20H12O5.H) & !is.na(sum_intensity.FITC)),
                 label  = "omit zeros")
) %>% bind_rows %>% {
  ggplot(data = .,
         aes(x = label,
             y = rweight, color = label)) +
    geom_boxplot(show.legend = F) +
    geom_jitter(size = 3,
                width = .2,
                show.legend = F) +
    scale_y_continuous(limits = c(0, .75)) +
    scale_color_paletteer_d("ggthemes::calc",
                            guide = guide_legend(override.aes = list(alpha = 1))) +
    labs(y = 'weighted  ',  x = "", color = "AM filter") +
    theme_minimal() +
    theme(
      axis.text.x = element_blank(),
      axis.title = element_text(size = 11),
      panel.grid = element_blank(),
      axis.ticks = element_line(),
      panel.background = element_rect(colour = "black", size = 1)
    )
}

Fig3C
```

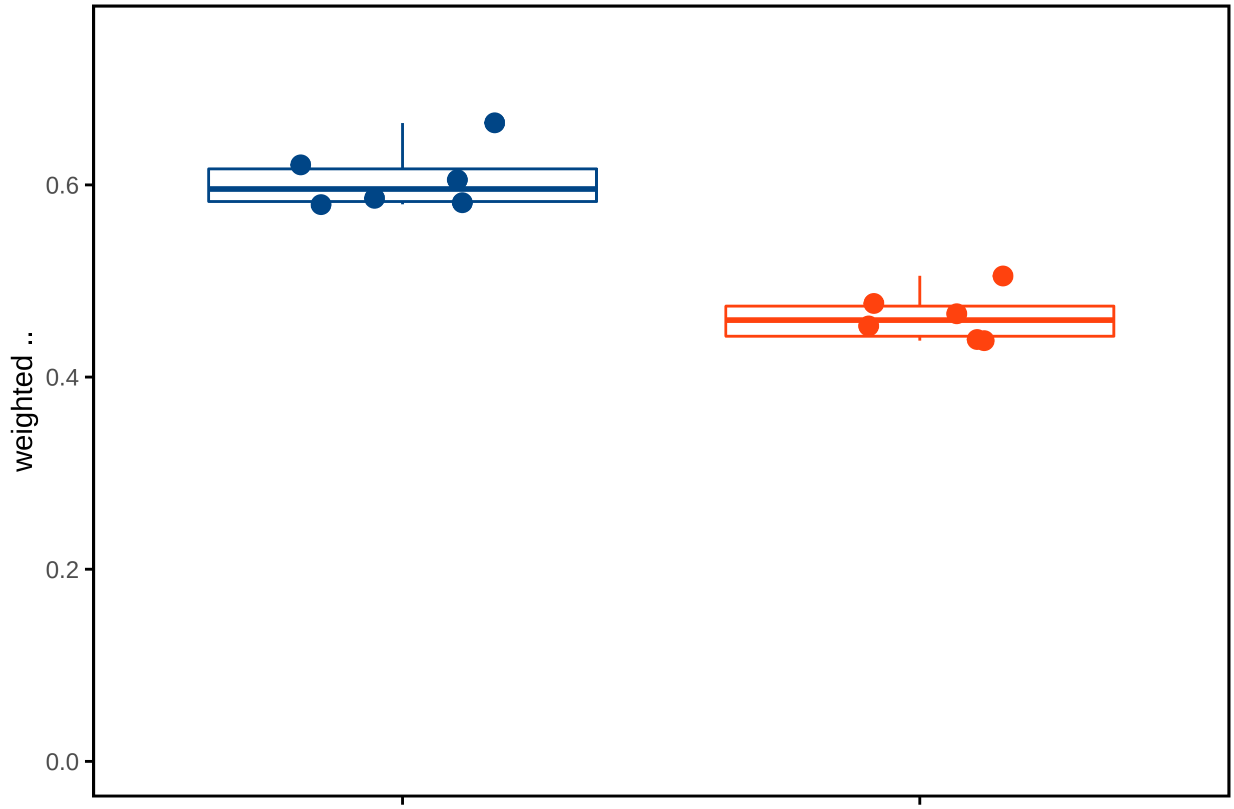


#### by cells

```
datasets_names <- data_w_ppm %>% group_by(file) %>% summarize(file = file[1]) %>% pull(file)

## iterate over:
## datasets_names

cells <-
  sapply(datasets_names, function(dataset_i) {
    ## in every iteration, different normalizations are assessed

    ## first, retreive prop and spec matrix for dataset_i
    prop_spec_matrix_OI <-
      prop_spec_matrix[[paste0(dataset_i, '/')]]


    ## now, the normalization parameters
    list({
      ## Rappez_et_al method; TIC-norm, zeros included

      AM_ion_intensity_OI <-
        data_w_ppm %>% filter(file ==  dataset_i) %>%               ## filter dataset of interest
        mutate(C20H12O5.H = C20H12O5.H_4ppm / TIC) %>%              ## normalize C20H12O5.H  by TIC
        pull(C20H12O5.H)

      cell_ion_intensity <-
        cell_normalization_Rappez_et_al(
          overlap_proportion_matrix = prop_spec_matrix_OI$overlap_proportion_matrix,
          overlap_specificity_matrix = prop_spec_matrix_OI$overlap_specificity_matrix,
```

```
        AM_ion_intensity = AM_ion_intensity_OI,
        AM_proportion_treshold_global = 0.3,
        skip_AM_zeros = FALSE                             ## keep zeros in AM vector
      )$cell_intensities

    data.frame(cell_id = rownames(cell_ion_intensity),
               C20H12O5.H_calculated = cell_ion_intensity,
               method = 'weighted average',
               AM_normalization = 'TIC',
               AM_filter = '0.3',
               AM_omit_zeros = "all values",
               file = dataset_i)
  },{
    ## Rappez_et_al method; TIC-norm, zeros included

    AM_ion_intensity_OI <-
      data_w_ppm %>% filter(file ==  dataset_i) %>%           ## filter dataset of interest

      mutate(C20H12O5.H = C20H12O5.H_4ppm / TIC) %>%          ## normalize C20H12O5.H  by TIC
      pull(C20H12O5.H)

    cell_ion_intensity <-
      cell_normalization_Rappez_et_al(
        overlap_proportion_matrix = prop_spec_matrix_OI$overlap_proportion_matrix,
        overlap_specificity_matrix = prop_spec_matrix_OI$overlap_specificity_matrix,
        AM_ion_intensity = AM_ion_intensity_OI,
        AM_proportion_treshold_global = 0.3,
        skip_AM_zeros = TRUE                               ## keep zeros in AM vector
      )$cell_intensities

    data.frame(cell_id = rownames(cell_ion_intensity),
               C20H12O5.H_calculated = cell_ion_intensity,
               method = 'weighted average',
               AM_normalization = 'TIC',
               AM_filter = '0.3',
               AM_omit_zeros = "omit zeros",
               file = dataset_i)
  })  %>% bind_rows()

 }, simplify = FALSE) %>% bind_rows()
```

```
cells <-
  cells %>% left_join(
    cells_SpaceM %>% select(file, cell_id, median_intensity.FITC, cell_area, eccentricity),
    by = c("file", "cell_id")
  )

cells <-  cells %>% group_by(method, AM_normalization,AM_filter,AM_omit_zeros,file) %>%
  mutate(n = length(C20H12O5.H_calculated))

cell_summary <-
  left_join(
    x = cells,
```

```
    y = data_w_ppm %>% group_by(file) %>%

      summarize(
        signal_C20H12O5.H = quantile(x = C20H12O5.H, probs = 1),
        signal_TIC = quantile(x = TIC, probs = 1),
        signal_fluorescein = quantile(x = median_intensity.FITC, probs = 1),
        AM_area = mean(am_area)
      ),
    by = 'file'
  ) %>%
  group_by(method, AM_normalization, AM_filter, AM_omit_zeros, file) %>%
  filter(          ## filter out NAs for the correlations
    !is.na(median_intensity.FITC) &!is.na(C20H12O5.H_calculated)) %>%
  summarize(
    signal_C20H12O5.H = signal_C20H12O5.H[1],
    signal_TIC = signal_TIC[1],
    signal_fluorescein = signal_fluorescein[1],
    AM_area = AM_area[1],
    r = cor(
      x = median_intensity.FITC,
      y = C20H12O5.H_calculated,
      method = "spearman",
      use = "pairwise.complete.obs"
    ),
    rweight = weightedCorr(
      ## weighted median_intensity.FITC
      x = median_intensity.FITC,
      y = C20H12O5.H_calculated,
      method = "Spearman",
      weights = 1 - n_neighbours(x = median_intensity.FITC) /
        max(n_neighbours(median_intensity.FITC))
    ),
    n = n[1]
  )
```

```
Fig3D <-
cell_summary  %>% {
  ggplot(data = .,
         aes(x = AM_omit_zeros,
             y = rweight, color = AM_omit_zeros)) +
    scale_y_continuous(limits = c(0, NA)) +
    geom_boxplot(outlier.colour = NA, show.legend = F) +
    geom_jitter(size = 3,
                width = .08,
                show.legend = T) +
    scale_color_paletteer_d("ggthemes::calc") +
    labs(y = 'weighted  ',  x = "normalization method") +
    labs(y = 'weighted  ',  x = "", color = "") +
    theme_minimal() +
    theme(
      axis.text.x = element_blank(),
      axis.title = element_text(size = 11),
      panel.grid = element_blank(),
      axis.ticks = element_line(),
```

```
      panel.background = element_rect(colour = "black", size = 1)
    )
}

Fig3D
```

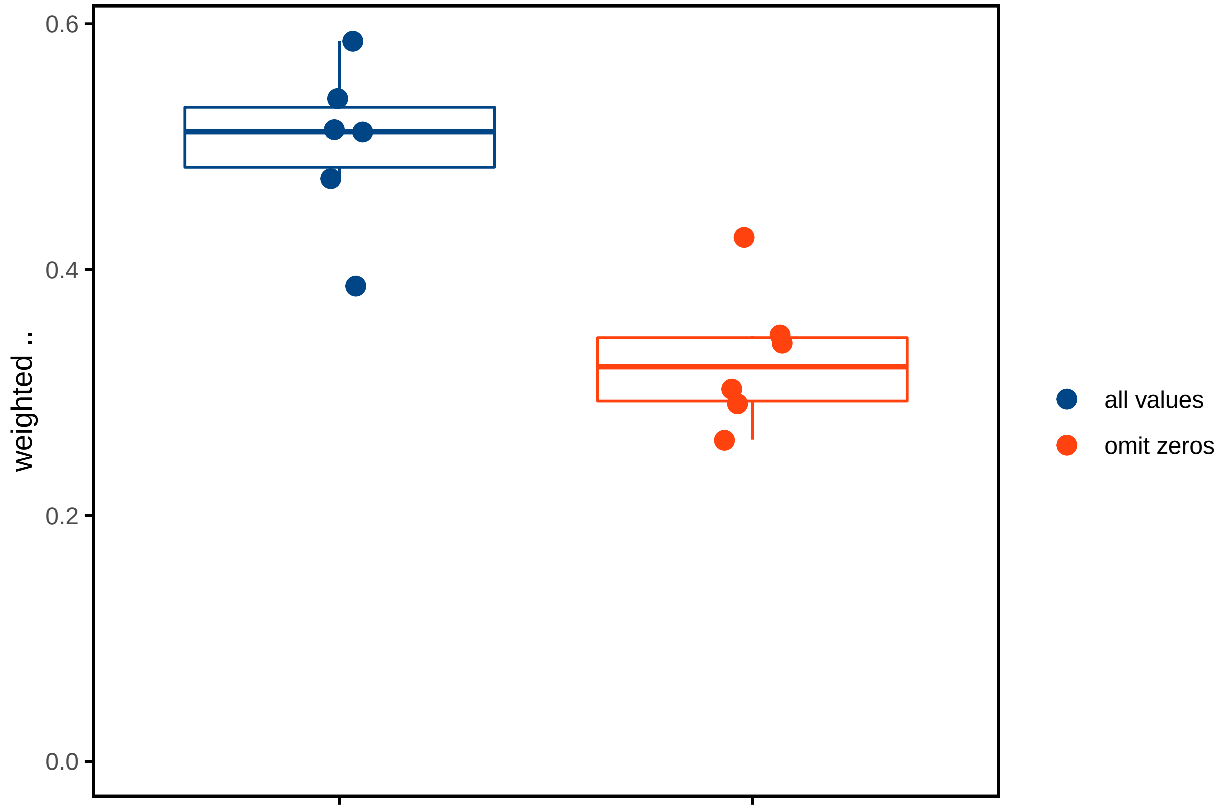


mass tolerances

```
## RAW file of interest
rawfile1 <- 'FDA_raw_files/2021-11-25_FDA_W2_100x100_a25ss30rf100_DANneg.RAW'
rawfile2 <- 'FDA_raw_files/2021-11-25_FDA_W345678_100x100_a25ss30rf100_DANneg.RAW'

## m/z of target ions
mass_target <- c(331.0612)
#tolerance_i <- NULL

data_w_ppm_ranges <-
  pbapply::pbsapply(seq(0.5,20, 1), function(tolerance_i){
    list({
      RawData <- readFileHeader(rawfile = rawfile2)
      FLU_signal <-
        readChromatogram(
          rawfile = rawfile2,
          mass = mass_target,
          tol = tolerance_i,
          type = "xic"
        )

      data_w_ppm %>% filter(file %in% paste0("2022-02-18_FDA_SpaceM/w", 3:7)) %>%
```

```
      mutate(C20H12O5.H_xppm = FLU_signal[[1]]$intensities[1:50000],
             ppm_tolerance = tolerance_i / 2)
    }, {
      RawData <- readFileHeader(rawfile = rawfile1)
      FLU_signal <-
        readChromatogram(
          rawfile = rawfile1,
          mass = mass_target,
          tol = tolerance_i,
          type = "xic"
        )

      data_w_ppm %>% filter(file %in% "2022-02-18_FDA_SpaceM/w2") %>%
        mutate(C20H12O5.H_xppm = FLU_signal[[1]]$intensities[1:10000],
               ppm_tolerance = tolerance_i / 2)
    }) %>% bind_rows()
  }, simplify = F) %>% bind_rows()

AMdata <-
data_w_ppm_ranges %>% group_by(ppm_tolerance, file) %>%
  summarise(zeros = mean(C20H12O5.H_xppm == 0))
```

```
## iterate over:
## datasets_names

cells <-
  pbsapply(data_w_ppm_ranges %>% group_split(ppm_tolerance, file), function(dataset_i) {
    ## in every iteration, different normalizations are assessed

    ## first, retreive prop and spec matrix for dataset_i

    prop_spec_matrix_OI <-
      prop_spec_matrix[[paste0(dataset_i$file[1], '/')]]


    ## now, the normalization parameters
    ## Rappez_et_al method; TIC-norm, zeros included

      AM_ion_intensity_OI <-
        dataset_i %>% mutate(C20H12O5.H = C20H12O5.H_xppm / TIC) %>%
        ## normalize C20H12O5.H  by TIC
        pull(C20H12O5.H)

      cell_ion_intensity <-
        cell_normalization_Rappez_et_al(
          overlap_proportion_matrix = prop_spec_matrix_OI$overlap_proportion_matrix,
          overlap_specificity_matrix = prop_spec_matrix_OI$overlap_specificity_matrix,
          AM_ion_intensity = AM_ion_intensity_OI,
          AM_proportion_treshold_global = 0.3,
          skip_AM_zeros = FALSE                ## keep zeros in AM vector
        )$cell_intensities

      data.frame(cell_id = rownames(cell_ion_intensity),
                 fraction_assigned = mean(!is.na(cell_ion_intensity)),
```

```
                 C20H12O5.H_calculated = cell_ion_intensity,
                 method = 'weighted average',
                 AM_normalization = 'TIC',
                 ppm_tolerance = dataset_i$ppm_tolerance[1],
                 AM_filter = '0.3',
                 AM_omit_zeros = "all values",
                 file = dataset_i$file[1])



  }, simplify = FALSE) %>% bind_rows()
```

```
cells <-
  cells %>% left_join(
    cells_SpaceM %>% select(file, cell_id, median_intensity.FITC, cell_area, eccentricity),
    by = c("file", "cell_id")
  )

cells <-
  cells %>% group_by(method,
                     ppm_tolerance,
                     AM_normalization,
                     AM_filter,
                     AM_omit_zeros,
                     file) %>%
  mutate(n = length(C20H12O5.H_calculated))
```

```
cell_summary <-
  cells %>% group_by(method, AM_normalization, AM_filter, AM_omit_zeros, ppm_tolerance, file) %>%
  filter(          ## filter out NAs for the correlations
    !is.na(median_intensity.FITC) &!is.na(C20H12O5.H_calculated)) %>%
  summarize(
    ppm_tolerance = ppm_tolerance[1],
    fraction_assigned = fraction_assigned[1],
    r = cor(
      x = median_intensity.FITC,
      y = C20H12O5.H_calculated,
      method = "spearman",
      use = "pairwise.complete.obs"
    ),
    rweight = weightedCorr(
      ## weighted median_intensity.FITC
      x = median_intensity.FITC,
      y = C20H12O5.H_calculated,
      method = "Spearman",
      weights = 1 - n_neighbours(x = median_intensity.FITC) /
        max(n_neighbours(median_intensity.FITC))
    ),
    n = n[1]
  )
```

```
Fig3F <-
cell_summary %>% group_by(ppm_tolerance) %>%
  summarise(rweight_mean = mean(rweight),
```

```
            rweight_sd = sd(rweight))  %>% {
  ggplot(data = .,
         aes(x = ppm_tolerance,
             y = rweight_mean, color = "A")) +
    #scale_y_continuous(limits = c(0, NA)) +
    #geom_boxplot(outlier.colour = NA, show.legend = F) +
    geom_point(size = 3,
               width = .01,
               show.legend = F) +
    geom_line(show.legend = F)+
    geom_errorbar(aes(ymin = rweight_mean- rweight_sd,
                      ymax = rweight_mean+ rweight_sd), color = "gray60")+
    scale_color_paletteer_d("ggthemes::calc") +
    labs(y = 'weighted  ',  x = "m/z bandwidth (ppm)") +
    theme_minimal() +
    geom_vline(xintercept = c(3,4), linetype = 2,color = c("red","gray60"))+
    theme(
      #axis.text.x = element_blank(),
      axis.title = element_text(size = 11),
      panel.grid = element_blank(),
      axis.ticks = element_line(),
      panel.background = element_rect(colour = "black", size = 1)
    )
            }
```

Fig3F

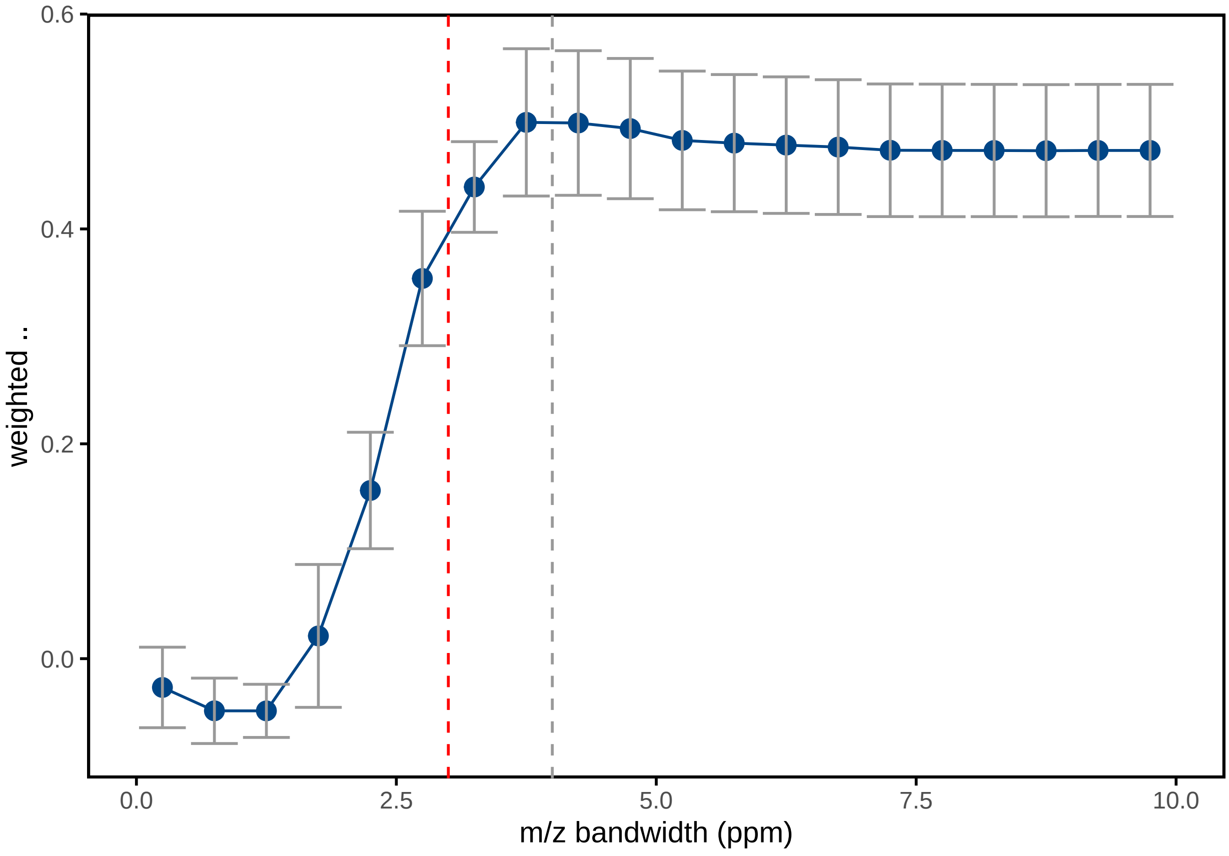

```
Fig3E <-
AMdata %>% group_by(ppm_tolerance) %>%
  summarise(zeros_mean = mean(zeros),
            zeros_sd = sd(zeros))  %>% {
              ggplot(data = .,
                     aes(x = ppm_tolerance,
                         y = zeros_mean, color = "A")) +
                #scale_y_continuous(limits = c(0, NA)) +
                #geom_boxplot(outlier.colour = NA, show.legend = F) +
                geom_point(size = 3,
                           width = .01,
                           show.legend = F) +
                geom_line(show.legend = F)+
                geom_errorbar(aes(ymin = zeros_mean- zeros_sd,
                                  ymax = zeros_mean+ zeros_sd), color = "gray60")+
                scale_color_paletteer_d("ggthemes::calc") +
                labs(y = expression(fraction~zeros~"in"~"[C"[20]*"H"[12]*"O"[5]*"-H]"^"-"),
                                    x = "m/z bandwidth (ppm)") +
                theme_minimal() +
                geom_vline(xintercept = c(3,4), linetype = 2,color = c("red","gray60"))+
                theme(
                  #axis.text.x = element_blank(),
                  axis.title = element_text(size = 11),
                  panel.grid = element_blank(),
                  axis.ticks = element_line(),
                  panel.background = element_rect(colour = "black", size = 1)
                )
            }

Fig3E
```

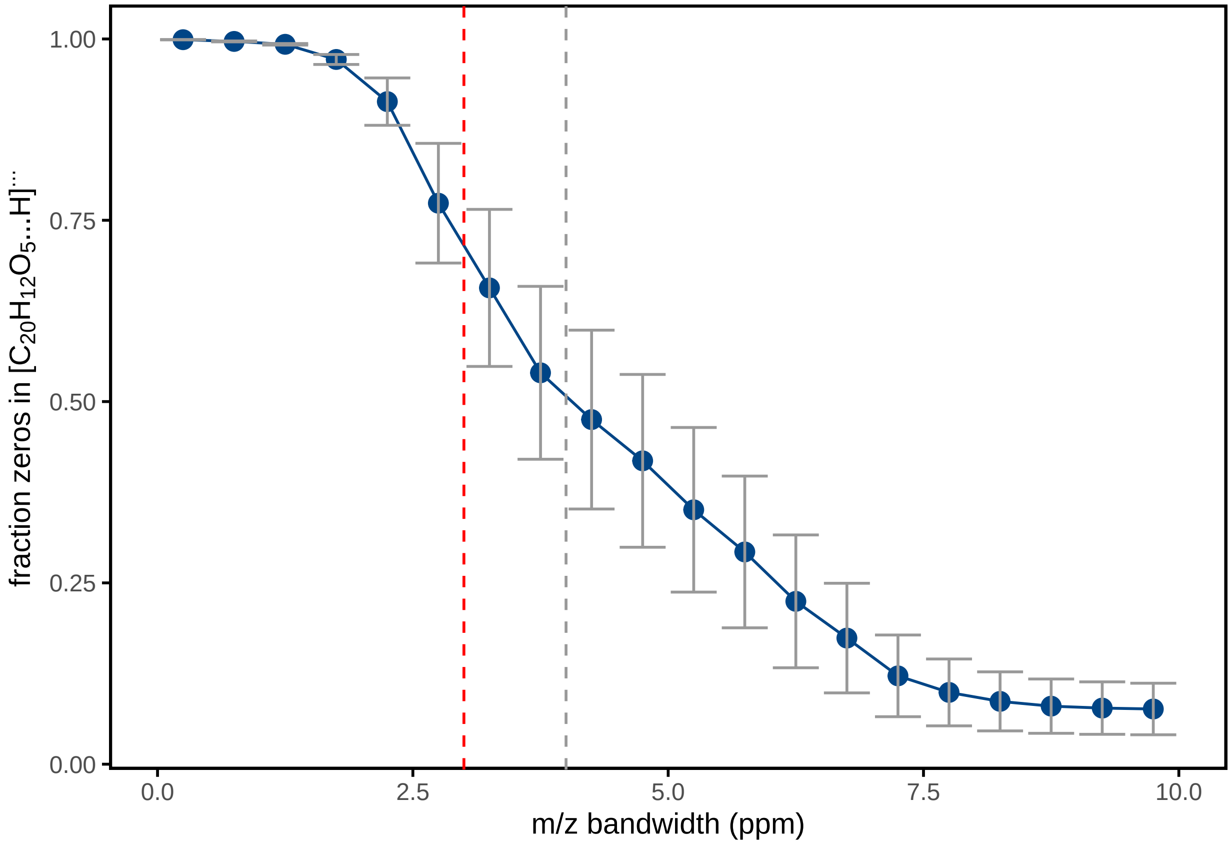


```
Fig3 <-
  cowplot::plot_grid(
    ncol = 1,
    cowplot::plot_grid(
      cowplot::plot_grid(
        Fig3A,
        Fig3B,
        align = "v",
        axis = "lr",
        ncol = 1,
        rel_heights = c(2, 1),
        #scale = .95,
        labels = c("", "B")
      ),
      Fig3C,
      Fig3D,
      nrow = 1,
      scale = .90,
      labels = c("A", "C", "D"),
      align = "h",
      axis = "tb",
      rel_widths = c(2, 1, 1.7)
    ),
    cowplot::plot_grid(Fig3E, Fig3F, scale = .9, labels = c("E", "F"))
  )+theme(panel.background = element_rect(color = NA, fill = "white"))

Fig3
```

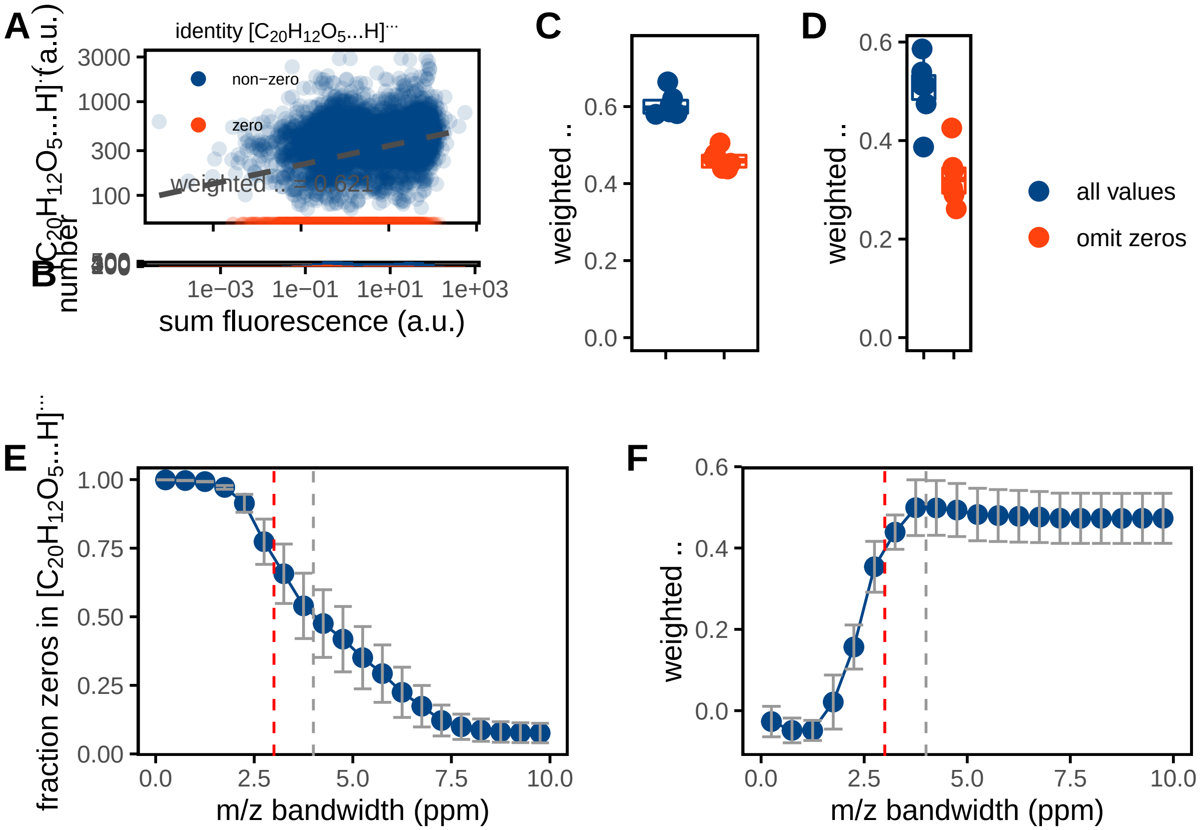


Figure 4

```
datasets_names <- data_w_ppm %>% group_by(file) %>% summarize(file = file[1]) %>% pull(file)

input_parameters <- data.frame(expand_grid(AM_proportion_treshold_global = seq(0.05,1,.1),
                       AM_specificity_treshold_cell = seq(0.05,1,.1)))

cells <-
  pbsapply(datasets_names, function(dataset_i) {
    ## in every iteration, different normalizations are assessed

    ## first, retreive prop and spec matrix for dataset_i
    prop_spec_matrix_OI <-
      prop_spec_matrix[[paste0(dataset_i, '/')]]


    mapply(AM_proportion_treshold_global_i = input_parameters$AM_proportion_treshold_global,
           AM_specificity_treshold_cell = input_parameters$AM_specificity_treshold_cell,
           function(AM_proportion_treshold_global_i,AM_specificity_treshold_cell){

             AM_ion_intensity_OI <-
               data_w_ppm %>% filter(file ==  dataset_i) %>%          ## filter dataset of interest
               mutate(C20H12O5.H = C20H12O5.H_4ppm / TIC) %>%         ## don't normalize C20H12O5.H
               pull(C20H12O5.H)

             cell_ion_intensity <-
               cell_normalization_Rappez_et_al(
                 overlap_proportion_matrix = prop_spec_matrix_OI$overlap_proportion_matrix,
```

```
                 overlap_specificity_matrix = prop_spec_matrix_OI$overlap_specificity_matrix,
                 AM_ion_intensity = AM_ion_intensity_OI,
                 AM_proportion_treshold_global = AM_proportion_treshold_global_i,
                 AM_specificity_treshold_cell = AM_specificity_treshold_cell,
                 skip_AM_zeros = FALSE                ## don't omit zeros in AM vector
               )$cell_intensities

             data.frame(cell_id = rownames(cell_ion_intensity),
                        C20H12O5.H_calculated = cell_ion_intensity,
                        method = 'weighted average',
                        AM_proportion_treshold_global = AM_proportion_treshold_global_i,
                        AM_specificity_treshold_cell = AM_specificity_treshold_cell,
                        file = dataset_i)

    }, SIMPLIFY = F) %>% bind_rows()

  }, simplify = FALSE) %>% bind_rows()



cells <-
  cells %>% left_join(
    cells_SpaceM %>% select(file, cell_id, median_intensity.FITC, cell_area, eccentricity),
    by = c("file", "cell_id")
  )

cells <-
  cells %>% group_by(method,
                     AM_specificity_treshold_cell,
                     AM_proportion_treshold_global,
                     file) %>%
  mutate(n = length(C20H12O5.H_calculated))
```

```
cell_summary <-
  cells %>%
  group_by(method, AM_specificity_treshold_cell, AM_proportion_treshold_global,file) %>%
  filter(          ## filter out NAs for the correlations
    !is.na(median_intensity.FITC) &!is.na(C20H12O5.H_calculated)) %>%
  summarize(
    r = cor(
      x = median_intensity.FITC,
      y = C20H12O5.H_calculated,
      method = "spearman",
      use = "pairwise.complete.obs"
    ),
    rweight = weightedCorr(
      ## weighted median_intensity.FITC
      x = median_intensity.FITC,
      y = C20H12O5.H_calculated,
      method = "Spearman",
      weights = 1 - n_neighbours(x = median_intensity.FITC) /
        max(n_neighbours(median_intensity.FITC))
    ),
    fraction_cells = length(C20H12O5.H_calculated) / n[1]
```

```
  )
```

```
sec_axis_factor <- 1.3


Fig4B <-
  cell_summary %>%
  filter(between(x = AM_proportion_treshold_global, left = .3, right = .4)) %>%
  group_by(AM_specificity_treshold_cell) %>%
  summarise(
    rweight_mean = mean(rweight),
    rweight_sd = sd(rweight),
    fraction_cells_mean = mean(fraction_cells) / sec_axis_factor,
    fraction_cells_sd = sd(fraction_cells) / sec_axis_factor
  )  %>% {
    ggplot(data = .,
           aes(x = AM_specificity_treshold_cell,
               y = rweight_mean)) +
      scale_y_continuous(
        limits = c(0, NA),
        sec.axis = sec_axis(trans =  ~ . * sec_axis_factor, name =
                              "fraction of assigned cells")
      ) +
      #geom_boxplot(outlier.colour = NA, show.legend = F) +
      geom_point(
        size = 3,
        color = palettes_d$ggthemes$calc[1],
        show.legend = F
      ) +
      geom_line(show.legend = F, color = palettes_d$ggthemes$calc[1]) +
      geom_errorbar(
        aes(ymin = rweight_mean - rweight_sd,
            ymax = rweight_mean + rweight_sd),
        color = "gray60",
        width = .05
      ) +
      geom_point(aes(y = fraction_cells_mean),
                 color = palettes_d$ggthemes$calc[2],
                 size = 3) +
      geom_line(
        aes(y = fraction_cells_mean),
        color = palettes_d$ggthemes$calc[2],
        linetype = 2
      ) +
      geom_errorbar(
        aes(
          ymin = fraction_cells_mean - fraction_cells_sd,
          ymax = fraction_cells_mean + fraction_cells_sd
        ),
        color = "gray60",
        width = .05
      ) +
      scale_color_paletteer_d("ggthemes::calc") +
      labs(y = 'weighted  ',  x = "sampling specificity cut-off") +
```

```
    theme_minimal() +
    theme(
      axis.title = element_text(size = 11),
      axis.title.y.left = element_text(colour = palettes_d$ggthemes$calc[1]),
      axis.title.y.right = element_text(colour = palettes_d$ggthemes$calc[2]),
      axis.text.y.left = element_text(colour = palettes_d$ggthemes$calc[1]),
      axis.text.y.right = element_text(colour = palettes_d$ggthemes$calc[2]),
      panel.grid = element_blank(),
      axis.ticks = element_line(),
      panel.background = element_rect(colour = "black", size = 1)
    )
}

Fig4B
```

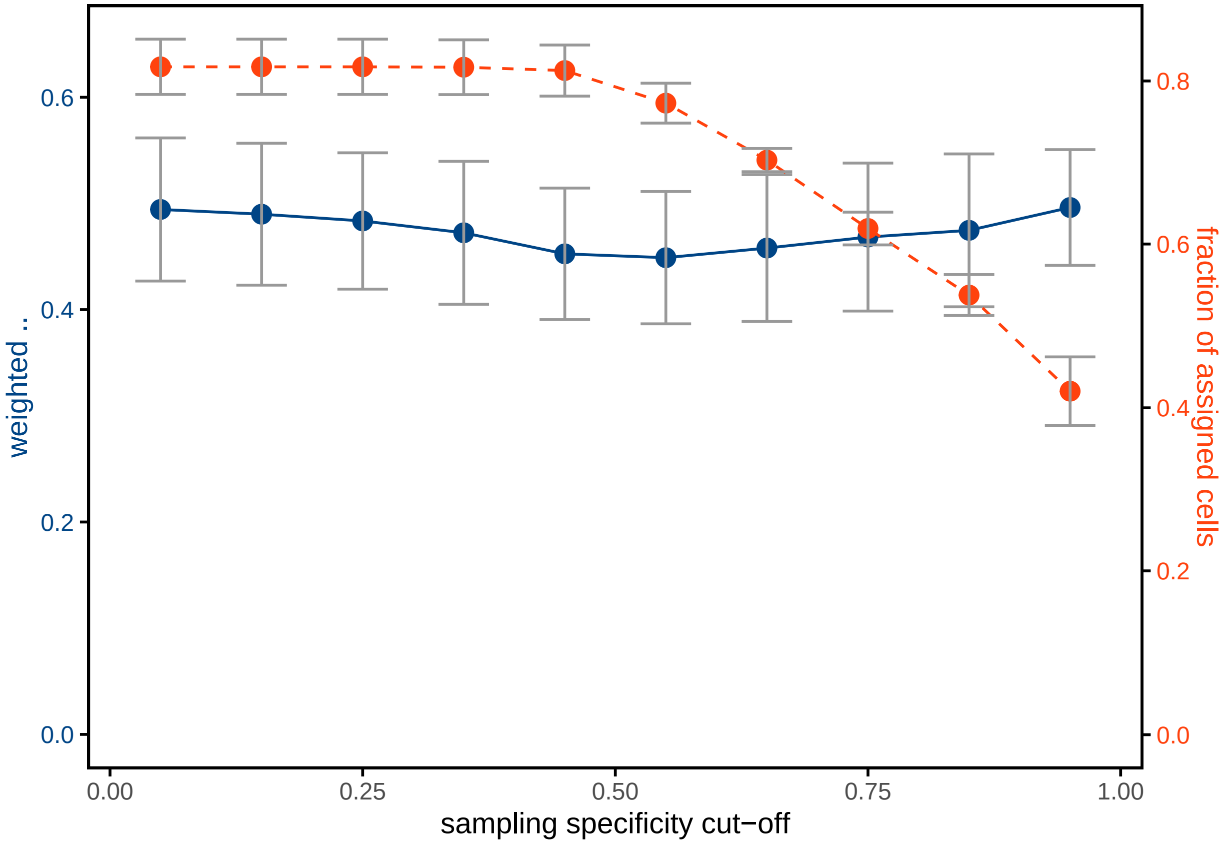


```
sec_axis_factor <- 1.3


Fig4A <-
  cell_summary %>%
  filter(between(x = AM_specificity_treshold_cell, left = 0, right = .1)) %>%
  group_by(AM_proportion_treshold_global) %>%
  summarise(
    rweight_mean = mean(rweight),
    rweight_sd = sd(rweight),
    fraction_cells_mean = mean(fraction_cells) / sec_axis_factor,
    fraction_cells_sd = sd(fraction_cells) / sec_axis_factor
```

```
)  %>% {
  ggplot(data = .,
         aes(x = AM_proportion_treshold_global,
             y = rweight_mean)) +
    scale_y_continuous(
      limits = c(0, NA),
      sec.axis = sec_axis(trans =  ~ . * sec_axis_factor, name =
                            "fraction of assigned cells")
    ) +
    #geom_boxplot(outlier.colour = NA, show.legend = F) +
    geom_point(
      size = 3,
      color = palettes_d$ggthemes$calc[1],
      show.legend = F
    ) +
    geom_line(show.legend = F, color = palettes_d$ggthemes$calc[1]) +
    geom_errorbar(
      aes(ymin = rweight_mean - rweight_sd,
          ymax = rweight_mean + rweight_sd),
      color = "gray60",
      width = .05
    ) +
    geom_point(aes(y = fraction_cells_mean),
               color = palettes_d$ggthemes$calc[2],
               size = 3) +
    geom_line(
      aes(y = fraction_cells_mean),
      color = palettes_d$ggthemes$calc[2],
      linetype = 2
    ) +
    geom_errorbar(
      aes(
        ymin = fraction_cells_mean - fraction_cells_sd,
        ymax = fraction_cells_mean + fraction_cells_sd
      ),
      color = "gray60",
      width = .05
    ) +
    scale_color_paletteer_d("ggthemes::calc") +
    labs(y = 'weighted  ',  x = "sampling proportion cut-off") +
    theme_minimal() +
    theme(
      axis.title = element_text(size = 11),
      axis.title.y.left = element_text(colour = palettes_d$ggthemes$calc[1]),
      axis.title.y.right = element_text(colour = palettes_d$ggthemes$calc[2]),
      axis.text.y.left = element_text(colour = palettes_d$ggthemes$calc[1]),
      axis.text.y.right = element_text(colour = palettes_d$ggthemes$calc[2]),
      panel.grid = element_blank(),
      axis.ticks = element_line(),
      panel.background = element_rect(colour = "black", size = 1)
    )
}
```

```
Fig4A
```

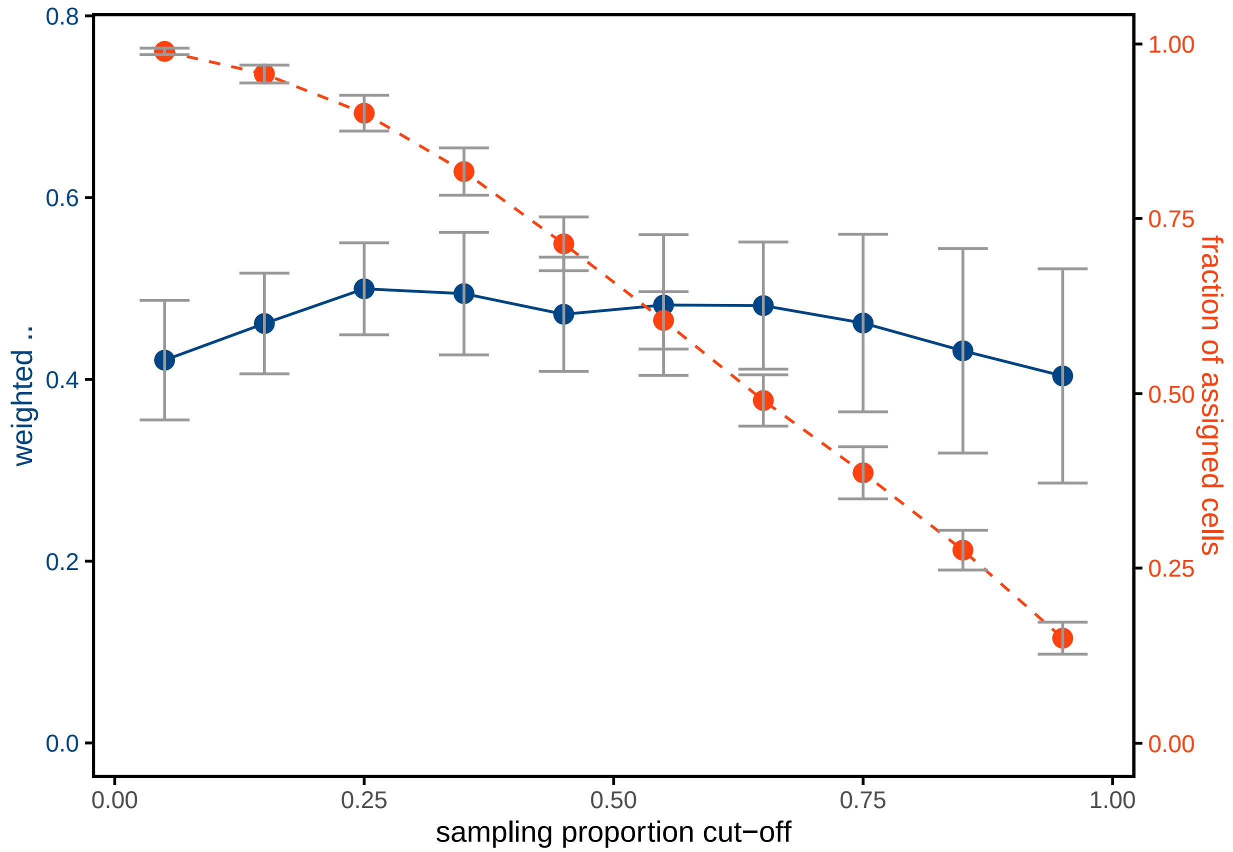


```
Fig4C <-
cell_summary %>%
  group_by(AM_proportion_treshold_global, AM_specificity_treshold_cell) %>%
  summarise(rweight_mean = mean(rweight),
            rweight_sd = sd(rweight),
            fraction_cells_mean = mean(fraction_cells),
            fraction_cells_sd = sd(fraction_cells))  %>% {
              ggplot(data = .,
                     aes(x = AM_proportion_treshold_global,
                         y = AM_specificity_treshold_cell,
                         fill = rweight_mean)) +
                coord_fixed(expand = FALSE)+
                geom_tile(show.legend = T) +
                scale_x_continuous(limits = c(-0.01,1.01))+
                scale_y_continuous(limits = c(-0.01,1.01))+
                scale_fill_gradient(low = "gray30",high = palettes_d$ggthemes$calc[2])+
                labs(fill = 'mean weighted  ',
                     x = "sampling proportion cut-off",
                     y = "sampling specificity cut-off") +
                theme_minimal() +
                theme(
                  axis.title = element_text(size = 11), legend.position = "top",
                  panel.grid = element_blank(),
                  axis.ticks = element_line(),
                  panel.background = element_rect(colour = "black", size = 1)
```

```
            )
        }
```

```
Fig4C
```

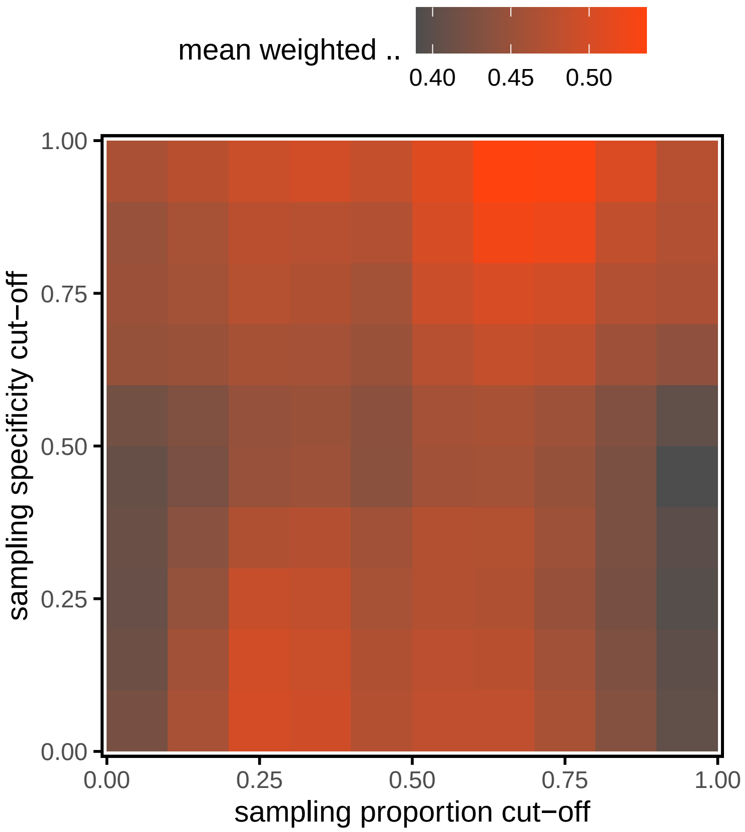


```
Fig4D <-
cell_summary %>%
  group_by(AM_proportion_treshold_global, AM_specificity_treshold_cell) %>%
  summarise(rweight_mean = mean(rweight),
            rweight_sd = sd(rweight),
            fraction_cells_mean = mean(fraction_cells),
            fraction_cells_sd = sd(fraction_cells) )  %>% {
              ggplot(data = .,
                     aes(x = AM_proportion_treshold_global,
                         y = AM_specificity_treshold_cell,
                         fill = fraction_cells_mean)) +
                coord_fixed(expand = FALSE)+
                geom_tile(show.legend = T) +
                scale_x_continuous(limits = c(-0.01,1.01))+
                scale_y_continuous(limits = c(-0.01,1.01))+
                scale_fill_gradient(low = "gray30",high = palettes_d$ggthemes$calc[2])+
                labs(fill = 'fraction of assigned cells',
                     x = "sampling proportion cut-off",
                     y = "sampling specificity cut-off") +
                theme_minimal() +
                theme(
                  axis.title = element_text(size = 11), legend.position = "top",
```

```
                  panel.grid = element_blank(),
                  axis.ticks = element_line(),
                  panel.background = element_rect(colour = "black", size = 1)
                )
            }

Fig4D
```

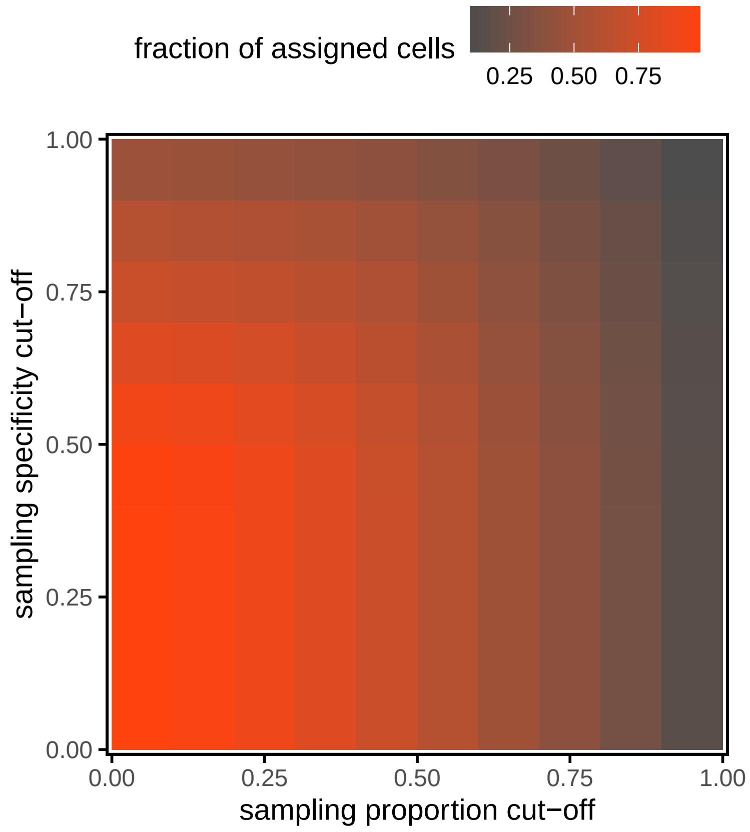


```
Fig4 <-
  plot_grid(
    Fig4A,
    Fig4B,
    Fig4C,
    Fig4D,
    ncol = 2,
    rel_heights = c(.6, 1),
    align = "hv",
    scale = .95,
    labels = c("A", "B", "C", "D")
  ) + theme(panel.background = element_rect(color = NA, fill = "white"))

Fig4
```

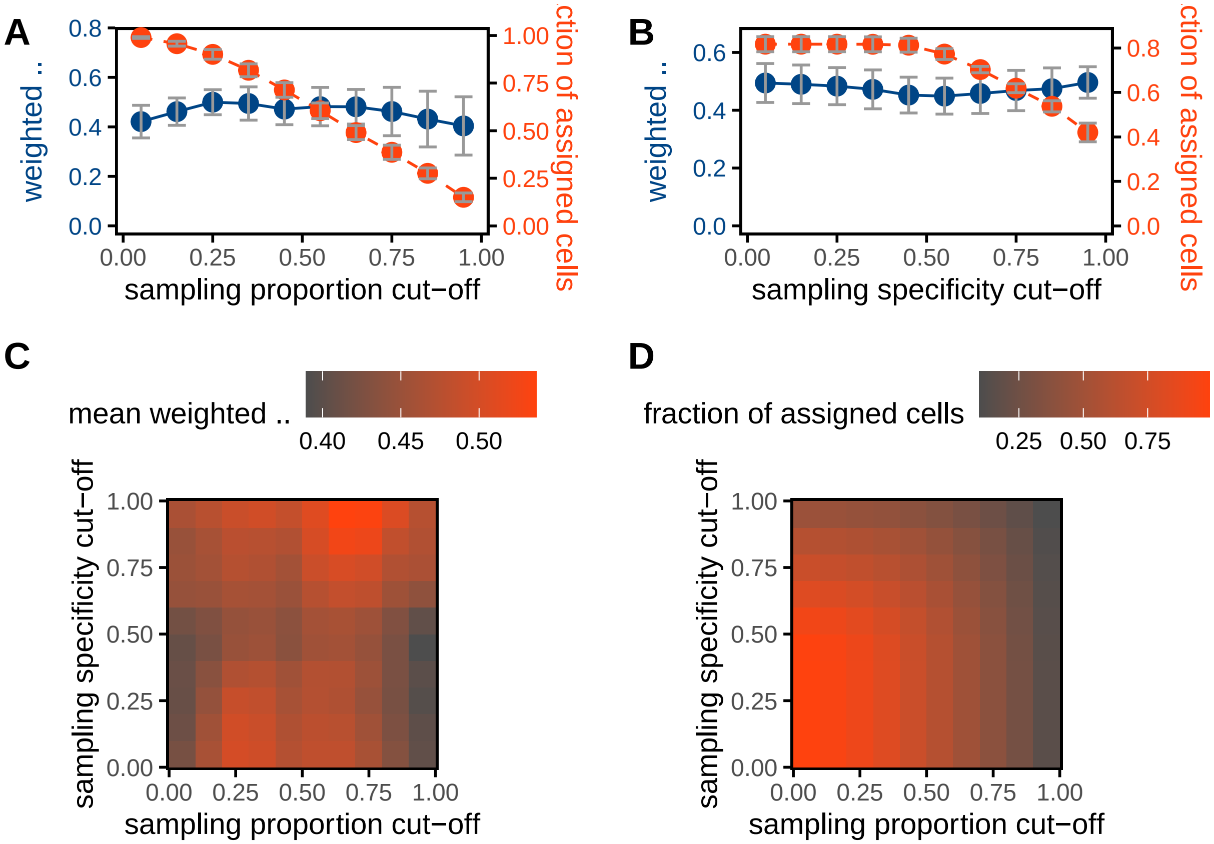


#### Extra figures

```
datasets_names <- data_w_ppm %>% group_by(file) %>% summarize(file = file[1]) %>% pull(file)


cells <-
  sapply(datasets_names, function(dataset_i) {
    ## in every iteration, different normalizations are assessed

    ## first, retreive prop and spec matrix for dataset_i
    prop_spec_matrix_OI <-
      prop_spec_matrix[[paste0(dataset_i, '/')]]


    ## now, the normalization parameters
    list({
      ## Rappez_et_al method; no TIC-norm, zeros included

      AM_ion_intensity_OI <-
        data_w_ppm %>% filter(file ==  dataset_i) %>%       ## filter dataset of interest
        mutate(C20H12O5.H = C20H12O5.H_4ppm) %>%         ## don't normalize C20H12O5.H
        pull(C20H12O5.H)

      cell_ion_intensity <-
        cell_normalization_Rappez_et_al(
          overlap_proportion_matrix = prop_spec_matrix_OI$overlap_proportion_matrix,
```

```
        overlap_specificity_matrix = prop_spec_matrix_OI$overlap_specificity_matrix,
        AM_ion_intensity = AM_ion_intensity_OI,
        AM_proportion_treshold_global = 0.3,
        skip_AM_zeros = FALSE                           ## don't omit zeros in AM vector
      )$cell_intensities

    data.frame(cell_id = rownames(cell_ion_intensity),
               C20H12O5.H_calculated = cell_ion_intensity,
               method = 'WA',
               AM_normalization = 'none',
               AM_filter = '0.3',
               AM_omit_zeros = "keep zeros",
               file = dataset_i)
  },
  {
    ## Rappez_et_al method; suppr model norm, zeros included

    AM_ion_intensity_OI <-
      data_w_ppm %>% filter(file ==  dataset_i) %>%           ## filter dataset of interest
      mutate(C20H12O5.H = C20H12O5.H_4ppm / ion_suppr_model) %>%
      ## don't normalize C20H12O5.H
      pull(C20H12O5.H)

    cell_ion_intensity <-
      cell_normalization_Rappez_et_al(
        overlap_proportion_matrix = prop_spec_matrix_OI$overlap_proportion_matrix,
        overlap_specificity_matrix = prop_spec_matrix_OI$overlap_specificity_matrix,
        AM_ion_intensity = AM_ion_intensity_OI,
        AM_proportion_treshold_global = 0.3,
        skip_AM_zeros = FALSE             ## don't omit zeros in AM vector
      )$cell_intensities

    data.frame(cell_id = rownames(cell_ion_intensity),
               C20H12O5.H_calculated = cell_ion_intensity,
               method = 'WA',
               AM_normalization = 'ion suppression model',
               AM_filter = '0.3',
               AM_omit_zeros = "keep zeros",
               file = dataset_i)
  },{
    ## LIM;  zeros included

    AM_ion_intensity_OI <-
      data_w_ppm %>% filter(file ==  dataset_i) %>%         ## filter dataset of interest
      mutate(C20H12O5.H = C20H12O5.H_4ppm ) %>%           ## don't normalize C20H12O5.H
      pull(C20H12O5.H)

    cell_ion_intensity <-
      cell_normalization_NNLS(
        overlap_proportion_matrix = prop_spec_matrix_OI$overlap_proportion_matrix,
        AM_ion_intensity = AM_ion_intensity_OI,
        AM_proportion_treshold_per_cell = 0.3,
        skip_AM_zeros = FALSE                       ## keep zeros in AM vector
```

```
        )$cell_intensities

      data.frame(cell_id = names(cell_ion_intensity),
                 C20H12O5.H_calculated = cell_ion_intensity,
                 method = 'LIM',
                 AM_normalization = 'none',
                 AM_filter = '0.3',
                 AM_omit_zeros = "keep zeros",
                 file = dataset_i)



    },{
      ## LIM; suppr model norm, zeros included

      AM_ion_intensity_OI <-
        data_w_ppm %>% filter(file ==  dataset_i) %>%            ## filter dataset of interest
        mutate(C20H12O5.H = C20H12O5.H_4ppm / ion_suppr_model) %>%
        ## don't normalize C20H12O5.H
        pull(C20H12O5.H)

      cell_ion_intensity <-
        cell_normalization_NNLS(
          overlap_proportion_matrix = prop_spec_matrix_OI$overlap_proportion_matrix,
          AM_ion_intensity = AM_ion_intensity_OI,
          AM_proportion_treshold_per_cell = 0.3,
          skip_AM_zeros = FALSE                ## keep zeros in AM vector
        )$cell_intensities

      data.frame(cell_id = names(cell_ion_intensity),
                 C20H12O5.H_calculated = cell_ion_intensity,
                 method = 'LIM',
                 AM_normalization = 'ion suppression model',
                 AM_filter = '0.3',
                 AM_omit_zeros = "keep zeros",
                 file = dataset_i)



    })  %>% bind_rows()

  }, simplify = FALSE) %>% bind_rows()
```

```
cells <-
  cells %>% left_join(
    cells_SpaceM %>% select(file, cell_id, median_intensity.FITC, cell_area, eccentricity),
    by = c("file", "cell_id")
  )

cells <-  cells %>% group_by(method, AM_normalization,AM_filter,AM_omit_zeros,file) %>%
  mutate(n = length(C20H12O5.H_calculated))
```

```
cell_summary <-
  cells %>%
```

```
  group_by(method, AM_normalization, AM_filter, AM_omit_zeros, file) %>%
  filter(          ## filter out NAs for the correlations
    !is.na(median_intensity.FITC) &!is.na(C20H12O5.H_calculated)) %>%
  summarize(
    r = cor(
      x = median_intensity.FITC,
      y = C20H12O5.H_calculated,
      method = "spearman",
      use = "pairwise.complete.obs"
    ),
    rweight = weightedCorr(
      ## weighted median_intensity.FITC
      x = median_intensity.FITC,
      y = C20H12O5.H_calculated,
      method = "Spearman",
      weights = 1 - n_neighbours(x = median_intensity.FITC) /
        max(n_neighbours(median_intensity.FITC))
    ),
    n = n[1]
  )


cell_summary <-
  cell_summary %>% mutate(AM_normalization =
                            factor(AM_normalization, levels = c("none", 'ion suppression model')))
```

```
FigS1B <-
cell_summary %>%  {
  ggplot(data = .,
         aes(x = AM_normalization,
             y = rweight, color = AM_normalization))+
    #geom_vline(xintercept = 3, linetype = 3, color = "gray30")+
    facet_wrap(~method)+
    scale_y_continuous(limits = c(0,NA))+
    geom_boxplot(outlier.colour = NA, show.legend = F)+
    geom_jitter(size = 3, width = .08, show.legend = T)+
    scale_color_paletteer_d("ggthemes::calc")+
    #geom_line(aes(group = file, color = file), linetype = 2, alpha = .7)+
    labs(y = 'weighted  ',  x = "", color = "normalisation")+
    theme_minimal()+
    theme(axis.text.x = element_blank(),
          axis.title = element_text(size = 11), panel.grid = element_blank(),
          axis.ticks = element_line(),
          panel.background = element_rect(colour = "black", size = 1))
}

FigS1B
```

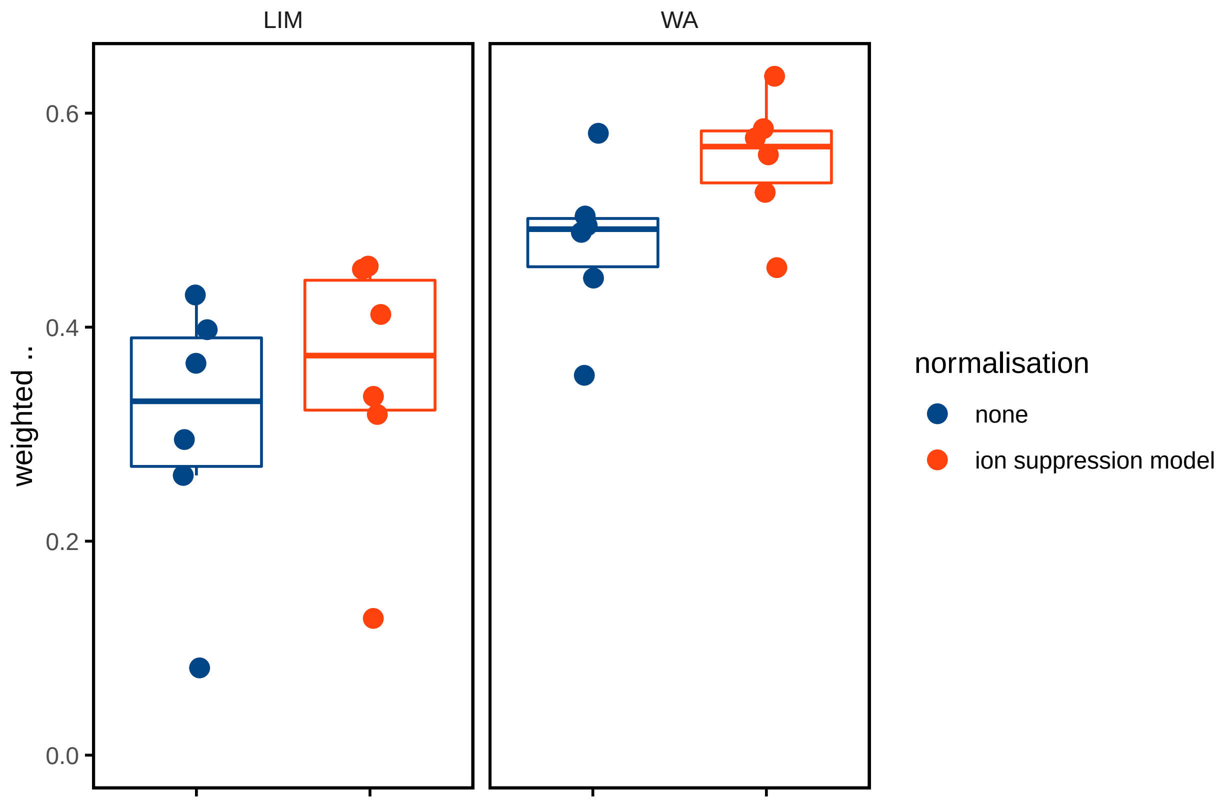


```
sampling_proportions <- unlist(sapply(prop_spec_matrix, function(i){
  f <- unlist(i$overlap_proportion_matrix)
  f[f!=0]
}))

specificity_proportions <- unlist(sapply(prop_spec_matrix, function(i){
  f <- unlist(i$overlap_specificity_matrix)
  f[f!=0]
}))


mean(
sampling_proportions < 1
)
```

```
## [1] 0.977851
```

```
p <- hist(sampling_proportions, breaks = 30)
```

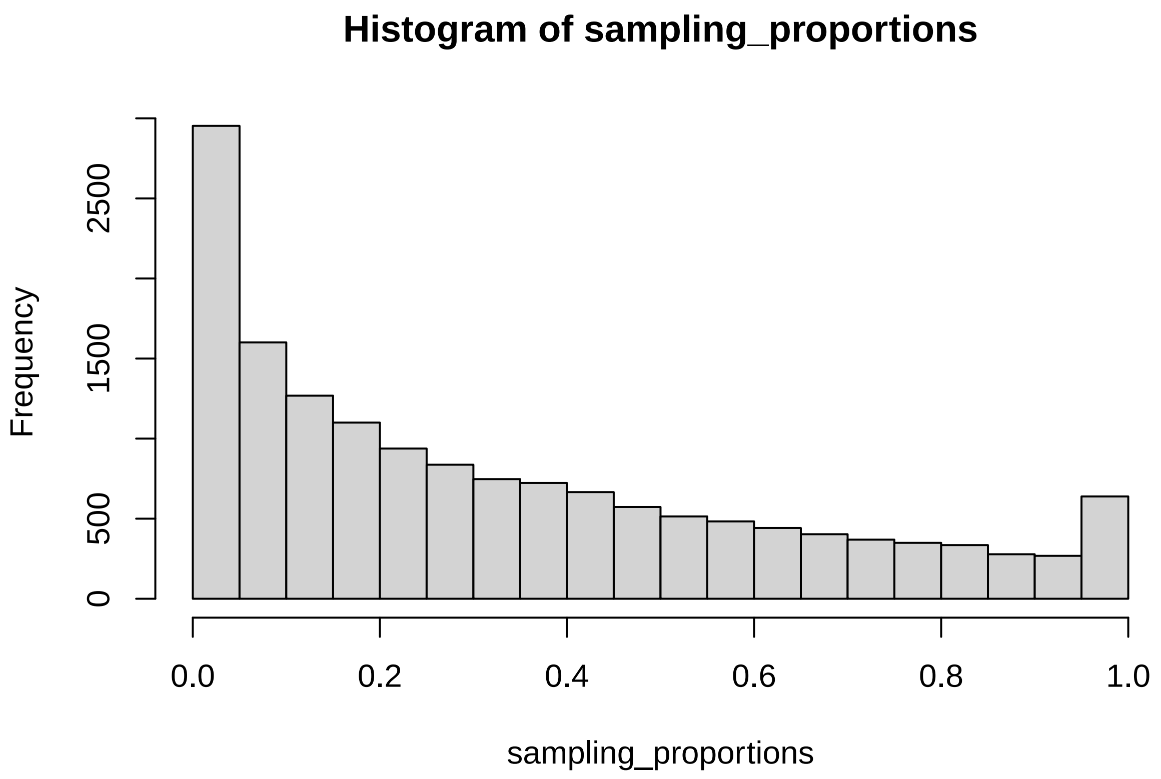


```
q <- hist(specificity_proportions, breaks = 30)
```

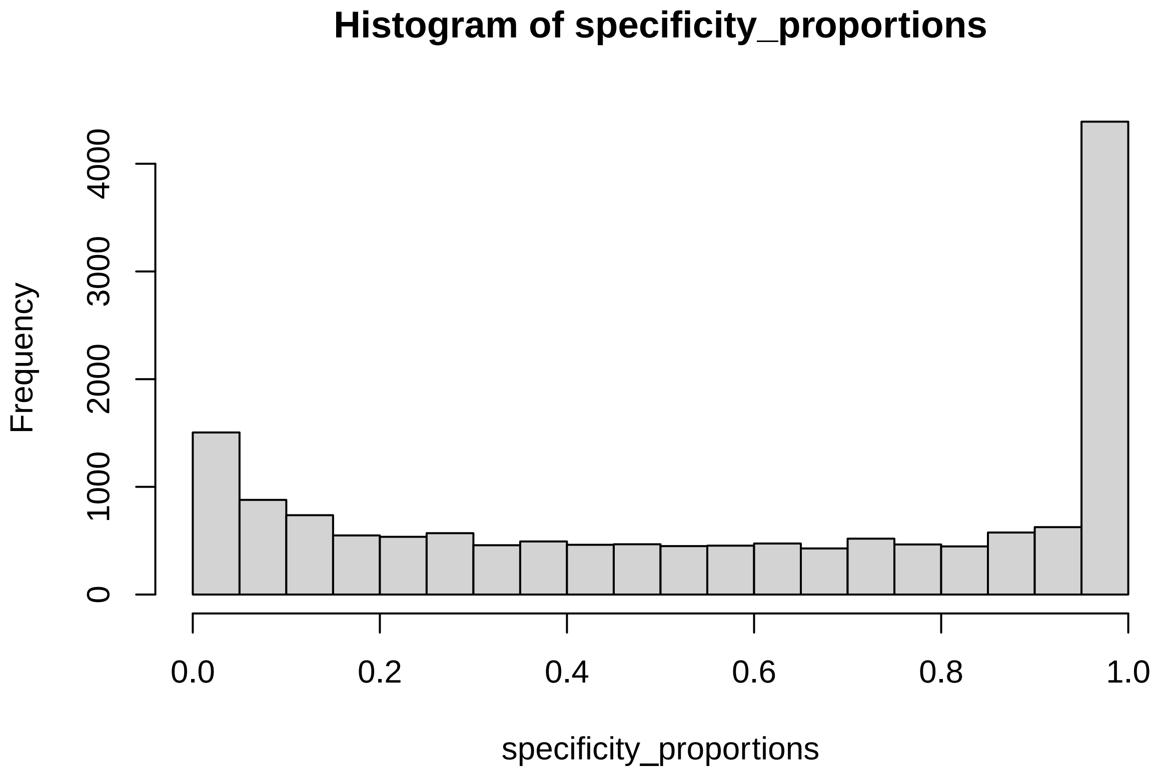


```
df <-
rbind(data.frame(x = p$breaks[-1],
           y = cumsum(p$counts / sum(p$counts)),
```

```
          set = "sampling proportions"),
     data.frame(x = q$breaks[-1],
                y = cumsum(q$counts / sum(q$counts)),
                set = "sampling specificities"))
```

```
FigS1A <-
  ggplot(data = df,
         aes(x = x, y = y, color = set)) +
  geom_point() +
  geom_line() +
  scale_color_paletteer_d("ggthemes::calc") +
  #geom_line(aes(group = file, color = file), linetype = 2, alpha = .7)+
  labs(y = 'cumulative fraction',  x = "sampling specificity or proportion", color = "") +
  theme_minimal() +
  theme(
    legend.position = c(0.05, 0.9),
    legend.justification = 0,
    legend.background = element_rect(color = NA, fill = NA),
    axis.title = element_text(size = 11),
    panel.grid = element_blank(),
    axis.ticks = element_line(),
    panel.background = element_rect(colour = "black", size = 1)
  )

FigS1A
```

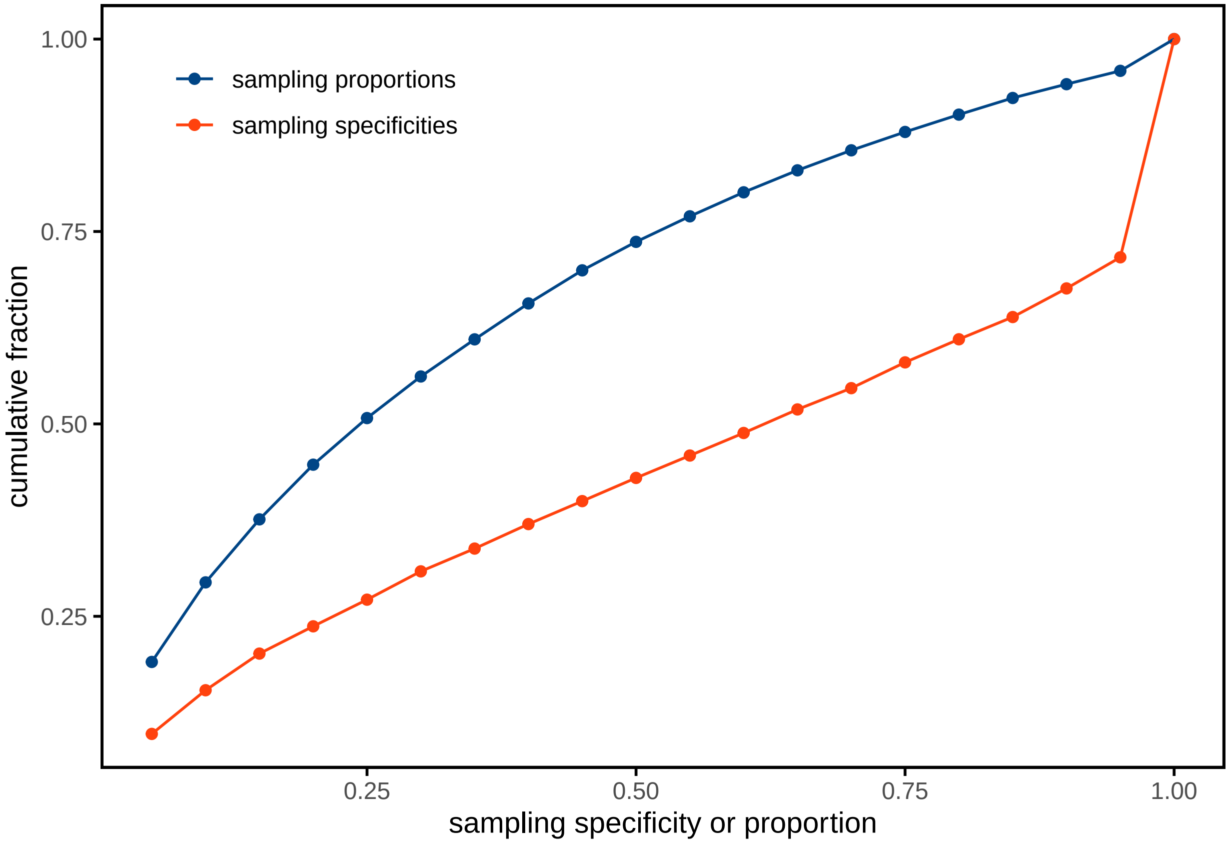

### Supplemental Information 1, R-code used for analysis, part II

2022-07-25

```
library(reticulate)
library(dplyr)
library(ggplot2)
library(paletteer)
library(quantreg)
library(tidyverse)
library(ggdendro)
library(dendextend)
library(Seurat)
```

##### Figure 2E and 2F

The co-culture dataset is available in the MetaboLights repository under the accession number MTBLS78 (https://www.ebi.ac.uk/metabolights/MTBLS78).

```
## load conda env to load Python objects
use_condaenv(condaenv = 'notebook', required = TRUE)
np <- import(module = "numpy", as = 'np')


## load sc dataset
dataset_co_cult <- np$load('marks_flitered_fluo.npy', allow_pickle = TRUE)

names(dataset_co_cult) <-
  c(
    "norm_MM",
    "cell_marks",
    "nucl_fluo",
    "cell_fluo",
    "marks_fluo",
    "marks_cell_overlap",
    "mark_area",
    "overlap_indices",
    "marks_fluo_overlap",
    "cell_area",
    "marks_cell_overlap_indexes",
    "marks_cellLabels",
    "marks_samplingArea",
    "pmi",
    "overLaps")


overlap_matrix <- matrix(data = 0, nrow = length(dataset_co_cult$cell_area),
                         ncol = length(dataset_co_cult$mark_area))
rownames(overlap_matrix) <- names(dataset_co_cult$cell_area)
```

```r
colnames(overlap_matrix) <- names(dataset_co_cult$mark_area)


for(cell_i in seq_along(dataset_co_cult$cell_marks)){

  for(AM_n in seq_along(dataset_co_cult$marks_cell_overlap[[cell_i]])){

    AM_i <- dataset_co_cult$cell_marks[[cell_i]][AM_n]
    overlap_matrix[cell_i, AM_i] <- dataset_co_cult$marks_cell_overlap[[cell_i]][AM_n]
  }
}

overlap_matrix[] <-
  sapply(seq(dim(overlap_matrix)[2]), function(AM_i){
    overlap_matrix[,AM_i] / dataset_co_cult$mark_area[[AM_i]]
  })
```

```r
## load metadata
sc <- read.csv(file = 'MORPHnMOL.csv')

sc_fluorescence <-
  data.frame(GFP = sc$Intensity_MeanIntensity_GFP_quantif,
             mCherry = sc$Intensity_MeanIntensity_mCherry_quantif)


sc_fluorescence <- sc_fluorescence %>% mutate(ratio = log(GFP /  mCherry))



ggplot(data = sc_fluorescence,
       aes(x = GFP, y = mCherry, color = ifelse(ratio< .8,"HeLa","NIH3T3")))+
  geom_point()+
  scale_y_log10()+
  scale_x_log10()
```

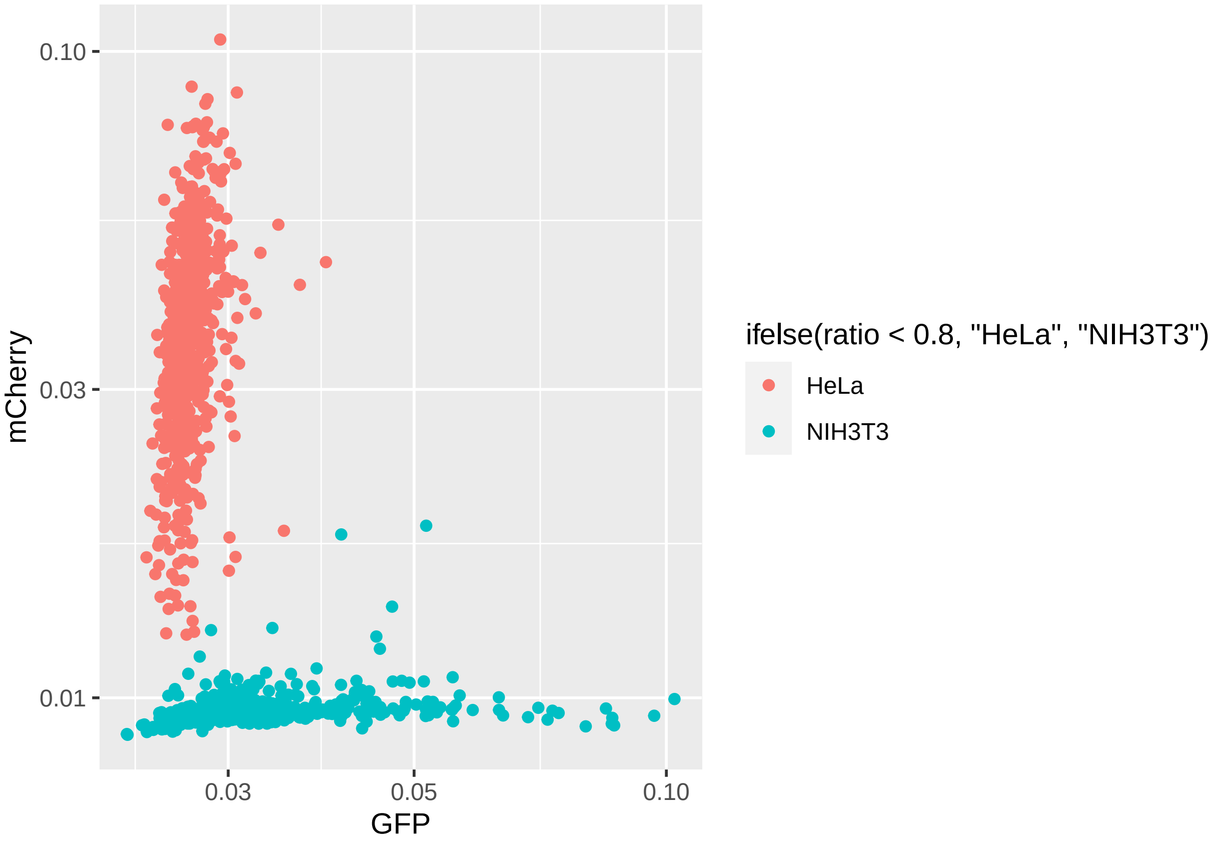


```
AMdata_sc <- read.csv(file = 'sm_annotation_detections.csv')

## loading SpaceM normalization functions
source('normalization_functions.r')

molecules <- intersect(colnames(AMdata_sc),colnames(sc))

AMdata_sc_plain <- AMdata_sc[,molecules]

overlap_spec_matrix <- overlap_matrix
overlap_spec_matrix[] <-
  t(apply(overlap_matrix, 1, function(i){
    i / sum(i)
  }))

norm <-
  cell_ion_intensity <-
  cell_normalization_Rappez_et_al(
    overlap_proportion_matrix = overlap_matrix,
    overlap_specificity_matrix = overlap_spec_matrix,
    AM_ion_intensity = AMdata_sc_plain,
    AM_proportion_treshold_global = 0.3,
    skip_AM_zeros = FALSE                                ## keep zeros in AM vector
  )
```

```
cell_intensities <- norm$cell_intensities[
  match(sc$ObjectNumber,rownames(norm$cell_intensities)),]

AMdata_sc$am_ratio <- apply(overlap_matrix,2,function(i){
  sum(i)
})


AMdata_sc <- AMdata_sc %>%
  mutate(ratio = AMdata_sc$C45H82NO8P / am_ratio,
         ratio = ratio / median(ratio[am_ratio == 1], na.rm = T))
```

```
### performing the regressions:

plots <-
  sapply(seq_along(AMdata_sc_plain), function(molecule_n){

    #molecule_i <- AMdata_sc_corrected[[molecule_n]]
    molecule_i <- AMdata_sc_plain[[molecule_n]]


    input_df <- data.frame(signal = molecule_i,
                           am_ratio = AMdata_sc$am_ratio) %>%
      mutate(ratio = signal / am_ratio,
             ratio = ratio / median(ratio[am_ratio == 1], na.rm = T),
             log_ratio = log(ratio, base = 10)) %>%
      filter(is.finite(log_ratio))

    if(dim(input_df%>%
          filter(is.finite(log_ratio) & am_ratio > 0.1 ))[1] < 10){
      return(NA)
    }

    model <- quantreg::rq(
      formula = log(ratio, base = 10) ~ log(am_ratio, base = 10),
      tau = .5,
      data =   input_df %>%
        filter(is.finite(log_ratio) & am_ratio > 0.1 )
    )

    input_df$pred_ratio <- 10 ^ predict(model,
                                        newdata = input_df %>%
                                          mutate(am_sampling_ratio = log(am_ratio, base = 10)))


    input_df %>%
      ggplot(data = .,
             aes(x = am_ratio,
                 y = ratio))+
      scale_y_log10(breaks = c(.25,5,.75,1,2.5,5,7.5,10,25,50,75,100))+
      #scale_y_continuous(limits = c(0,50))+
      scale_x_log10(breaks = seq(0,1,.2), limits = c(.1,1))+
      #facet_wrap(analyte_nice~., nrow = 4)+
      geom_point(alpha = .35, size = .7, color = palettes_d$ggthemes$calc[4])+
```

```
      geom_line(aes(y = pred_ratio), color = "red")+
      scale_color_paletteer_d("ggthemes::calc") +
      #geom_smooth(method = "lm",linetype = 2, se = F, color = "gray30")+
      geom_hline(yintercept = 1, color = "gray30",
                 linetype = 2)+
      labs(x = "sampling proportion",
           y = expression(italic( )~'='~'MS-signal '/' sampling proportion (a.u.)'))+
      #paste0('  = MS-signal / sampling proportion (a.u.)'))+
      theme_minimal()+
      theme(axis.text.x = element_text(size = 9),
            axis.title = element_text(size = 11), panel.grid = element_blank(),
            axis.ticks = element_line(),
            panel.background = element_rect(colour = "black", size = 1), strip.text =
              element_text(size = 9, face = "bold"))


  }, simplify = F)

### one of the molecules:
molecule_n <- 25
plots[[molecule_n]]
```

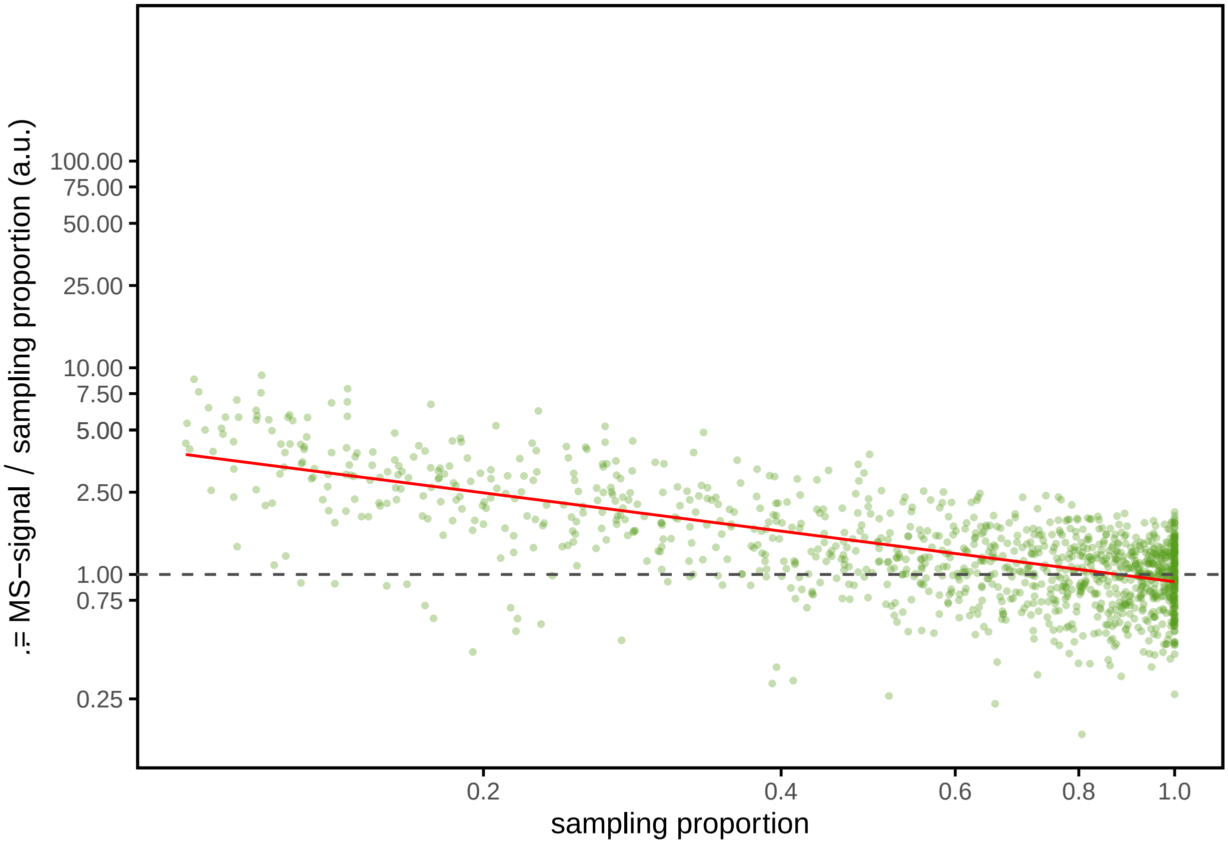


compensate for ion suppression using the upper regressions and perform normalizations:

```
AMdata_sc_corrected <- AMdata_sc_plain

AMdata_sc_corrected[] <-
```

```r
sapply(seq_along(AMdata_sc_plain), function(molecule_n){

  molecule_i <- AMdata_sc_plain[[molecule_n]]


  input_df <- data.frame(signal = molecule_i,
                         am_ratio = AMdata_sc$am_ratio) %>%
    mutate(ratio = signal / am_ratio,
           ratio = ratio / median(ratio[am_ratio == 1], na.rm = T),
           log_ratio = log(ratio, base = 10))

  if(dim(input_df%>%
       filter(is.finite(log_ratio) & am_ratio > 0.1 ))[1] < 10){

    molecule_i <- AMdata_sc_plain[[10]]


    input_df_main <- data.frame(signal = molecule_i,
                                am_ratio = AMdata_sc$am_ratio) %>%
      mutate(ratio = signal / am_ratio,
             ratio = ratio / median(ratio[am_ratio == 1], na.rm = T),
             log_ratio = log(ratio, base = 10))

    model <- quantreg::rq(
      formula = log(ratio, base = 10) ~ log(am_ratio, base = 10),
      tau = .5,
      data =   input_df_main %>%
        filter(is.finite(log_ratio) & am_ratio > 0.1 )
    )

    input_df$pred_ratio <- 10 ^ predict(model,
                                        newdata = input_df %>%
                                          mutate(am_sampling_ratio =
                                                   log(am_ratio, base = 10)))


    input_df$signal / input_df$pred_ratio


  } else {
    model <- quantreg::rq(
      formula = log(ratio, base = 10) ~ log(am_ratio, base = 10),
      tau = .5,
      data =   input_df %>%
        filter(is.finite(log_ratio) & am_ratio > 0.1 )
    )


    input_df$pred_ratio <- 10 ^ predict(model,
                                        newdata = input_df %>%
                                          mutate(am_sampling_ratio =
                                                   log(am_ratio, base = 10)))
```

```
      input_df$signal / input_df$pred_ratio

    }




  }, simplify = F)


####


norm_corrected <-
  cell_ion_intensity <-
  cell_normalization_Rappez_et_al(
    overlap_proportion_matrix = overlap_matrix,
    overlap_specificity_matrix = overlap_spec_matrix,
    AM_ion_intensity = AMdata_sc_corrected,
    AM_proportion_treshold_global = 0.3,
    skip_AM_zeros = FALSE                                ## keep zeros in AM vector
  )



cell_intensities_cor <- norm_corrected$cell_intensities[match(sc$ObjectNumber,
                                                              rownames(norm$cell_intensities)),]
```

Generate Seurat objects, make UMAPS

```
#### filter cells
include_cells <- apply(cell_intensities,1,function(i){mean(is.na(i))}) < .1

rownames(sc_fluorescence) <- rownames(cell_intensities_cor)

## HeLa-mCherry
## NIH3T3-GFP


scPlain <-
  CreateSeuratObject(
    counts = t(cell_intensities),
    assay = "metabolites",
    meta.data = data.frame(ratio = sc_fluorescence$ratio,
                           celltype = ifelse(sc_fluorescence$ratio < .8,"HeLa","NIH3T3"),
                           row.names = rownames(sc_fluorescence)
    ),
    project = "co-culture"
  )

scPlain$total_counts_per_cell <- colSums(scPlain@assays$metabolites@data)
scPlain$fraction_non_zero <- apply(scPlain@assays$metabolites@data, 2, function(cell){
  mean(cell != 0)
})
```

```r
#FeatureScatter(scPlain, "total_counts_per_cell", "fraction_non_zero", pt.size = 0.5)

selected_cells <- names(include_cells)[include_cells]
scPlain <- subset(scPlain, cells = selected_cells)

features <- rownames(scPlain)


## scaling
scPlain <- ScaleData(scPlain, features = features,
                     do.scale = TRUE,
                     do.center = TRUE)


## PCA
scPlain <- FindVariableFeatures(scPlain)
scPlain <- RunPCA(scPlain)

## elbow plot
#ElbowPlot(scPlain)

## umap
scPlain <- RunUMAP(scPlain, dims = 1:5, n.neighbors = 200, metric = 'cosine')

#Seurat::DimPlot(scPlain, group.by = 'celltype')


###

scCorrected <-
  CreateSeuratObject(
    counts = t(cell_intensities_cor),
    assay = "metabolites",
    meta.data = data.frame(ratio = sc_fluorescence$ratio,
                           celltype = ifelse(sc_fluorescence$ratio < .8,"HeLa","NIH3T3"),
                           row.names = rownames(sc_fluorescence)
    ),
    project = "co-culture"
  )

scCorrected$total_counts_per_cell <- colSums(scCorrected@assays$metabolites@data)
scCorrected$fraction_non_zero <- apply(scCorrected@assays$metabolites@data, 2, function(cell){
  mean(cell != 0)
})

#FeatureScatter(scCorrected, "total_counts_per_cell", "fraction_non_zero", pt.size = 0.5)

selected_cells <- names(include_cells)[include_cells]

scCorrected <- subset(scCorrected, cells = selected_cells)

features <- rownames(scPlain)
```

```r
## scaling
scCorrected <- ScaleData(scCorrected, features = features,
                         do.scale = TRUE,
                         do.center = TRUE)


## PCA
scCorrected <- FindVariableFeatures(scCorrected)
scCorrected <- RunPCA(scCorrected) #, features = features)

## elbow plot
#ElbowPlot(scCorrected)


## umap
scCorrected <- RunUMAP(scCorrected, dims = 1:4, n.neighbors = 200, metric = 'cosine')

#Seurat::DimPlot(scCorrected, group.by = 'celltype')
```

```r
Fig2E <-
  {
    list(
      data.frame(
        scPlain@reductions$umap@cell.embeddings,
        celltype = scPlain@meta.data$celltype,
        analysis = factor("none", levels = c("none","ISM (non-supervised)"))
      ),
      data.frame(
        scCorrected@reductions$umap@cell.embeddings,
        celltype = scCorrected@meta.data$celltype,
        analysis = factor("ISM (non-supervised)", levels = c("none","ISM (non-supervised)"))
      )) %>% bind_rows()

  } %>%
  ggplot(data = ., aes(x = UMAP_1, y = UMAP_2, color = celltype))+
  coord_fixed()+
  geom_point(size = .5, lwd=0)+
  guides(color = guide_legend(override.aes = list(alpha = 1, size = 3)))+
  theme_void()+
  labs(color = "", x = 'UMAP 1', y = 'UMAP 2')+
  scale_color_paletteer_d("ggthemes::calc") +
  facet_wrap(~analysis, ncol = 1)+
  theme(plot.background = element_rect(fill = "white", colour = "NA"),
        legend.justification = c(1,.5), strip.text = element_text(size = 12, face = "bold"),
        axis.title.y = element_text(angle = 90, size = 9),
        axis.title.x = element_text(size = 9))

Fig2E
```

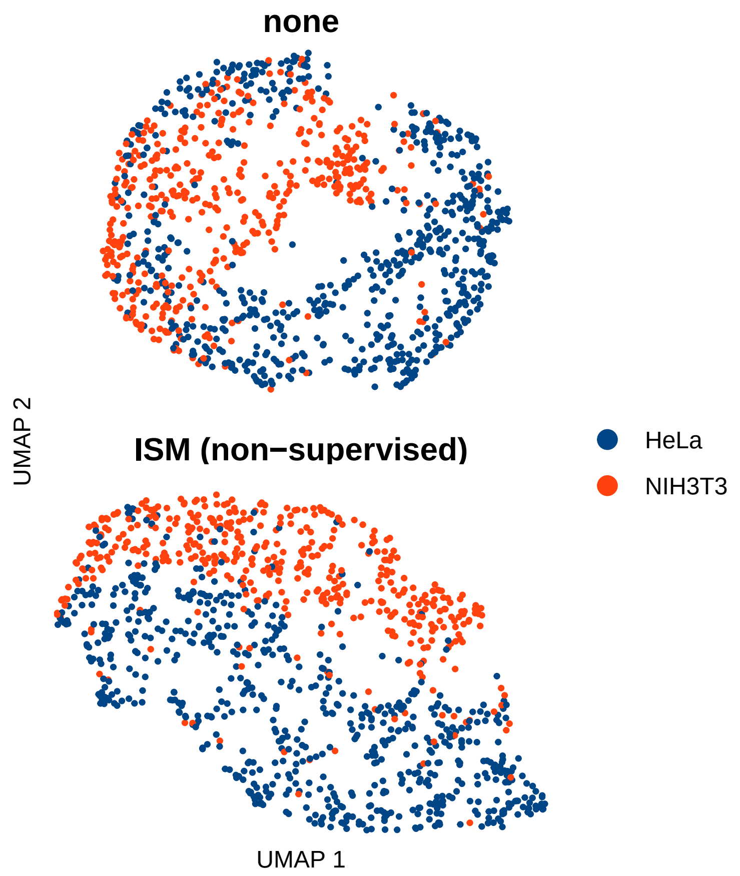


calculate intermixing

```
####

cell_intensities_cor_dist <-
  cell_intensities_cor[include_cells,] %>%
  scale %>%
  dist  %>%
  as.matrix

cell_intensities_cor_dist <-
  scCorrected@reductions$umap@cell.embeddings %>%
  scale %>%
  dist  %>%
  as.matrix


cell_intensities_cor_dist[upper.tri(cell_intensities_cor_dist, diag = TRUE)] <- NA

sc_fluorescence$celltype <- ifelse(sc_fluorescence$ratio < .8,"HeLa","NIH3T3")
sc_fluorescence$cellID <- sc$ObjectNumber



cell_intensities_dist <-
  scPlain@reductions$umap@cell.embeddings %>%
  scale %>%
```

```
  dist  %>%
  as.matrix



cell_intensities_dist_intermixing <-
  sapply(seq_along(cell_intensities_dist[1,]), function(row_n){
    parent <- rownames(cell_intensities_dist)[row_n]
    parent <- sc_fluorescence$celltype[match(parent, sc_fluorescence$cellID)]

    nn <- names(sort(cell_intensities_dist[row_n,-row_n])[1:10])
    nn <- sc_fluorescence$celltype[match(nn, sc_fluorescence$cellID)]
    data.frame(parent = parent, frac_same = mean(nn %in% parent))
  }, simplify = F) %>% bind_rows()

cell_intensities_dist_intermixing %>% group_by(parent) %>%
  summarise(frac_same = mean(frac_same))
```

```
## # A tibble: 2 x 2
##   parent frac_same
##   <chr>      <dbl>
## 1 HeLa       0.824
## 2 NIH3T3     0.728
```

```
cell_intensities_cor_dist <-
  scCorrected@reductions$umap@cell.embeddings %>%
  scale %>%
  dist  %>%
  as.matrix


cell_intensities_dist_cor_intermixing <-
  sapply(seq_along(cell_intensities_cor_dist[1,]), function(row_n){

    parent <- rownames(cell_intensities_cor_dist)[row_n]
    parent <- sc_fluorescence$celltype[match(parent, sc_fluorescence$cellID)]


    nn <- names(sort(cell_intensities_cor_dist[row_n,-row_n])[1:10])
    nn <- sc_fluorescence$celltype[match(nn, sc_fluorescence$cellID)]
    data.frame(parent = parent, frac_same = mean(nn %in% parent))
  }, simplify = F) %>% bind_rows()




Fig2F <-
  rbind(cell_intensities_dist_cor_intermixing %>% mutate(comparison = "ISM"),
        cell_intensities_dist_intermixing %>% mutate(comparison = "none")) %>%
  mutate(comparison = factor(comparison, levels = c("none","ISM"))) %>%
  ggplot(.,
         aes(x = comparison, y = 1-frac_same, fill = comparison))+
  stat_summary(fun = mean, fun.min = median, fun.max = median,
               geom = "bar",  show.legend = F,
```

```
               width = 0.7, lwd = 0.2)+
  stat_summary(fun.data = mean_se, geom = "errorbar", width = .2)+
  scale_fill_manual(values = palettes_d$ggthemes$calc[c(1,4)])+
  coord_flip()+
  scale_x_discrete(limits=rev)+
  labs(y =
         expression("intermixing "*
                      italic('(fraction of 10 nearest neighbors from the other cell type)')),
       x = "")+
  theme_minimal()+
  theme(axis.text.x = element_text(size = 9),
        axis.title = element_text(size = 11), panel.grid = element_blank(),
        axis.ticks = element_line(),
        panel.background = element_rect(colour = "black", size = 1),
        plot.background = element_rect(color = NA, fill = "white"))

Fig2F
```

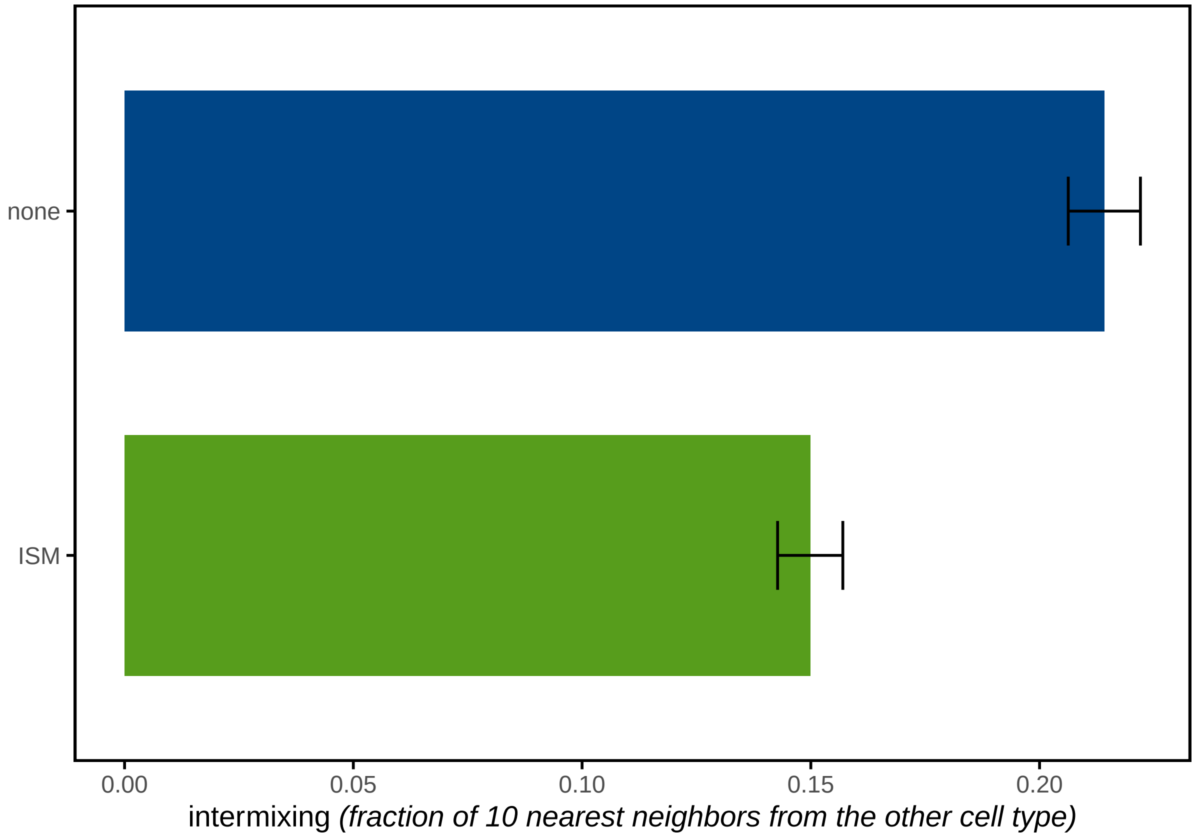
